# Tau modulates nuclear integrity, chromatin, and cholesterol synthesis genes via the nuclear envelope

**DOI:** 10.64898/2025.12.05.692577

**Authors:** Lisa Diez, Renata Ponce-Lina, Cagla Kilic, Rithika Sankar, Frieda Balduin, Cong Wang, Pia Grundschoettel, Koki Sakurai, Bilge Askin, Chiara Bordier, Tejaswini Karra, Katharina Tipp, Boglarka Krisztina Nagy-Herczeg, Maryam Mohamaddokht, Leandre Ravatt, Satabdee Mohapatra, Alvaro Dominguez-Baquero, Ramesh Adakkattil, Maximilian Franck, Sabrina Hübschmann, Elena De Domenico, Marc Beyer, Tomohisa Toda, Helena Radbruch, Tara L. Spires-Jones, Alexander von Appen, Thomas Ulas, Andrew Newman, Fan Liu, Susanne Wegmann

## Abstract

The intracellular re-distribution of the neuronal microtubule-associated protein Tau, from the axon into the somatodendritic compartment, is a physiological stress-related event and occurs early in Alzheimer’s disease (AD). Nuclear envelope distortions have been linked to the presence and aggregation of pathological Tau near the nucleus in these diseases. How physiologically increased soma Tau levels, enabling Tau interactions with the nucleus, impact nuclear integrity and neuronal physiology is unclear. Combining proximity biotinylation interactomics with chromatin imaging and molecular assays, we show that soluble Tau interacts with proteins coordinating chromatin at the inner nuclear membrane, including lamin B receptor and SUN1. This interaction promotes nuclear envelope invaginations and damage and changes the coordination of DNA at the nuclear lamina. Increasing somatodendritic Tau is sufficient to upregulate the expression of multiple transcription factors implicated in AD pathogenesis and to reduce expression of genes involved in cholesterol biosynthesis, which seem coordinated at lamin associated domains. These nuclear envelope-related mechanisms suggest that physiological, neuronal stress-related somatodendritic Tau missorting can initiate chromatin-related cascades important for early changes in AD and tauopathies.

## Introduction

Tau, encoded by the *MAPT* gene, is an intrinsically disordered protein that has a high affinity to microtubules (MTs) (Mandelkow *et al*, 1995; Jeganathan *et al*, 2008). The majority of Tau is expressed in neurons of the central nervous system. In mature neurons, Tau is enriched in the axon where its microtubule association is regulated by phosphorylation and other post-translational modifications (PTMs) (Parra Bravo *et al*, 2024). In conditions where Tau gets “activated”, the subcellular distribution of phosphorylated Tau – detached from MTs – changes, and similar levels of soluble cytosolic Tau can be found throughout axon, soma, and dendrites. This subcellular phospho-Tau redistribution (= missorting) can be triggered by various physiological neuronal stress conditions, including temperature changes, mechanical stimulation, and neuronal hyperactivity (Sultan *et al*, 2011; Schneeweis *et al*, 2024; Braun *et al*, 2020), and is considered an early pathological change in Tau-related brain disorders like frontotemporal dementia (FTD) and Alzheimer’s disease (AD).

Pathological accumulation of phosphorylated Tau in neuronal cell bodies can be observed in a number (>20) of neurological conditions, e.g., in traumatic brain injury, epilepsy, autoimmune brain diseases, and – most prominently – AD and primary tauopathies (Spillantini & Goedert, 2013; Chang *et al*, 2021). It is widely believed that the accumulation of soluble phospho-Tau precedes Tau aggregation in neuronal cell bodies, which is a pathological hallmark and correlate of neuronal loss in these diseases. The actual, neurophysiological reason and function of somatodendritic Tau “missorting” remains unclear. It is likely that the interaction of Tau with molecules and organelles in the somatodendritic compartment, like the nucleus, builds the foundation for stress-related Tau functions. Notably, these interactions of Tau would be enabled in both physiological stress and disease conditions.

In human AD and FTD brains, as well as in cell models of tauopathy, Tau was shown to cluster at the outer nuclear envelope (NE) and to impair nuclear transport through nuclear pores (Jiang & Wolozin, 2021; Kang *et al*, 2021; Eftekharzadeh *et al*, 2018; Diez *et al*, 2022). This could affect chromatin organization and gene expression. Accordingly, changes in chromatin and gene expression occur in AD and FTD brains and in brains of tauopathy animal models (Rico *et al*, 2018; Frost *et al*, 2014; Klein *et al*, 2019). Global chromatin relaxation (Frost *et al*, 2014), neurotoxic re-activation of retroviral transposable elements (Ramirez *et al*, 2022; Guo *et al*, 2018), and activation of senescence-like expression profiles (Musi *et al*, 2018) have been reported to correlate with Tau pathology.

In the physiological context, Tau inside the nucleus was shown to interact with DNA, histone proteins, and the nucleolus (Maina *et al*, 2018; Sultan *et al*, 2011; Liu & Götz, 2013; Benhelli-Mokrani *et al*, 2018), with suggested functions in gene expression, DNA protection and compaction, and nucleolar stress response. However, there is currently no consensus about the impact of Tau on chromatin, and reported effects range from Tau-induced global chromatin relaxation and transcriptional activation (Klein *et al*, 2019; Frost *et al*, 2014) to stabilization of chromatin compaction and transcriptional repression (Benhelli-Mokrani *et al*, 2018; Rico *et al*, 2018). These inconclusive reports may – at least in part – result from difficulties to distinguish between physiological nuclear actions of soluble somatodendritic Tau *versus* pathological actions or loss of nuclear functions of oligomeric and aggregated Tau.

In this study, we aimed at understanding whether Tau missorting – i.e., the increase of Tau in the somatodendritic compartment – is sufficient to induce nuclear distortions and changes in neuronal chromatin and gene expression, and which Tau interactions underlie these effects. The herein newly identified interactions of Tau with nuclear envelope proteins, like the lamin B receptor (LBR) and SUN1, provide a molecular mechanism for neuronal chromatin and gene expression alterations in response to physiological and disease-associated soluble Tau missorting.

## Results

### Pathological Tau is associated the neuronal nuclear envelope distortions in the brain

In human Alzheimer’s disease (AD) brains, diffuse misfolded (Alz50+) and aggregated Tau accumulate in the soma of neurons (Figure 1A). In neurons having soluble phosphorylated Tau (pS202/pS205/pT231), presumably prior to developing Tau aggregation pathology, p-Tau can be found at the nuclear envelope (NE; labeled with anti-SUN1 antibody; Figure 1B). Neurons with larger p-Tau aggregates in the cell body did not show this Tau pattern (Figure 1B), indicating that cytosolic phosphorylated Tau may have a special affinity to the NE. This is in line with previous observations of our own and others, for example, that Tau clusters forming during the aggregation process associated with the NE (Kang *et al*, 2021; Hochmair *et al*, 2022; Eftekharzadeh *et al*, 2018) and that Tau interactions with nuclear pore proteins (Nups) affect neuronal nuclear transport (Eftekharzadeh *et al*, 2018).

**Figure 1.**
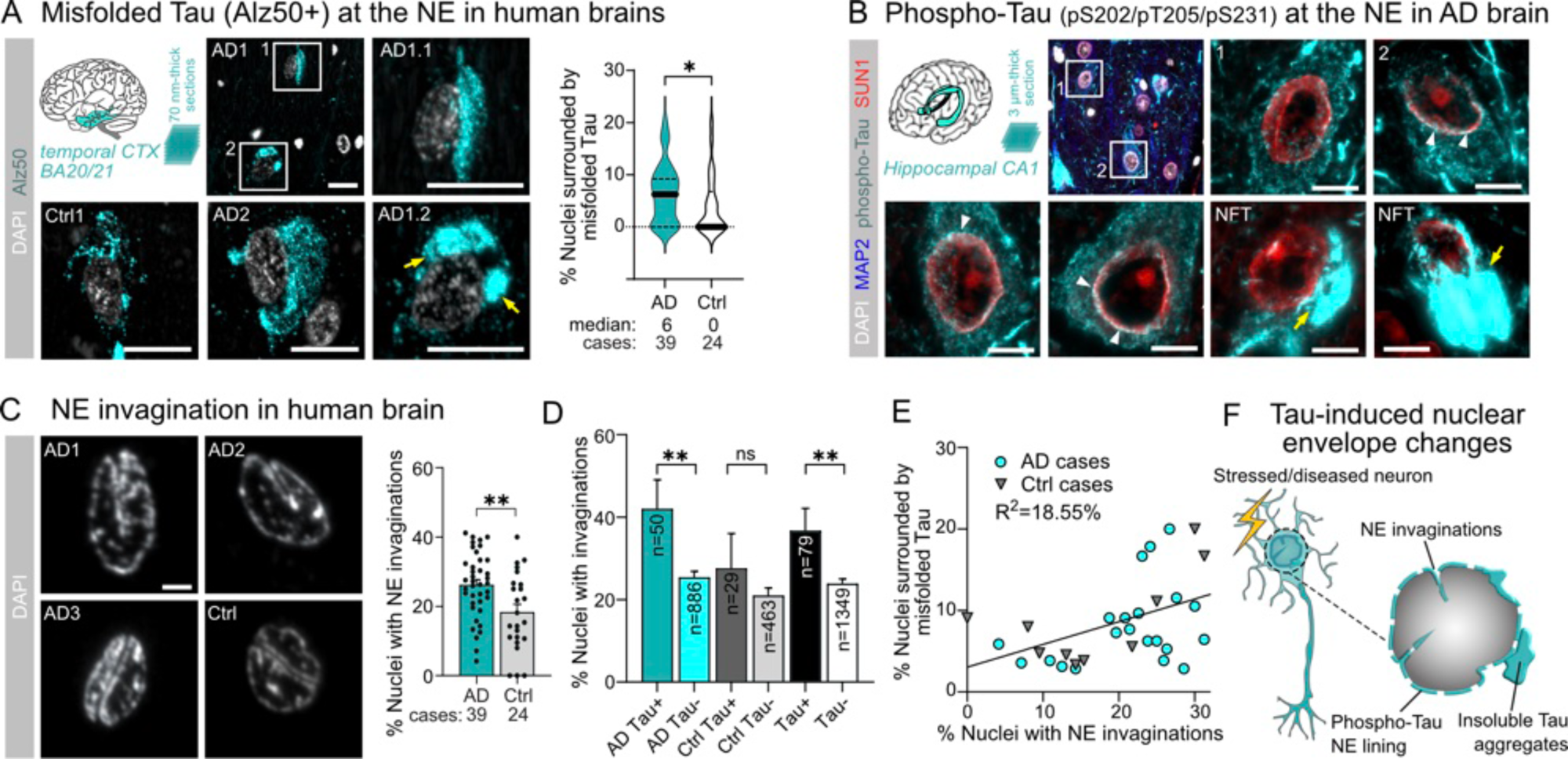
Tau accumulation and nuclear envelope distortion in Alzheimer’s disease (AD) brains. (A) Misfolded Tau (Alz50+) near the nucleus in AD (39 cases) and control (24 cases) brain sections (70 nm-thick) of temporal cortex (BA20/21). Data shown as median±quartiles. Student’s t-test with Mann-Whithey post-test. Scale bars = 10 µm. (B) Representative images of immunolabeled nuclear envelope (NE) protein SUN1 and phospho-Tau (pS202/pT205/pT231) in neurons of AD brains (hippocampal CA1). Neurons with diffuse granular phospho-Tau staining in the soma show phospho-Tau “foci” in proximity to SUN1 in the NE (white arrowheads). Tangles (NFTs) do not show this Tau arrangement. Scale bars = 5 µm. (C) AD and control brain nuclei showing nuclear membrane folds (“invaginations”). Data shown as mean±SEM, one data point per case. Student’s t-test with Tukey post-test. Scale bars = 2 µm. (D) Percent of neuronal nuclei having invaginations (nuclear membrane folds) in relation to the presence of misfolded/aggregated Tau near the nucleus in AD and control brains (related to Figure 1C). Data shown as mean±SD, n = number of nuclei as indicated. Two-tailed, unpaired Student’s t-test comparing with and without Tau at nucleus within AD and Ctrl groups. (E) Correlation between the percent of nuclei having nuclear invaginations and phospho-Tau near the nucleus in AD and control brain sections. Data points present individual cases, linear regression indicated. (F) Illustration of nuclear alterations observed in AD brains. Neurons with Tau in the soma tend to have more NE invaginations. Small phospho-Tau “clusters” appear around the NE and insoluble Tau aggregates next to the nucleus.

In addition, AD (Braak 5/6) compared to control (Braak 0/1) brains had more nuclei with NE deformations – folds and invaginations –, whereby the occurrence of invagination was higher in neurons with misfolded Tau in the soma (Figure 1C,D; Supplemental Video V1-4). In both AD and control brains, the number of nuclei surrounded by aggregated Tau correlated with the number of neuronal nuclei having invaginations (Figure 1E). These data suggest that the accumulation of non-aggregated phosphorylated Tau in the neuronal soma may be associated with nuclear envelope distortions (Figure 1F). The details of Tau interactions with the NE, and their consequences, remain unclear though.

### Increased somatodendritic soluble Tau is sufficient to induce nuclear shape changes

Tau protein levels - in healthy neurons high in axons and low in soma and dendrites - increase in the soma in physiological and pathological neuronal stress conditions. To investigate whether the somatodendritic missorting of soluble Tau is sufficient, or whether pathological misfolding would be necessary, to induce NE alterations, we modulated soma Tau levels in cultured hippocampal neurons using adeno-associated virus (AAV) constructs. A reduction in Tau levels were achieved by inhibiting endogenous Tau transcription (Tau KD) through expression of a *MAPT*-targeted zinc-finger protein array coupled to a gene-silencing KREB transcription factor, ZFP90 (Wegmann *et al*, 2021), which reduced Tau levels by >50% compared to untreated neurons (Tau base; Figure 2A, Figure EV1A). The Tau knockdown AAV construct expressed equal amounts of ZFP90 and GFP as a transduction marker, separated by a self-cleaving 2a peptide. A control AAV expressing target-less zinc finger array (ZFP72) did not change Tau levels compared to GFP expressing or non-treated neurons (Figure EV1B). To produce elevated somatodendritic Tau levels, we expressed GFP-labeled full-length wildtype Tau (2N4R isoform; Tau OE) or FTD-mutants Tau^P301L^ (TauPL OE) and Tau^P301L/S320F^ (TauPL/SF OE; Figure 2A). GFP-Tau^P301L^ does not aggregate in neurons, yet has a higher oligomerization propensity than wildtype Tau due to increased accessibility of an aggregation-prone motif (PHF6*; amino acids *^306^VQIVYK^311^*) in the beginning of R3 in the Tau repeat domain (Von Bergen *et al*, 2000). GFP-Tau^P301L/S320F^, having an additional pro-aggregation mutation in R3, spontaneously forms NFT-like aggregates in the soma of a subset (5-20%) of neurons (Jorge-Oliva *et al*, 2023; Hochmair *et al*, 2025). Here we used these FTD-Tau variants to test the effect of Tau oligomerization and aggregation on the NE. After 7 days of AAV expression, Tau KD neurons had <50% of total endogenous Tau left (by Western blot; Figure 2B). Tau OE and TauPL OE neurons had a 5-7-fold increase in total soluble Tau. TauPL/SF OE neurons showed only a 2-fold increase in soluble Tau (RIPA buffer extracts), which may be linked to the sequestering of soluble Tau into detergent-insoluble NFTs.

**Figure 2.**
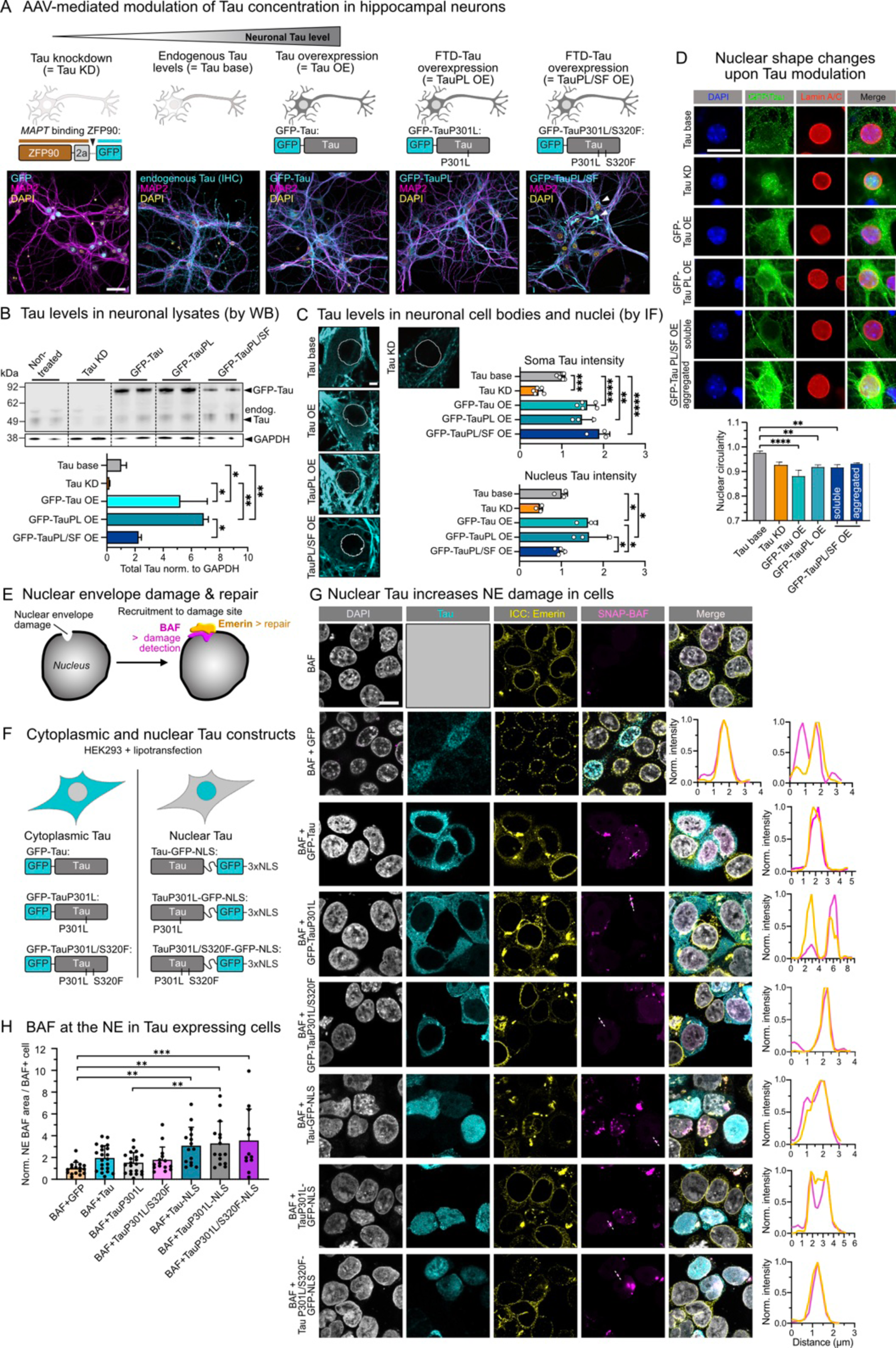
Soluble Tau in the soma increases nuclear Tau and induces nuclear envelope damage. (A) Modulating Tau levels in primary mouse hippocampal neurons using adeno-associated viruses (AAVs). Overexpression of GFP-tagged human wild-type Tau (2N4R isoform) (Tau OE), FTD-mutant Tau^P301L^ (TauPL OE) or FTD-mutant Tau^P301L/S320F^ (TauPL/SF OE) increases Tau levels compared to endogenous mouse Tau (Tau base), whereas the Tau (*MAPT*) gene silencing zinc-finger protein transcription factor (Wegmann *et al*, 2021) (ZFP90) decreases endogenous Tau levels (Tau KD). Note that neurons with AAV ZFP90-2a-GFP express GFP and ZFP90 as separate proteins due to the 2a self-cleavage site. Images show expressed GFP, immunolabeled MAP2, and DAPI in AAV transduced neurons (DIV12). In untreated neurons (Tau base), endogenous Tau was immunostained as well. Scale bars = 50 µm. (B) Western blot of neuronal lysates (RIPA buffer extracts) showing, compared to Tau base neurons, increased soluble Tau levels in Tau OE, TauPL OE, and TauPL/SF OE neurons and reduced Tau levels in Tau KD neurons. Glia were removed from cultures by AraC treatment. data shown as mean±SD, n = 2 experiments. One-way ANOVA with Dunnett post-test. (C) Quantification of Tau levels in neuronal somata and nuclei based on Tau immunofluorescence intensity. Positions of nuclei, determined from DAPI channel (not shown), are outlined in white. data shown as mean±SD, n = 3 images and 50-100 neurons per condition. One-way ANOVA with Dunnett post-test. Scale bars = 5 µm. (D) Nuclear circularity determined based on Lamin A/C signal in hippocampal neurons with different Tau levels (Tau variant overexpression or Tau KD) compared to Tau base neurons. Representative images of Tau base (Tau immunostaining), Tau OE, TauPL OE, TauPL/SF OE, and Tau KD neurons. Nuclear circularity was quantified using Lamin A/C signal to define the nuclear boundary. n = 20-48 neurons per condition, data shown as mean±SD, one-way ANOVA with Tukey post-test. Scale bar = 1 µm. (E) Schematic model of NE rupture and repair, showing local accumulation of BAF at sites of NE damage and subsequent recruitment of Emerin during the repair process. (F) Tau constructs to test effect of (mostly) cytoplasmic or nuclear Tau on NE damage. Cytoplasmic: GFP-Tau, GFP-Tau^P301L^ and GFP-Tau^P301L/S320F^. Nuclear (NLS): Tau-GFP–NLS, Tau^P301L^-GFP-NLS, and Tau^P301L/S320F^-GFP-NLS. Negative controls were untransfected or expressed GFP. (G) Representative images of HEK293 cells co-expressing the GFP-labeled Tau constructs and SNAP-BAF. BAF accumulates at NE damage sites. Fluorescence intensity line plots along white dashed lines in BAF channel show colocalization of BAF and Emerin, indicating initiated NE repair processes in all conditions. Scale bars = 10 µm. (H) Quantification of SNAP-BAF accumulation at the NE of HEK293 cells co-expressing GFP (control) or Tau constructs (see panel G) and SNAP-BAF. For each condition, the SNAP-BAF signal area at the NE was calculated for BAF+ cells. Data shown as mean±SD, normalized to controls (SNAP-BAF+GFP). n=15-20 cells per condition from 3 experiments. One-way ANOVA with Tukey post-test.

It was previously suggested that endogenous Tau can be found inside the nucleus, particularly in conditions of stress-induced somatodendritic missorting (Violet *et al*, 2014; Benhelli-Mokrani *et al*, 2018). To test how AAV-modulated Tau levels correlated with nuclear Tau levels, we determined total Tau immunofluorescence (endogenous + GFP-Tau) in the soma and nucleus of AAV-treated neuronal cultures (Figure 2C). We found that both somatic and nuclear Tau levels were reduced to ∼30% in Tau KD neurons (Figure 2C). In Tau OE and TauPL OE neurons, somatic and nuclear Tau levels were ∼1.5-fold increased compared to untreated controls (Tau base). Notably, in hippocampal neurons of AD brains, we measured an up to ∼2.5-fold increased levels of diffuse, non-aggregated Tau in the soma (Figure EV1C). Interestingly, TauPL/SF OE neurons had ∼2-fold increased Tau in the soma but nuclear Tau levels were comparable to untreated neurons. Together, these data suggest that the levels of somatic and nuclear Tau are correlated in the case of soluble Tau variants, while pro-aggregant TauPL/SF cannot enter the nucleus and resides in the cytoplasm.

Finally, to test the effect of Tau levels on nuclear shape, we measured the circularity of neuronal nuclei after immunostaining for Lamin A/C. Expression of all Tau variants led to a decrease in nuclear circularity compared to control neurons, indicating an effect of elevated Tau levels on NE integrity (Figure 2D), whereby overexpression of wildtype Tau had a stronger effect than FTD-mutant TauPL and TauPL/SF expression. Neurons with TauPL/SF aggregates in the soma showed little effects on nuclear circularity (non-significant), as did Tau KD neurons. These data support the idea that elevated levels of soluble, non-aggregated Tau in the soma – and nucleus - are sufficient to induce nuclear shape changes and envelope distortions in neurons.

### Increased levels of Tau in the nucleoplasm induce nuclear envelope damage

Next, we tested whether increased soluble Tau levels were sufficient to induce NE rupture, and whether this was related to the presence of Tau outside (in cytoplasm) or inside (in nucleoplasm) the nucleus. NE rupture can be detected by local accumulation of BAF (barrier-to-autointegration factor) at NE damage sites, and the subsequent recruitment of Emerin to these sites during repair (Halfmann *et al*, 2019) (Figure 2E). We expressed SNAP-BAF together with different GFP-labeled Tau and Tau-NLS (nuclear localization signal) variants in human cells for 24 h (Figure 2F) and determined the amount of SNAP-BAF accumulation at the NE, as well as its colocalization with Emerin (by immunostaining). In all cells expressing GFP (control), Tau, or Tau-NLS, Emerin was recruited from the NE into BAF accumulations (Figure 2G), indicating the successful initiation of NE repair processes. Quantification of BAF accumulation at the NE showed that all Tau variants carrying a recombinant NLS had significantly more NE ruptures than GFP-expressing control cells (Figure 2H). Tau variants without NLS, primarily expressed in the cytoplasm, showed only a small, non-significant increase in NE BAF accumulation, i.e., NE ruptures. These data indicate that soluble Tau inside the nucleus promotes NE damage but does not impact initial steps (Emerin recruitment) of the repair process.

### Proximity biotinylation suggests Tau interactions with inner nuclear envelope

To understand how soluble Tau can induce NE distortion and damage, we decided to identify nuclear interaction partners of Tau by proximity biotinylation. We fused full-length human wildtype (TurboID-Tau) or FTD-mutant Tau^P301L^ (TurboID-Tau^P301L^; Figure 3A,B) to the biotin ligase TurboID (Branon *et al*, 2018), which facilitated rapid (within minutes upon addition of biotin) biotinylation of proteins in proximity (∼35 nm distance) to Tau or Tau^P301L^ in their native cellular environment. We transiently expressed TurboID-Tau and TurboID-Tau^P301L^ in human neuroblastoma cells (SH-SY5Y line), lysed the cells in hypotonic buffer, and concentrated their nuclei by centrifugation in order to enrich nuclear Tau interaction partners. Biotinylated nuclear proteins that have been in proximity to Tau or Tau^P301L^ were pulled-down from the nuclear pellet using magnetic streptavidin-coated beads and analyzed by mass spectrometry (Figure 3B,C,D; Figure EV2A,B). Expression of TurboID alone was used to obtain a control interactome for unspecific TurboID activity. Notably, while the TurboID-Tau fusion constructs (MW ∼85 kDa) were mostly located in the cytoplasm, the TurboID only control (MW ∼39 kDa) was abundant in the nucleus (Figure 3C), rendering it a stringent control for identified potential nuclear Tau interactors. In addition, as positive control and reference interactome for protein-specific nuclear interactions, we also determined the nuclear proximity biotinylation proteome for TurboID fused to the C-terminal fragment of TDP-43 (TurboID-TDP-43ctf; Figure EV2C-F), a component of insoluble cytoplasmic TDP-43 aggregates in ALS/FTD brains (Igaz *et al*, 2008). A proximity biotinylation interactome of TDP-43ctf has previously been determined in mouse neuroblastoma cells (Chou *et al*, 2018).

**Figure 3.**
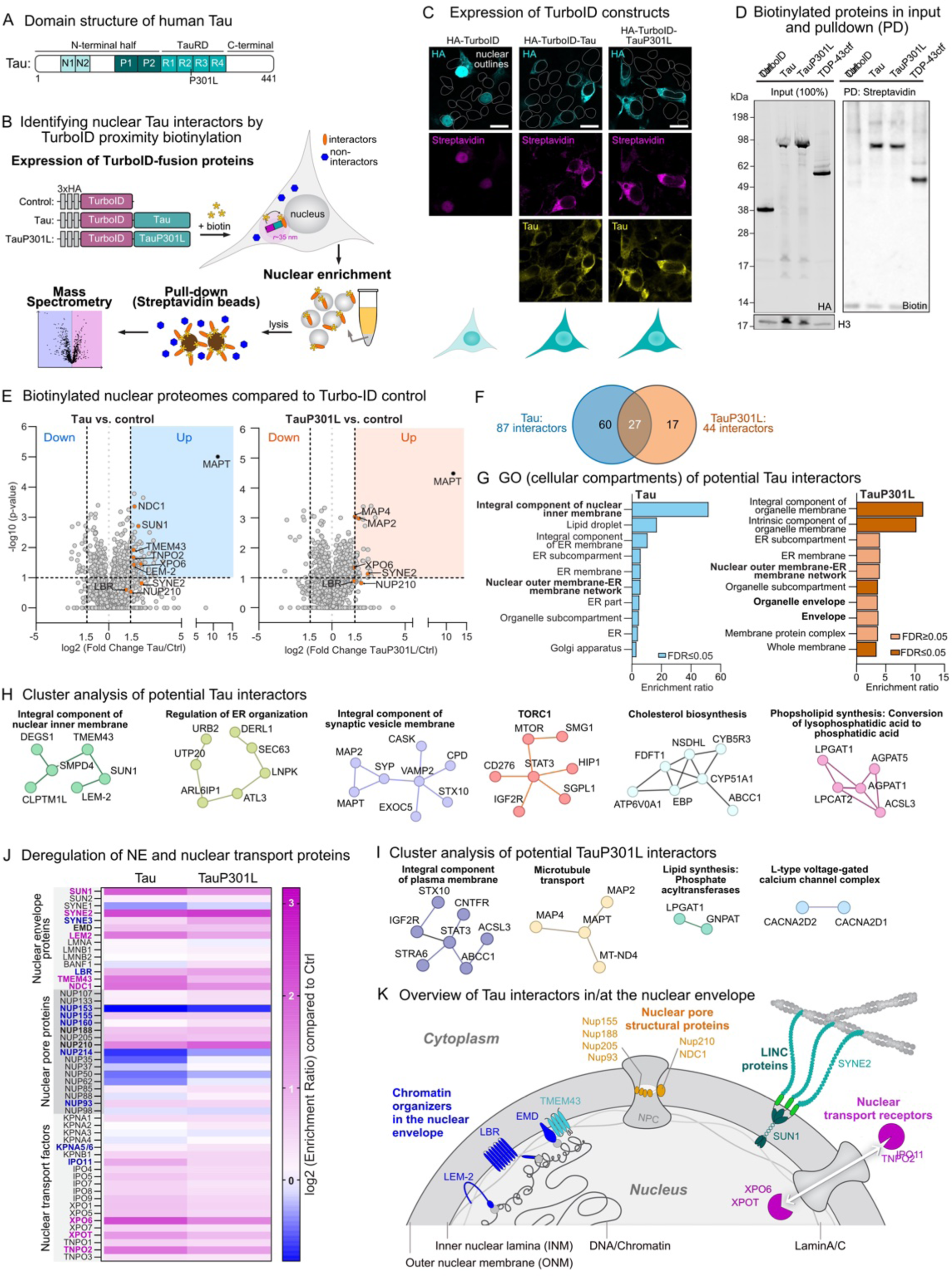
Nuclear Tau proximity labeling identifies nuclear envelope and nuclear transport proteins. (A) Domain structures of human Tau (2N4R isoform) with indicated position of the P301L FTD-mutation in repeat R2. Tau domain structure shows the position of the N-terminal inserts, N1 and N2, and proline-rich regions P1 and P2 in the N-terminal projection domain, the 4 pseudo-repeats R1-R4 in the microtubule-binding domain. (B) Workflow of nucleus-specific proximity biotinylation and identification of proteins associated with HA-TurboID-Tau, HA-TurboID-Tau^P301L^, and HA-TurboID control. TurboID constructs are expressed in SH-SY5Y cells, biotinylation of interacting proteins enabled by addition of biotin to the culture medium. Nuclear biotinylated proteins are enriched through streptavidin pulldown from nuclear enriched fractions, followed by MS analysis. (C) Expression of TurboID constructs in SH-SY5Y neuroblastoma cells. Anti-HA shows expression and cellular localization of HA-TurboID constructs, successful biotinylation is verified by anti-Streptavidin staining. Tau immunostaining detects both endogenous and recombinant Tau. Scale bars = 10 µm. (D) Example western blot for streptavidin pull-down experiments. Anti-HA antibody detects TurboID constructs in nuclear lysates (input). Biotinylated proteins in streptavidin bead pull-down factions are detected with using anti-Biotin antibody. Histone-3 (H3) was used as loading control for nuclear fractions/inputs. Nuclear lysates from cells expressing TurboID fused to the C-terminal fragment of TDP-43 (TurboID-TDP-43ctf) were included as positive control. (E) Volcano plots of proximity-biotinylated proteins in TurboID-Tau (left) and TurboID-Tau^P301L^ (right) each compared to TurboID only control. Significant differences between experimental groups and control were identified using a two-sided t-test. A log2[fold change] >1.5 and p-value <-log10[1] were used as threshold for significance. Selected interactors are indicated as orange data points and labeled with gene/protein identities. Data retrieved from 5 biological replicates. (F) Venn diagram showing overlap of significant interactors between Tau and Tau^P301L^. (G) Pathway enrichment analysis (Gene ontology (GO), Cellular compartment) for Tau (left) and Tau^P301L^ (right). (H) STRING analysis of potential Tau interactors (log2[fold change] >1.5 and p-value <-log10[1]) identified several functional clusters, corresponding to related biological modules (e.g., Integral component of nuclear inner membrane, regulation of endoplasmic reticulum (ER) organization, integral component of synaptic vesicle membrane, target of rapamycin complex 1 (TORC1), cholesterol biosynthesis, and phospholipid synthesis). (I) STRING analysis of potential Tau^P301L^ interactors (log2[fold change] >1.5 and p-value <-log10[1]) identified several functional clusters, corresponding to related biological modules (e.g., Integral component of plasma membrane, microtubule transport, lipid synthesis, and L-type voltage-gated calcium channel complex). (J) Heat map representing deregulation of selected nuclear envelope (NE) and nuclear transport proteins relative to TurboID control. (K) Scheme showing identified potential Tau and TauP301L interactors in and at the nuclear envelope. Identified proteins can be categorized as (i) chromatin organizers in the NE (LEM2, LBR, EMD), (ii) structural membrane and nuclear pore complex (NPC) proteins (Nup210, NDC1, Nup155, Nup188, Nup205, Nup93), (iii) LINC complex proteins (SUN1, SYNE2), and (iv) nuclear transport receptors (XPO6, XPOT, TNPO2, IPO11).

Proteome analysis of biotinylated proteins from five independent cultures revealed multiple significantly enriched proteins in the TurboID-Tau (87 proteins) and TurboID-Tau^P301L^ (44 proteins) proximity proteome compared to TurboID alone (log2[fold change Tau/Ctrl] >1.5, p-value <10^-1^) (Figure 3E; Supplemental Tables T1, T2). As expected for bait proteins in our approach, Tau (MAPT) itself was maximally enriched in the Tau and Tau^P301L^ biotinylation proteomes (Figure 3E). Among the significantly enriched proteins, we found 27 proteins that were enriched in the biotinylation proteomes of both Tau variants (∼30% of potential TurboID-Tau interactors (27/87), ∼60% of TurboID-Tau^P301L^ interactors (27/44); Figure 3F). Interestingly, the TurboID-Tau proteome showed more overlap with the proteome of TurboID-TDP-43ctf ((37/87) = 43% potential Tau interactors), a protein that shuttles between nucleo-and cytoplasm (Figure EV2G; Supplemental Table T3).

Gene ontology (GO) analysis for biotinylated proteins enriched in TurboID-Tau and TurboID-Tau^P301L^ over TurboID control proteomes clustered potential Tau interactors into the cellular components *nuclear inner membrane*, *ER membrane*, and *nuclear outer membrane* (Figure 3G). Tau^P301L^ had overall fewer potential interactors, which clustered in the cellular components *nuclear outer membrane* and *organelle envelope*. Cluster analysis of significantly enriched proteins (log2[fold change] > 1.5 and-log10[p-value] > 1) using STRING (Szklarczyk *et al*, 2023) grouped potential Tau interactors in the inner nuclear and in the synaptic vesicle membrane, and suggested their involvement in ER organization, TORC1 complex, and cholesterol and phospholipid biosynthesis (Figure 3H). Cholesterol synthesis and lipid metabolism were also the strongest enriched categories in GO Biological processes for Tau interactors. hits for potential Tau interactors (FigureEV2H, Supplemental Table T4). For TauPL, potential interactors, clustered, for example, as component of the plasma membrane, were involved in microtubule transport and lipid synthesis, i.e., phosphate acyltransferases, or were calcium channel subunits (Figure 3I). Potential interactors of our positive control, TDP-43ctf, clustered in categories associated with the *nuclear outer membrane* and *nuclear envelope* (Figure EV2H), confirming previous results (Chou *et al*, 2018). In summary, GO and cluster analyses suggested that many of the nuclear Tau interactors are involved in NE and ER associated processes, i.e., chromatin organization at the inner nuclear membrane and metabolic regulation including cholesterol and lipid synthesis. Notably, since the NE is an extension of the endoplasmic reticulum (ER), interactions of Tau with components of NE and ER could be functionally and/or structurally related, and could both contribute to nuclear shape changes and NE damage induced by soluble Tau.

Tau interactions with the NE may explain how elevated soluble Tau levels could directly induce NE dysfunction and distortion. We find that both Tau and Tau^P301L^ may interact with NE proteins involved in chromatin organization at the inner nuclear lamina as well as with several proteins involved in nuclear transport (Nups and nuclear transport receptors, NTRs; Figure 3J). Thereby, especially NE proteins seem to be more enriched among potential Tau than TauPL interactors. Identified NE-specific proteins could be grouped into four protein categories (Figure 3K): (i) INM embedded proteins (LBR (= lamin B receptor), SUN1, and LEM-domain proteins, LEM-2 and EMD (= Emerin)) that coordinate chromatin association and repression at the inner leaflet of the NE (Zuleger *et al*, 2011), as well as lamin associated inner nuclear membrane protein TMEM43, important for NE structure and anchoring Emerin in the envelope (Bengtsson & Otto, 2008). (ii) Structural nuclear pore complex (NPC) proteins (Nup210, NDC1) in the nuclear envelope-embedded ring of the NPC (Schmidt & Görlich, 2016). (iii) Components of the LINC (linker of nucleoskeleton and cytoskeleton) complex (SYNE2 (=Nesprin-2), SUN1) required to translate cytoskeletal mechanical forces into changes in chromatin structure and gene expression (Sosa *et al*, 2013; Rothballer & Kutay, 2013); (iv) Nuclear transport receptors (e.g., TNPO2 and XPO6) that regulate the nuclear import and export, respectively, of biomolecular cargo through NPCs (Terry *et al*, 2007). Notably, we did not identify potential interactions of Tau with FG-rich Nups, i.e., Nup98 or Nup62, which we had previously reported to interact with Tau (Eftekharzadeh *et al*, 2018; Diez *et al*, 2022). One may account this to the rather short (10 min) biotinylation reaction time and the potential inaccessibility of Nups in the dense FG-Nup phase of the pore (Wente & Rout, 2010). Interestingly, chromatin associated proteins of the inner NE leaflet (LBR, SUN1, LEM-2, and EMD) were not identified as significant interactors of Tau^P301L^, indicating that oligomerization-prone FTD-mutant Tau may engage less in intranuclear interactions. This supports our observations that Tau^P301L/S320F^ barely enters the nucleus at high cytosolic levels (Figure 2C) and that soluble wildtype Tau – entering the nucleus - is most effective in inducing NE distortion (Figure 2D). LBR, however, was previously found in small cytoplasmic FTD-Tau aggregates (Jiang *et al*, 2021).

### Tau interacts with LBR and SUN1 at the inner nuclear envelope

The detection of multiple chromatin binding envelope proteins as potential interactors of Tau was a surprising finding. To verify these interactions, we performed co-immunoprecipitation (co-IP) of endogenous Tau (client) with different NE proteins (bait proteins) - LBR, Nesprin-2, and Lamin A/C - from nuclear enriched fractions of SH-SY5Y cells (Figure 4A, Figure EV3A). Intensity measurement indicated a mild interaction of Tau with LBR but not LaminA/C or Nesprin-2. Reciprocally, from cells overexpressing GFP-Tau, we were able to pull-down SUN1 and LBR (clients) together with GFP-Tau (bait) from nuclear lysates (Figure 4B, Figure EV3B,C,D). These data confirmed a physical interaction of Tau with LBR and SUN1. Notably, SUN1 is part of LINC complexes that are responsible for mechanical force transduction from the cytoskeleton onto chromatin (Carley *et al*, 2021). Tau effects on LINC have been recently suggested in FTD-Tau expressing cells (Sohn *et al*, 2023).

**Figure 4.**
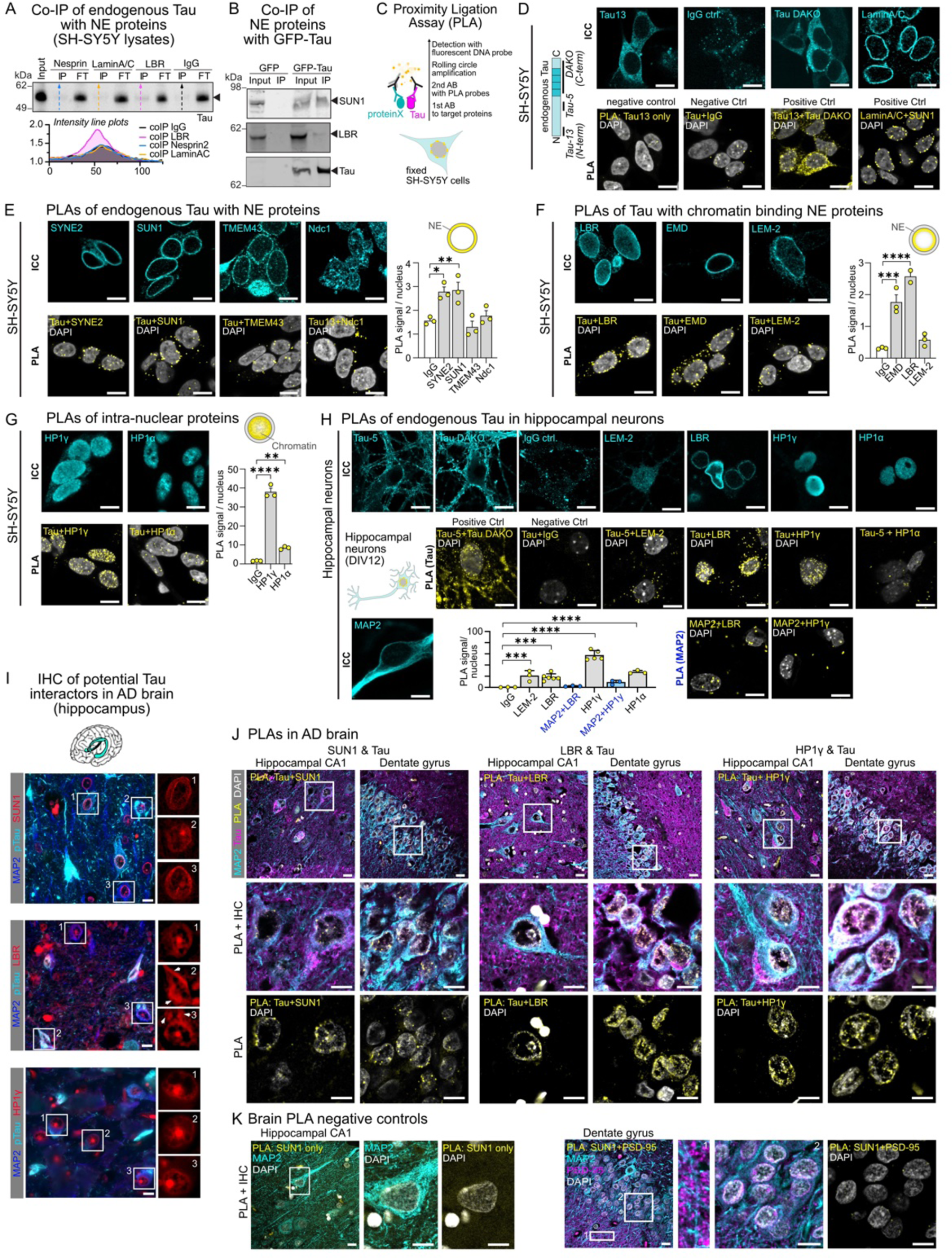
Tau interacts with NE-associated chromatin organizers in cells, neurons, and the brain. (A) Co-immunoprecipitation (Co-IP) of endogenous Tau with NE proteins Nesprin, Lamin A/C and LBR in SH-SY5Y cell lysates using high-salt buffer for cell lysis. Input (total lysate) and pulldown (IP) samples for NE proteins and control IgG are shown. The flow-through (FT) fraction is also included. Tau is detected at ∼55 kDa. Intensity line plots along gel lanes (direction indicated by dashed lines) show increased signal over background for LBR. (B) Co-IP of SUN1 and LBR with GFP-Tau in SH-SY5Y cell lysates using high-salt buffer for cell lysis. (C) Principle of proximity ligation assays (PLAs) for *in situ* detection of proteins in close proximity (<40 nm) to Tau. (D) Positive and negative controls for endogenous Tau PLAs in naive SH-SY5Y cells. The negative control (Tau+IgG) shows few unspecific fluorescent PLA signal spots, whereas the Tau positive control using two Tau antibodies (Tau-13+Tau DAKO) results in high PLA signal throughout the cell body. The positive control for nuclear envelope localization, LaminA/C+SUN1, leads to PLA signal around the nucleus. Scale bars = 10 µm. (E) PLA of endogenous Tau with NE proteins in SH-SY5Y cells. Quantification of PLA signal per nucleus indicate Tau proximity to SUN1 and Nesprin-2. PLA signal quantified in nuclear ROIs. n = 200-400 nuclei per condition. One-way ANOVA with Dunnett post-test. Scale bars = 10 µm. (F) PLA of endogenous Tau with inner NE proteins in SH-SY5Y cells. Quantification of PLA signal per nucleus indicated Tau proximity to Emerin and LBR. PLA signal quantified in nuclear ROIs. n = ∼200 per condition. One-way ANOVA with Dunnett post-test. Scale bars = 10 µm. (G) PLA of endogenous Tau with intranuclear proteins in SH-SY5Y cells. Quantification of PLA signal per nucleus validates HP1γ as interactor of Tau. PLA signal was quantified in nuclear ROIs. n = ∼300 nuclei per condition. One-way ANOVA with Dunnett post-test. Scale bars = 10 µm. (H) PLA of endogenous Tau in mouse hippocampal neurons (DIV12). The negative Control (Tau+IgG) leads to few unspecific fluorescent spots, whereas Tau positive control (Tau-5+Tau DAKO) results in high PLA signal throughout the soma and projections. PLA signal quantified in nuclear ROIs. Note, strong PLA signal is produced for Tau+LBR and Tau+HP1γ, but not for MAP2+LBR and MAP2+HP1γ. n = ∼50 nuclei per condition. One-way ANOVA with Dunnett multiple comparison test. Scale bars = 10 µm. (I) Immunostaining of potential Tau interactors (SUN1, LBR, HP1γ) in human AD brain sections (hippocampal CA1). Zoom-ins show selected neuronal nuclei/cell bodies having different phospho-Tau (pS202/pT205/pT231) levels and aggregation states. Note, LBR seems to co-aggregate with p-Tau in NFTs in neuronal somata (arrow heads in zoom-ins). Scale bars = 10 µm. (J) Immunostaining for MAP2 (cyan) and total Tau (pink) in combination with PLAs (yellow) for total Tau with SUN1 (left), LBR (middle), and HP1γ (right) in AD brain sections (hippocampal CA1 and dentate gyrus). White squares indicate location of zoom-ins shown below for combined IHC+PLA and PLA with DAPI. Scale bars = 10 µm. (K) PLA negative controls (SUN1 only, hippocampal CA1; SUN1+PSD-95, dentate gyrus) show no PLA signal in human brain. Scale bars = 10 µm.

To test whether endogenous Tau would be close to other identified potential Tau interactors in the intact cellular environment, and whether these potential interactions would actually occur in or at the nucleus, we performed microscopy-based proximity ligation assays (PLAs). In these assays, two proteins – Tau and a respective partner protein – are labeled with specific primary antibodies in otherwise untreated, fixed cells or tissue, and their close proximity (<40 nm) is detected upon local fluorescent signal generation and amplification (= foci at interaction locations; Figure 4C). To outline the nuclei, DNA was labeled in parallel using DAPI. First, specificity and background signal of PLAs were detected in human neuronal cells (SH-SY5Y line) using a human Tau specific antibody (Tau13, mouse, Tau N-terminal end) alone or in combination with non-binding IgG (rabbit) as negative controls (no PLA signal detected), or Tau13 together with an antibody targeting another region of Tau (Tau DAKO, rabbit, Tau C-terminus) as positive control (strong cytoplasmic PLA signal, few PLA spots in the nucleus) (Figure 4D). A combination of antibodies against SUN1 (epitope in NE intraluminal space) and LaminA/C, interacting at the inner nuclear membrane (INM), was used as positive control to produce NE PLA signal. Additionally, for each individual antibody used in PLAs, the cellular binding pattern was first verified by immunofluorescence. To test the proximity of endogenous Tau with potential nuclear interactors, we quantified the PLA signal produced at the NE for NE proteins (Figure 4E,F) and inside the nucleus for intranuclear proteins (Figure 4G) in comparison to the IgG negative control. Among NE candidate interactors, we found the LINC complex proteins SYNE2 (Nesprin-2) and SUN1 and the INM embedded chromatin binders, EMD and LBR to produce significant PLA signal with Tau (Figure 4E,F).

In addition, we also probed the proximity of Tau with the histone associated proteins HP1α and HP1γ, which mediate the binding of inner NE proteins to chromatin and have been shown to interact with Tau *in vitro* (Abasi *et al*, 2024), as well as of the heterochromatin marks H3K9me3 and H4K20me3 because Tau has been reported to stabilize chromatin compaction (Rico *et al*, 2018). For the HP1 proteins, strong PLA signal occurred throughout the nucleus (Figure 4G). For heterochromatin histone marks, weaker but still significant signal was detected (Figure EV3E). This indicated that Tau interacted with additional chromatin-associated proteins.

Next, we tested whether the proximity of endogenous Tau with NE chromatin binders, HP1 proteins, and heterochromatin marks would occur in neurons. PLAs in primary hippocampal mouse neurons (DIV12) showed significantly enriched signal for LEM-2, LBR, HP1γ, and HP1α, but not for any of the histone marks tested (Figure 4H; Figure EV3F). Furthermore, to test whether the proximity to inner NE proteins, i.e., LBR and its binding partner HP1γ, was specific to Tau or would also occur for related proteins, we performed PLAs of LBR and HP1γ with MAP2, another neuronal microtubule-associated protein with a domain organization and structure similar Tau and of constitutive high abundance in the somatodendritic compartment. MAP2 did not produce PLA signal with LBR and Hp1γ (Figure 4H). This indicated that the potential intranuclear interaction with chromatin binding proteins were specific to Tau. Notably, we could not test the proximity of Tau to SUN1 or SYNE2 (Nesprin-2) in primary neurons by PLA because of a lack of antibodies enabling specific immunodetection of the murine protein variants.

### Tau in proximity to inner NE chromatin binders in the human brain

To test whether our observation of Tau in proximity to chromatin organizing protein in the inner NE would also occur in the human brain, we performed PLAs in FFPE-fixed human brain sections. We chose to perform the assays in AD brains, i.e., hippocampal CA1 area and dentate gyrus, because of the abundant Tau missorting and pathology in these brain regions. First, we verified the specificity of antibody binding in human brain sections for SUN1, LBR, HP1γ, and LEM-2 by immunofluorescence (Figure 4I, Figure EV4A,B). SUN1 in hippocampal neurons showed the expected NE localization as well as a single condensed circular accumulation in the center of the nucleus. Similarly, LBR localized to the nuclear envelope and formed a single condensed spot inside nuclei (Figure 4I). Additionally, LBR co-localized with cytoplasmic Tau aggregates, supporting previous observation of LBR in optogenetically induced small FTD-mutant Tau clusters (Essepian *et al*, 2025; Jiang & Wolozin, 2021). HP1γ had a peculiar distribution with strong signal in a single condensed accumulation inside the nuclei and additional weak signal at the NE and at strings connecting the NE with the inner nuclear accumulation; reminiscent of a “cart-wheel” shape. This DNA shape was found more frequently in hippocampal neurons of AD than of control brains (Figure EV4C).

Next, we performed PLAs in combination with anti-Tau antibodies (Tau13 or Tau DAKO) (Figure 4J, Figure EV4D). In parallel to PLAs and DNA labeling by DAPI, we immunolabeled neuronal MAP2 and Tau in the same brain sections, which enabled us to identify neuronal nuclei and distinguish between neurons with different levels of Tau in their cell bodies (i.e., no Tau, diffuse Tau, aggregated Tau in NFTs). For the NE proteins SUN1, LBR and LEM-2, PLA signal occurred at the NE in CA1 and DG neurons, as well as at a single condensed area in the center of a subset of CA1 neuronal nuclei, which may resemble PLA signal produced by inner nuclear condensed accumulations of the NE proteins that we observed by immunostaining (Figure 4I, Figure EV4D). For HP1γ, PLA signal was found in all areas of DAPI signal, including the NE, the inner nuclear accumulation, and DNA “strings” bridging between the NE and the accumulation (also compare HP1γ immunostaining in Figure 4I). Negative controls for PLA in human brain, i.e., anti-SUN1 antibody alone or NE-residing SUN1 in combination with postsynaptic PSD-95 that are located far from one another, did not produce PLA signal (Figure 4K). Notably, PLA signal for Tau with chromatin associated proteins occurred in neurons with and without Tau accumulation in the soma (according to immunofluorescence), suggesting that even low soluble somatodendritic - and nuclear - Tau levels are sufficient to put Tau in proximity of these proteins. This matches our observations for PLA of soluble endogenous Tau in wildtype neurons. In summary, these data suggest that Tau in the human brain associates with inner NE chromatin organizers and their HP1 adaptor proteins, even at low nuclear Tau levels. Close proximity – indicating indirect or direct interactions - of soluble Tau with chromatin binding proteins like SUN1, LBR and LEM-2 could be relevant for Tau-induced NE distortions or be part of a physiological Tau function in chromatin regulation, or both.

### Tau interactions with LBR and SUN1 induce NE invaginations and nuclear inclusions

Having confirmed the interaction of Tau with LBR and SUN1, we aimed to investigate whether they would contribute to Tau-related nuclear envelope invaginations. Removal or knockdown of LBR and SUN1 likely destabilize the NE, thereby prohibiting to test whether these proteins would be necessary for Tau-related NE distortions. We therefore generated expression constructs that enabled us to artificially enhance the interaction of Tau with LBR and SUN1 using the cell permeable synthetic rapamycin analogue rCD1 (Vester *et al*, 2022) and determine the effects on the NE by microscopy: Co-expression of Tau fused to FKBP (mRFP-FKBP-Tau) with LBR or SUN1 fused to SNAP (LBR-eCFP-SNAP, SUN1-eCFP-SNAP) in cells allowed to determine effects on the NE upon Tau overexpression and upon additional molecular coupling of Tau to LBR or SUN1. We found that overexpression of Tau was sufficient to induce the frequent occurrence of nuclear envelope invaginations and circular nuclear, Tau-filled “inclusions” (Figure 5A). Enhancing the interactions of Tau with LBR and SUN1 – through coupling of FKBP-Tau to LBR-SNAP or SUN1-SNAP using rCD1 compound - further enhanced the occurrence of NE invaginations and inclusions (Figure 5A,B). Thereby, the number of inclusions was favored over the number of invaginations in the presence of rCD1. Detailed microscopic evaluation of the inclusions suggested that the circular Tau inclusions originated from NE protrusions (= invaginations) into the nucleoplasm, representing cytoplasm inclusions enveloped by parts of the NE containing LBR and SUN1. This idea was confirmed by the presence of soluble Tau and tubulin inside circular inclusions, and the presence of NE proteins - LBR, SUN1, LaminB1, LaminA/C and nuclear pore complexes - in inclusion “envelopes” (Figure 5C). Further, the nuclear inclusions did not co-localize with other prominent nuclear organelles, i.e., nucleoli (NPM1+) and nuclear speckles (SRRM2+). Similar results were obtained for co-expression of TauP301L with LBR and SUN1 (Figure EV5A,B).

**Figure 5.**
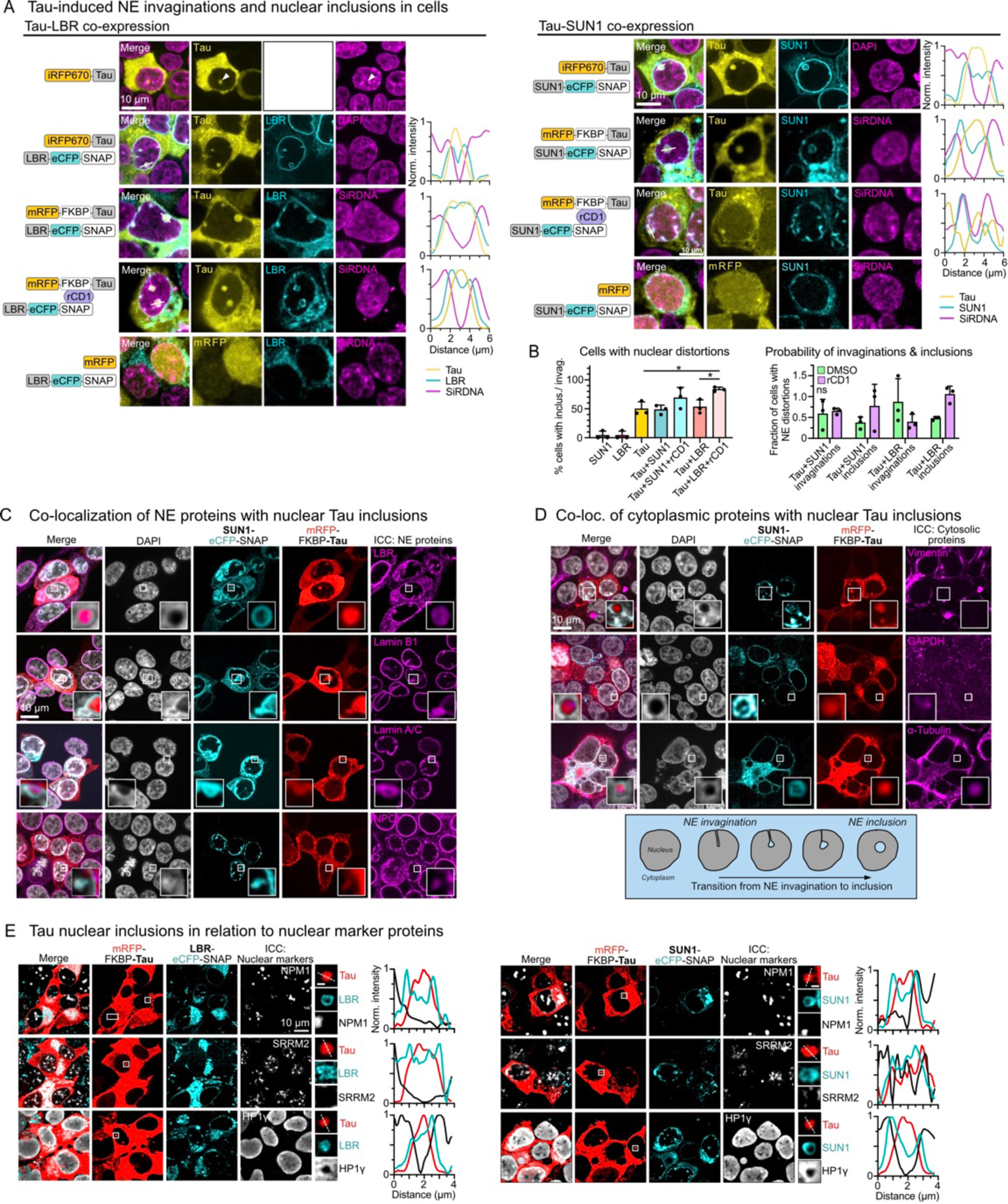
Tau induced NE invaginations and cytoplasm-filled nuclear inclusions. (A) Tau-induced nuclear envelope (NE) invaginations and nuclear inclusions in HEK293 cells. Cells expressing iRFP670–Tau or mRFP-FKBP-Tau alone, or in combination with LBR–eCFP–SNAP (left) or SUN1–eCFP–SNAP (right) in the absence or presence of the chemical coupler rCD1 inducing FKBP-SNAP binding. Normalized fluorescence intensity line plots along white lines in merged images illustrate the spatial relationship between Tau, LBR or SUN1, and DNA in/at NE invaginations and inclusions. Scale bars = 10 µm. (B) Percentage of cells with NE invaginations and/or inclusions (left) upon mRFP-FKBP-Tau overexpression alone or in combination with LBR-eCFP-SNAP or SUN1-eCFP-SNAP, with and without rCD1 addition. Fractions of cells having NE invaginations or inclusions (right) in mRFP-FKBP-Tau plus SUN1-eCFP-SNAP (Tau+SUN1) and mRFP-FKBP-Tau plus LBR-eCFP-SNAP (Tau+LBR) expressing cells, with and without rCD1 treatment. Data shown as mean±SD, N=20-99 cells from 3 experiments, one-way ANOVA with Tukey post-test. (C) Co-localization of NE proteins with nuclear Tau inclusions. Cells co-expressing mRFP–FKBP–Tau and SUN1–eCFP–SNAP were immunostained for endogenous NE proteins LBR, Lamin B1, Lamin A/C, and nuclear pore complex (NPC). White squares indicate regions shown at higher magnification. Scale bar = 10 µm. (D) Co-localization of cytoplasmic proteins with nuclear Tau inclusions. Cells co-expressing mRFP–FKBP–Tau and SUN1–eCFP–SNAP were immunostained for cytoplasmic proteins (vimentin, GAPDH, and β-tubulin). White squares indicate regions shown at higher magnification. Scale bar = 10 µm. Box shows proposed model for transition of invaginations into nuclear cytoplasm inclusions (E) Tau nuclear inclusions in relation to nuclear organelle markers. Cells co-expressing mRFP–FKBP–Tau and LBR–eCFP–SNAP (left) or SUN1–eCFP–SNAP (right) were immunostained for nucleolar markers NPM1, nuclear speckle protein SRRM2, and chromatin binding protein HP1γ. White squares indicate regions shown at higher magnification used for normalized intensity line plots. Scale bar = 10 µm.

Importantly, intensity line plots through individual Tau inclusions showed that DNA (DAPI signal) and the DNA-binding protein HP1γ were enriched around Tau inclusions, immediately adjacent to the LBR/SUN1 containing NE envelopes, likely through coupling of DNA to LBR, SUN1, and other chromatin binding proteins in the inner NE (Figure 5C,D,E). This DNA condensation at the surface of abnormal, Tau-induced nuclear inclusions and NE invaginations could have a large impact on chromatin regulation and, hence, gene expression.

### Tau condenses with LBR in on DNA

We were especially intrigued by the identification of LBR as nuclear Tau interactor. Tau/LBR interactions at the inner NE could change LBR’s DNA binding, thereby impacting chromatin arrangement and gene expression. LBR links chromatin to the nuclear envelope through its N-terminal, nucleoplasmic projection domain (LBR-npd), directly through its DNA interaction domain (Tudor domain, aa 1-89) or via binding of HP1 proteins to its globular II domain (aa 90-211) (Olins *et al*, 2010) (Figure 6A). PONDR plots along the amino acid sequence of LBR revealed that its N-terminal region (LBR-npd), which is protruding from the INM into the nuclear lumen and encodes the DNA binding activity, was especially disordered (Figure 6A). Hp1γ, which was reported to mediate LBR binding to DNA (Polioudaki *et al*, 2001) and form condensates in the presence of DNA (Li *et al*, 2023), showed a disordered region in the middle of its sequence. In addition, when looking at structural characteristics of LBR-npd, HP1γ, and other potential nuclear Tau interactors, we found that also LEM-2, and Emerin had predicted high levels of protein disorder (PONDR score > 0.5) and/or had, like Tau (MAPT), the tendency to form condensates/granules (CatGRANULE score > 0.75 (Bolognesi *et al*, 2016); PhasePro (http://predict.phasep.pro/ (Chen *et al*, 2022)) (Figure EV6A,B). Based on these characteristics, we hypothesized that LBR-npd could form condensates with DNA and that the presence of Tau may alter the interaction of LBR-npd with DNA.

**Figure 6.**
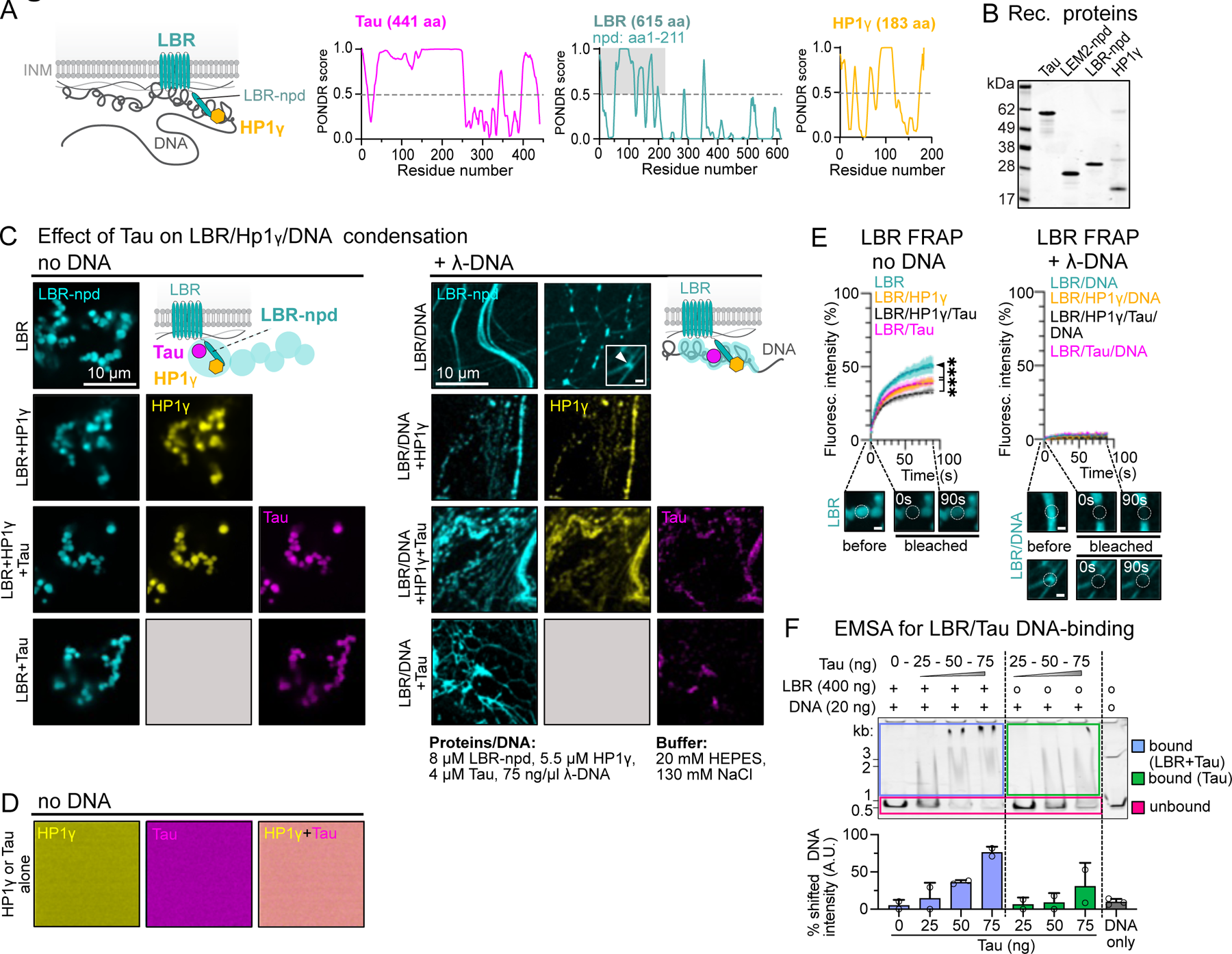
Tau changes LBR-npd condensates and its binding to DNA. (A) Schematics of the inner nuclear membrane protein LBR and the interaction of its N-terminal nuclear projection domain (npd) with DNA - direct or via HP1 proteins. In addition, PONDR predictions of protein disorder along the peptide sequences of LBR, HP1γ and Tau are shown. LBR-npd, the central region of HP1γ, and Tau’s N-terminal projection domain are largely disordered (PONDR > 0.5). (B) SDS-PAGE of purified recombinant Tau, LEM2-npd, LBR-npd, and HP1γ used for *in vitro* assays. (C) LBR-npd (including 1% LBR-npd-Dylight488) condensation in the presence and absence of HP1γ (including 1% HP1γ-Dylight555), Tau (including 2% Tau-Dylight650), or both. Left: in the absence of DNA, LBR-npd readily forms grape-like assemblies of spherical condensates, into which HP1γ and Tau co-partition. In the presence of λ-DNA (right), LBR-npd attaches to and bundles DNA. HP1γ joins LBR-npd on thin DNA threads an in thicker LBR-npd condensates. Tau is found only in condensed LBR-npd on DNA. Scale bars = 10 µm. (D) Control conditions showing no condensation of HP1γ, Tau, or both with and without λ-DNA in the used assay buffer conditions. Scale bars = 10 µm. (E) FRAP of LBR-npd shows liquid-like recovery in the absence of DNA, whereby HP1γ and Tau reduce the mobile fraction of LBR-npd independently and co-cooperatively. Bound to DNA, LBR-npd shows no FRAP regardless of the absence or presence of HP1γ and/or Tau. Data shown as mean±SD, n=10-15 condensates per condition from 3 experiments. One-way ANOVA with Tukey post-test comparing the average of last 10 mean values of each condition. Scale bars = 1 µm. (F) Electrophoretic mobility shift assays (EMSA) of LBR-npd (400 ng) binding to DNA (500 bp, 20 ng) in the presence of 0, 25, 50, or 75 ng Tau. Upshift of DNA signal in gel lanes (blue/green rectangles) indicates binding of added proteins to DNA. Unbound DNA runs at the lower end of the gel (pink rectangle). Bottom: Quantification of % bound DNA upon addition of LBR-npd and Tau (blue), or Tau only (green). N=2 independent experiments.

Using recombinant Tau, LBR-npd, and HP1γ proteins (Figure 6B), we first tested whether LBR-npd would show phase separation behavior – alone and in combination with DNA – by microscopy. Because it is unclear which DNA base pair sequences LBR is binding to, we used λ-phage DNA (λ-DNA, 47 kbp) covering a wide spectrum of base pair sequences. Without DNA, LBR-npd by itself readily formed small spherical condensates (diameter 1-2 μm) that assembled into larger, grape-like clusters (Figure 6C, left). Tau and HP1γ, or both together, efficiently partitioned into and co-enriched in LBR-npd condensates but did not form condensates in the absence of LBR-npd (Figure 6D). Fluorescence recovery after photobleaching (FRAP) of condensed LBR-npd showed that Tau and HP1γ cooperatively reduced the LBR-npd molecular mobility inside condensates (Figure 6E), indicating that Tau and HP1γ may independently interact with LBR-npd, possibly at separate binding sites.

Next, we added DNA to our experimental setup. The addition of λ-DNA to LBR-npd, under the same buffer conditions, changed the assembly of LBR-npd: the grape-like LBR-npd condensate assemblies disappeared and LBR-npd attached to the added DNA strands (Figure 6C, right; Figure EV6C), resulting in thick DNA bundles coated by LBR-npd. In less coated DNA regions, LBR-npd condensates were attached to and bridging between thinly LBR-npd coated DNA threads. This arrangement of LBR-ndp on DNA was reminiscent of the wetting of protein condensates on cytoskeletal fibers (King & Petry, 2020). In the presence of HP1γ, Tau, or both, LBR-npd still associated with DNA, and HP1γ and Tau colocalized with the larger condensed LBR-npd clusters. FRAP of LBR-npd associated with DNA revealed a mobile molecular fraction of only ∼3% (Figure 6E, Figure EV6D) in all conditions, indicating a tight association of LBR-npd with DNA, regardless of the presence of HP1γ or Tau.

To further assess how the presence of Tau could influence the direct interaction between LBR and DNA, we examined the binding of LBR-npd to DNA by electrophoretic mobility shift assays (EMSAs; Figure 6F). Mixing DNA (500 bp fragment; 20 ng = 1 ng/μl) with LBR-npd (400 ng = 20 ng/μl) in the presence of increasing Tau concentrations (0, 25, 50, 75 ng = 0, 1.25, 2.5, 3.75 ng/μl) showed that already small amounts of Tau (25 ng) substantially increased the binding of LBR-ndp to DNA (upshift of signal for DNA/protein complexes). This suggested a strong activity of Tau in enhancing HP1γ-independent binding of LBR-npd to DNA, notably, below LBR-npd concentrations needed for microscopic condensation. Interestingly, Tau alone also bound DNA already at low concentrations, starting at 50 ng/μl (Figure 6F), which supports previous observations of Tau binding to DNA (Sultan *et al*, 2011; Benhelli-Mokrani *et al*, 2018; Hua *et al*, 2003; Sjöberg *et al*, 2006).

Together, these data suggest that low levels of intranuclear Tau can enhance HP1-independent binding of LBR-npd to chromatin, whereas higher nuclear Tau levels may alter LBR’s homotypic interactions and HP1-binding by changing its condensation behavior.

### Elevated Tau levels modulate neuronal chromatin

Previous reports suggested that Tau could modulate neuronal chromatin in transgenic flies and AD brains (Frost *et al*, 2014; Klein *et al*, 2019). Our data indicated that Tau-induced NE invaginations and nuclear inclusions induce abnormal, local DNA condensation (Figure 5) and that Tau can influence the binding of LBR to HP1 and DNA (Figure 6). To better understand the connection between increased somatodendritic and nuclear Tau levels and neuronal chromatin organization, we investigated histone modifications associated with gene repression, H3K9me3 (marking constitutive heterochromatin (Padeken *et al*, 2022)), and gene activation, H3K9ac (marking enhancers or promotors (Gates *et al*, 2017)) by immunofluorescence (Figure 7A,B). Compared to GFP-expressing control neurons (Tau base), the number per nucleus but not size of H3K9me3-positive chromocenters (ChrCs) increased in Tau OE and TauPL OE neurons (Figure 7C; Figure EV7A,B). Notably, not all H3K9me3-positive chromocenters also contained highly condensed DNA (= high DAPI intensity). Tau KD did not change H3K9me3-positive ChrCs. Western blots of neuronal lysates confirmed these results (Figure EV7C,D). For H3K9ac, Tau OE (ns), TauPL OE, and Tau KD all increased general nuclear fluorescence intensity outside of ChrCs compared to control (Tau base) neurons. Western blots showed the same trend but remained none-significant.

**Figure 7.**
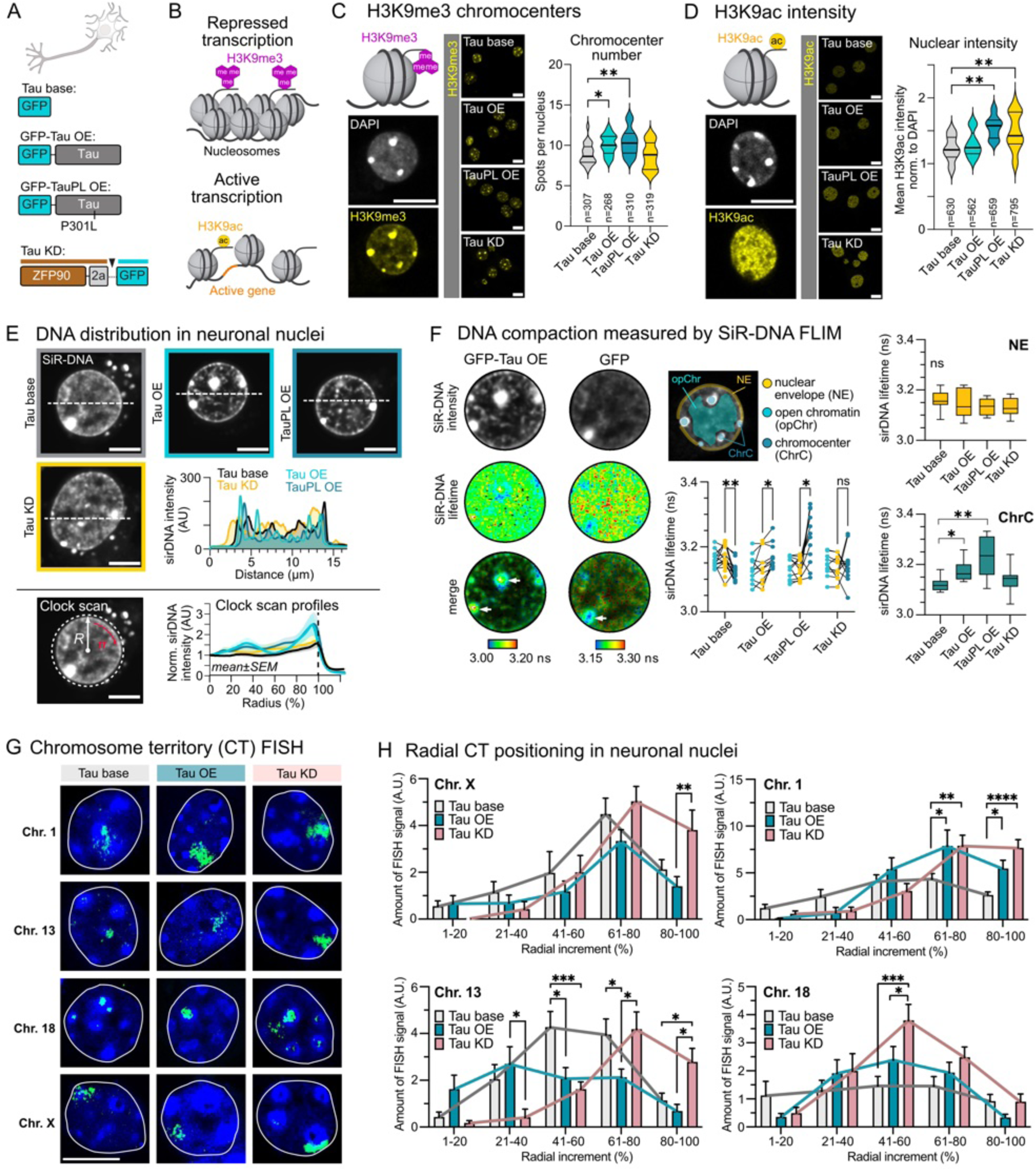
Somatodendritic and nuclear soluble Tau induces heterochromatin alterations. (A) Modulation of Tau levels in primary mouse hippocampal neurons via adeno-associated viruses (AAVs) used to study heterochromatin alterations. Overexpression of GFP (endogenous Tau levels, Tau base), GFP-tagged human wild-type Tau (2N4R isoform) (Tau OE), FTD-mutant GFP-Tau^P301L^ (TauPL OE), or the gene silencing zinc-finger protein transcription factor (Wegmann *et al*, 2021), ZFP90, that decreases endogenous Tau levels (Tau KD). (B) Histone modifications associated with gene repression (H3K9me3) and gene activation (H3K9ac). (C) H3K9me3 chromocenter number in neurons with different Tau levels. n = number of nuclei (indicated), 3 biological replicates, and 3 images analyzed per condition. One-way ANOVA with Dunnett post-test. Scale bars = 10 µm. (D) H3K9ac immunofluorescence intensity in neurons with different Tau levels. n = number of nuclei (indicated), 3 biological replicates, and 3 images analyzed per condition. One-way ANOVA with Dunnett post-test. Scale bars = 10 µm. (E) DNA (siR-DNA) distribution in nuclei of neurons with different Tau levels. Representative images of individual nuclei for each treatment. Line-scan intensity profile (middle right) show siR-DNA intensity along white dashed lines in images. Lower right: Radial DNA profiles (clock scan analyses; n= 5-13 nuclei) show DNA enrichment at the nuclear envelope in Tau OE and TauPL OE compared to Tau base and Tau KD neurons. Scale bars = 5 µm. (F) FLIM of siR-DNA in neurons with different Tau levels. Compaction/condensation of DNA correlates with a decrease in siR-DNA lifetime. Representative intensity and lifetime maps of nuclear ROIs are shown for GFP (Tau base) and GFP-Tau (Tau OE) expressing neurons. White arrows indicate position of chromocenters. Middle and right: Quantification of most probable SiR-DNA lifetimes for nuclear sub-compartments: Chromocenters (ChrC), open chromatin (opChr), and nuclear envelope (NE). Note that Tau OE and TauPL OE nuclei show increased siR-DNA lifetime in ChrCs. n = 11-17 nuclei per condition from two experiments. Data shown as ChrC, NE, and opChr for individual nuclei (before/after plot; Two-way ANOVA with Tukey post-test) and for comparison of NE and ChrC between groups (box-whisker plots, median with 5-95 percentile, one-way ANOVA with Tukey post-test. (G) Representative chromosome FISH images collected from non-transduced (Tau base), Tau OE, and Tau KD neurons show the positioning of chromosome territories (CT) in the equatorial plane of neuronal nuclei for chromosomes ChrX, 1, 13, and 18. DNA (DAPI) is shown in blue, CTs in green. Scale bar = 10 µm. (H) Radial positioning of chromosomes ChrX, 1, 13, and 18 in the equatorial plane of neuronal nuclei with different Tau levels. Radial positioning of Chr’s was determined by measuring the absolute FISH signal intensity across concentric nuclear shells from the nuclear interior (1–20%) to the periphery (81–100%). N = 9-19 nuclei per condition, two-way ANOVA with Tukey post-test.

These data indicated that increased cytosolic and nuclear Tau levels elevate H3K9me3-associated chromatin repression, while simultaneously increasing gene activation related to H3K9Ac. Tau reduction led to gene activation. Together this suggests that somatodendritic Tau mis-sorting, and the simultaneous increase in the amount of nuclear Tau, may favor chromatin repression, and that high Tau levels induce additionally gene activation.

### Tau changes chromatin association with the nuclear envelope

In addition to histone modifications, the spatial organization and condensation of interphase chromatin serves as a higher-order regulatory level of gene expression. Thereby, DNA-dense ChrCs and the nuclear envelope play important roles. Both ChrCs and chromatin associated with the nuclear lamina (lamina-associated domains, LADs) contain mostly repressed genes (van Steensel & Belmont, 2017). We tested whether modulating Tau levels would change DNA distribution and condensation in neuronal nuclei. Nuclei of untreated or GFP-expressing neurons (Tau base) and Tau KD neurons had homogenously distributed DNA outside of chromocenters (diffuse DAPI signal) (Figure 7E). Nuclei of neurons with elevated Tau (Tau OE and TauPL OE) seemed to have a higher concentration of DNA at the NE, whereas their nucleoplasm seemed largely void of diffuse DNA signal, with few, small DNA-dense “spots”. Intensity line plots across nuclei and analysis of average radial DNA distribution (clock scan) confirmed this observation.

Next, we tested whether the redistribution of DNA in Tau OE and TauPL OE neurons was associated with changes in DNA compaction, which is thought to somewhat correlate to gene repression. Fluorescence lifetime imaging (FLIM) of siR-DNA (siR-Hoechst) labeled DNA in living neurons. siR-DNA is a far-red, cell permeable live cell DNA probe that can be used to estimate DNA compaction based on its fluorescent lifetime being quenched with increasing DNA density (Hockings *et al*, 2020). Using this approach, we found that neurons with all Tau levels had similar siR-DNA lifetimes at the NE (Figure 7F), indicating that the redistribution of DNA towards the NE in Tau OE and TauPL OE neurons was not associated with stronger DNA compaction at the NE. Surprisingly, however, Tau OE and TauPL OE neurons showed strongly increased Sir-DNA lifetimes – indicative of DNA decondensation - in the core of DNA ChrCs. The Sir-DNA lifetimes of the remaining parts of the nucleoplasm (open chromatin) were again similar in all conditions.

Finally, we tested whether Tau may induce changes in the nuclear positioning of specific chromosomes. In the interphase nucleus, chromosomes occupy distinct territories in the 3D-volume of the nucleus (Cremer & Cremer, 2001). The spatial arrangement of chromosome territories (CTs) can influence the expression of harbored genes. Certain chromosomes (Chr’s), e.g., Chr1, Chr13, ChrX, and Chr18 have been shown to preferentially locate at the nuclear periphery in interphase nuclei of mammalian cells (Mehta *et al*, 2010). Interestingly, LBR was shown to contribute to ChrX silencing by associating it to the NE (Chen *et al*, 2016). Furthermore, ChrX was shown to re-localize from the NE into the nuclear interior in epileptic neurons of human brains (Borden & Manuelidis, 1988), a condition that is also linked to somatodendritic Tau re-sorting upon neuronal hyperactivity.

We performed fluorescent in-situ hybridization (FISH) of chromosomes (Chromosome PAINT) to test whether modulation of Tau levels would change the radial positioning of these chromosomes in neuronal nuclei (Figure 7G,H). Analyzing the equatorial plane of nuclei labeled for individual chromosomes, we found that the LBR-associated ChrX was shifted towards the nuclear periphery upon Tau reduction (Tau KD) compared to untreated neurons (Tau base). Chr1 was shifted towards the periphery in both Tau OE as well as TauKD neurons. Chr13 shifted from the nuclear center to the periphery in Tau KD neurons and kept its radial positioning but dispersed (= broader radial distribution) in Tau OE neurons. Different from the other chromosomes tested, Chr18 was located in DAPI-positive ChrCs; it appeared to condense half-way between the nuclear center and the periphery upon Tau KD. Together, these data indicate that Tau levels can have a profound influence on CT organization in neuronal nuclei, whereby the removal of Tau seems to have the strongest effect. This suggests that low levels of endogenous nuclear Tau - present in naïve neurons – are sufficient to influence chromosome organization, however, with so far unknown effects on the expression of genes encoded by the affected chromosomes.

In summary, our data suggest that both elevation and reduction of Tau modulate neuronal chromatin on multiple levels, including histone modification, DNA distribution, and CT positioning.

### Elevated Tau upregulates transcription factors regulating stress response pathways

The identified Tau-related changes in chromatin regulation (Figure 7) and LBR-DNA interactions (Figure 6) suggested that somatodendritic Tau missorting would influence gene expression. To examine which genes are responsive to changes in somatodendritic/nuclear Tau, we sequenced the RNA isolated from neurons containing different Tau levels (Tau base, Tau OE, and Tau KD; Figure 8A). To minimize contribution of glial RNA, the primary cultures where treated with AraC to kill dividing cells (Figure EV8A).

**Figure 8.**
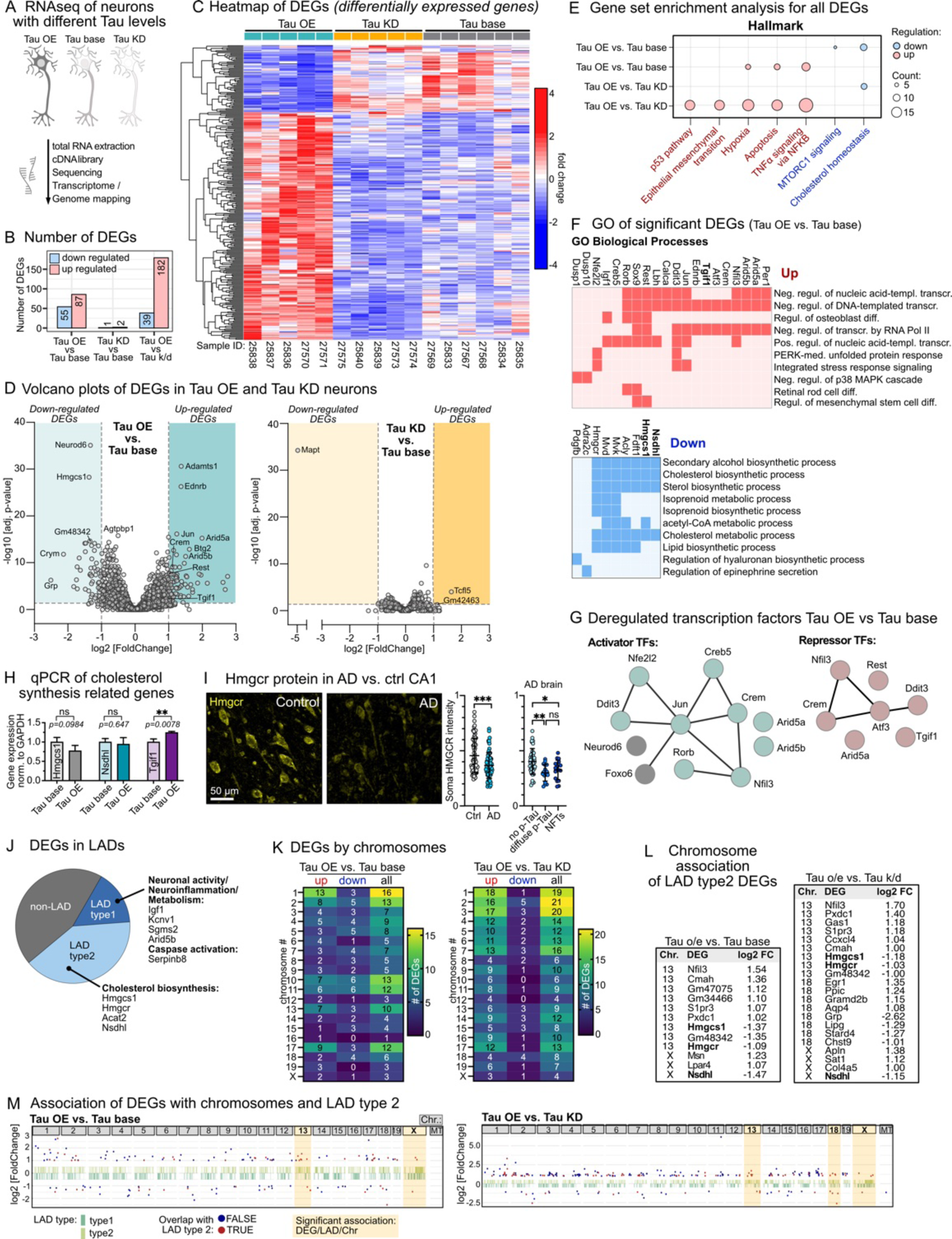
Impact of Tau levels on neuronal transcription. (A) Schematics of RNAseq (total RNA, rRNA depletion) in Tau base, Tau OE, and Tau KD neurons. (B) Number of significant DEGs in Tau OE and Tau KD compared to Tau base, and Tau OE compared to Tau KD neurons. Data were collected from N = 5-6 neuronal cultures. (C) Heatmap of significant differentially expressed genes (DEGs) in Tau OE, Tau KD and Tau base neurons. (D) Volcano plots of DEGs in Tau OE vs Tau base, and Tau KD vs Tau base neurons. Threshold of significance was set to log2[Fold Change] > 1 and adjusted p-value < 0.05. Names of selected genes (mouse) are indicated. (E) Gene set enrichment “Hallmark” for all DEGs. Up and down-regulated hallmark pathways are shown for Tau OE vs Tau base, and Tau OE vs Tau KD. (F) Gene ontology (GO *Biological Processes*) of significant DEGs in Tau OE compared to Tau base neurons are shown separately for up-(top) and down (bottom)-regulated DEGs. (G) Networks of transcription factors (TFs) among significant DEGs in Tau OE vs. Tau base neurons, grouped by their (main) activity as transcriptional repressors and activators. Notably, except *Neurod6* and *Foxo6*, all TFs are upregulated in Tau OE neurons (see Figure EV10B). Quantitative PCR (qPCR) of selected genes (*hmgcs1, nsdhl* and *tgif1*) involved in cholesterol biosynthesis in Tau base and Tau OE neurons. Expression values were determined from RNA isolated from neuronal cultures and normalized to GAPDH expression. Data shown as mean±SD, n=3 experiments. (H) Hmgcr protein levels are reduced in AD brain. Representative immunofluorescence images and quantification of Hmgcr signal intensity in neuronal somata in control and AD hippocampal sections. Student’s t-test. For AD, signal intensity of neurons having no phosphorylated Tau (no p-Tau), diffuse p-Tau, or aggregated p-Tau in neurofibrillary tangles (NFTs) are compared as well. One-way ANOVA with Tukey post-test. Scale bar = 50 µm. (I) LAD association of DEGs in Tau OE compared to Tau base neurons. Of 142 total DEGs, 22 DEGs (15%) are located in LADs type1 and 57 DEGs (40%) are located in LAD type2. Selected gene names of DEGs for each LAD type are indicated. LAD type2 genes include cholesterol biosynthesis genes *Hmgcr*, *Hmgcs1*, and *Nsdhl* that are downregulated in Tau OE neurons. (J) Chromosome location of DEGs in Tau OE compared to Tau base (left) or Tau KD (right) neurons, for up-and down-regulated and all DEGs. Number in heat map gives exact number of DEGs in chromosomes. (K) Gene names of DEGs overlapping with LAD type2 and significantly enriched in chromosomes Chr13 and ChrX for Tau o/e versus Tau base neurons, and in chromosomes Chr13, 18, and X for Tau OE vs. Tau KD neurons. (L) Chromosomal distribution of LAD-associated DEGs. DEGs in LADs type2 are distributed across Chr1-19 and ChrX. Significant enrichment of LAD type2 DEGs occurs in chromosomes Chr13 and ChrX for Tau OE compared to Tau base neurons (left), and in chromosomes Chr13, Chr18 and ChrX for Tau OE compared to Tau KD neurons (right).

Increasing Tau (Tau OE) led to an upregulation (n = 87) and down regulation (n = 55) of several genes (significance cut-off: fold change > 2, adjusted p-value < 0.05; Figure 8B,C,D; Figure EV8B,C,D and S9A; Supplemental Table T5,T6). Reduction of Tau (Tau KD) had comparably small effects on gene expression (significantly up-regulated genes: n = 2; down-regulated genes: n = 1 (*MAPT*)), consistent with the idea that increased concentrations of Tau in and around the nucleus, as a consequence of Tau overexpression or somatodendritic resorting in neuronal stress conditions, could function as signal to initiate gene expression changes.

Gene set enrichment analysis (GSEA; considering all transcriptomic changes; Figure 8E; Figures EV9A,B) showed that increased Tau levels (Tau OE vs. Tau base) upregulated pathways of TNFα (proinflammatory) signaling, apoptosis, and oxidative stress (hypoxia) response. These pathways are associated with AD (Tao *et al*, 2024; Kumari *et al*, 2023; Decourt *et al*, 2017). In Tau OE neurons, upregulation of these pathways involved upregulation of a number of transcription factors (TFs), including activators and repressors (Figure 8F,G; Figure EV10A,B).

wPGSA gene-set analysis of all DEGs (except transposable elements, TEs), which predicts the activities of transcription regulators responsible for the observed gene expression changes (Kawakami *et al*, 2016) (wPSGA t-score indicates expression changes in genes regulated by a certain TF; TFs with a wPSGA t-score > +4 or <-4 and adjusted p-value < 10^-5^ are predicted to be involved in gene deregulation; a negative t-score indicates an overall decreased expression of the TF-regulated genes). wPGSA suggested that gene upregulation upon Tau overexpression (t-score > +4), including upregulation of TF expression, was controlled by three TFs, i.e., *Jun*, *Gcr*, and *Bmi1* (Figure EV10C). The immediate early gene *Jun* is a component of the AP-1 complex that, for example, suppresses metalloproteases in response to neuronal depolarization (Rylski *et al*, 2009). In AD, *Jun* was suggested to drive aberrant TE mobilization and to be associated with the innate immune response. In our data set, *Jun* is among the transcriptomically upregulated TFs (Figure EV10B). The glucocorticoid receptor, *Gcr*, has anti-inflammatory and immunosuppressive functions and is regulated by, as well as controls, AP-1 expression in a negative feed-back loop to regulate stress response (Karin & Chang, 2001). *Bmi1* is a major component of the polycomb group complex 1 that functions as an essential epigenetic repressor of multiple regulatory genes, e.g., involved in DNA damage repair, p53 repression and neuronal antioxidative processes (Chatoo *et al*, 2009). Notably, *Gcr* and *Bmi1* themselves were not among the significant DEGs in our data set.

For the downregulation of genes upon Tau overexpression (t-score <-4), wPSGA predicted numerous TFs to be involved, indicating that higher Tau concentrations might increase activity and/or amount of gene repressing factors. This was in line with the observed upregulation of repressive TFs (Figure EV10B). For example: (i) The c-AMP responsive element modulator *Crem*, which regulates metabolic pathways and reactions and is a negative regulator of synaptogenesis in response to neuronal activity (Aguado *et al*, 2009). (ii) The transcriptional repressor *Rest*, which becomes activated in differentiated neurons confronted with ischemic and seizure conditions to protect against neuronal death (Hwang & Zukin, 2018; Cheng *et al*, 2022), for example, by repressing the expression of pre-and post-synaptic proteins necessary for neurotransmission (McClelland *et al*, 2014). In AD, *Rest* and its protective role are downregulated (Lu *et al*, 2014; Aron *et al*, 2023), and *Rest* dysregulation is also implicated in the pathogenesis of epilepsy, Huntington’s disease, Parkinson’s disease (McClelland *et al*, 2011; Zuccato *et al*, 2003; Chuang *et al*, 2009). (iii) *Tgif1*, a TGFβ/Smad-induced repressor and regulator of the Wnt pathway that is involved regulating DNA repair and homeostasis (Canchi *et al*, 2019). In AD, *Tgif1* is downregulated and Wnt signaling suppressed (Martínez & Inestrosa, 2021). Other upregulated TFs that have previously been linked to AD and Tau pathology included the ER-stress and unfolded-protein response responsive transcription factors *Atf3* and *DDIT3/Chop*, the hypoxia-responsive factor *Hif1α,* associated with APP processing, Aβ production and Tau pathology (Alexander *et al*, 2022), the antioxidant transcription factor *NFE2L2/Nrf2*, *Rorb* that has been identified to be expressed in neuronal subtypes particularly vulnerable to Tau pathology (Leng *et al*, 2021), and *Arid5b* involved in metabolic reprogramming and upregulated in Aβ-transgenic mice (Guo *et al*, 2024; Cichocki *et al*, 2018). In addition, we detected *Foxo6*, a transcriptional activator involved in insulin signaling and regulating apoptosis, inflammation, and oxidative stress responses (Li *et al*, 2025; Hyun Kim *et al*, 2011), as well as *Neurod6*, important for neurodevelopment and differentiation and downregulated in AD brains (Satoh *et al*, 2014), to be downregulated.

Upregulated non-TF genes included the lectin-cleaving metalloprotease *Adamts1*, a GWAS risk gene for AD and suggested to improve repair and synaptic plasticity after seizure-induced neuronal damage (Gottschall & Howell, 2015; Balusu *et al*, 2023), and gastrin-releasing peptide, *Grp*, that stimulates excitatory neurons, likely by binding to *Grp* receptors on inhibitory neurons and reducing their inhibitory activity (Yang *et al*, 2017). Furthermore, we found some long non-coding RNAs (lncRNAs) to be deregulated in Tau OE neurons (Figure EV9A,C), including upregulation of *Neat1*, an essential component of paraspeckles. *Neat1* is also upregulated in AD brains (An *et al*, 2018) and has been implicated in decreasing apoptosis in response to metabolic stress (Chai *et al*, 2022), cGAS-STING-IRF3 signaling (Morchikh *et al*, 2017), and regulation of potassium channel activity (Barry *et al*, 2017).

Together, these data reveal that increased soluble Tau in the somatodendritic and nuclear compartments is sufficient to upregulate the expression of numerous TFs that stir inflammatory and cell stress-related pathways observable in neurological conditions presenting with somatodendritic Tau re-sorting, i.e., epilepsy and chronic stress (Tai *et al*, 2016; Lopes *et al*, 2016), as well as in AD and other neurodegenerative diseases (Orr *et al*, 2017). Some of the Tau-induced TF changes identified, e.g., *Rest* and *Tgif1*, are opposite from what has been observed in late-stage AD brains, indicating that they represent pro-pathological adjustments in the transcriptome occurring in response to early Tau changes.

### Elevated Tau reduces cholesterol synthesis genes

Pathways downregulated by increased Tau levels (Tau OE vs. Tau base) appeared to be TORC1 signaling, a regulator of cell metabolism and autophagy (Chantranupong *et al*, 2015; Hu *et al*, 2019) and of neuronal dendritic growth and plasticity (Li *et al*, 2009; Kovács *et al*, 2007), and cholesterol biosynthesis (Figure 8E; Figure EV9B; Supplemental Table T7). GO analysis of significant DEGs supported the reduced expression of genes involved in cholesterol, lipid, and Acyl-CoA synthesis (Figure 8F, bottom).

Among the downregulated genes were *Acly* (ATP citrate Lyase), *Hmgsc1* (3-hydroxy-3-methylglutaryl-CoA (HMG-CoA) synthase) and *Hmgcr* (HMG-CoA reductase), which catalyze the essential first steps in the mevalonate pathway of cholesterol biosynthesis. Other enzymes in the mevalonate pathway, i.e., *Mvk* and *Mvd*, as well as *Nsdhl* and *Ftfd1*, essential for later steps during the cholesterol synthesis, were also downregulated. Notably, the upregulated transcriptional repressor *Tgif1* is also involved in cholesterol biosynthesis regulation by negatively regulating *ACAT2* involved in cholesterol esterification (Pramfalk *et al*, 2014) and apolipoproteins (Melhuish *et al*, 2010). By real-time qPCR, we could verify a significant increase in *Tgif1* transcripts in Tau OE neurons (Figure 8H), similar to what has previously been observed in AD brains (Canchi *et al*, 2019). mRNA levels of *Hmgcs1* and *Nsdhl*, representatively chosen as genes directly involved in cholesterol synthesis, appeared by RT-qPCR to be non-significantly downregulated in Tau OE cultures. However, immunofluorescence showed reduced levels of *Hmgcr* protein in hippocampal neurons of AD compared to control brains (Figure 8I), which is in line with previous reports (Varma *et al*, 2021), but *Hmgcr* reduction did not correlate with the accumulation of soluble or aggregated phospho-Tau in neuronal somata.

### LAD and Chromosome association of genes deregulated by Tau

Since we found that Tau interacts with LBR and SUN (and potentially other chromatin organizing proteins at the NE), we wondered whether a signature of LAD associated genes could be found in the transcriptome data of neurons with elevated Tau levels. LADs of type1 are consistently associated with the nuclear lamina, contain constitutively repressed genes, and contribute to genome stability and cell identity, whereas LADs of type2 are flexible genome regions that are facultative repressive and facilitate dynamic, context-specific gene regulation (van Steensel & Belmont, 2017). LBR has been shown to bind LADs (Van Schaik *et al*, 2025) and to coordinate ChrX on the nuclear lamina (Chen *et al*, 2016). Determining the LAD association of significant DEGs in Tau OE neurons (assignment based on data for midbrain human neurons (Shah *et al*, 2023)), we found that > 50% of DEGs (79 of 142) were associated with LADs (LAD type1: 22 DEGs = 15% of all DEGs; LAD type2: 57 DEGs = 40% of all DEGs) in Tau OE compared to Tau base neurons, confirming a pronounced effect of Tau on genes associated with the NE (Figure 8J; Supplemental Table T9). LAD type1 assigned DEGs in Tau OE neurons did not cluster in certain GO pathways but coded, for example, for proteins involved in neuronal activity and metabolism. LAD type2 assigned DEGs clustered in cholesterol synthesis pathways (4 DEGs) and coded for a number of TFs deregulated by Tau. This indicated that elevated Tau levels influence the expression of genes located in genomic regions that are context-dependent coordinated at the NE.

To further support these findings, we performed CUT&Tag (Cleavage Under Targets & Tagmentation) for the chromatin activation mark H3K9ac, which identifies genomic positions that are activated. CUT&Tag confirmed a strong activation of intronic and non-coding distal intergenic regions in Tau OE compared to Tau base neurons (Figure EV11). Further, many genome regions activated in Tau OE neurons associated with LADs type2 as well as TEs (i.e., LINEs, SINEs, and LTRs). TEs have previously been linked to Tau pathology(Ramirez *et al*, 2022; Guo *et al*, 2018) and spreading (Liu *et al*, 2023).

We next analyzed whether Tau-responsive genes would be chromosome specific. DEGs in Tau OE neurons were distributed throughout chromosomes Chr1-19 and ChrX, with most of them located on Chr1 (16 genes; 1.0% of Chr1 genes), Chr2 (13 genes, 0.6 % of Chr2 genes), Chr10 (13 genes; 0.8% of Chr10 genes), Chr11 (12 genes; 0.5% of Chr11 genes), Chr13 (10 genes; 1.0% of Chr13 genes), and Chr17 (12 genes; 1.1% of Chr17 genes) (Figure 8K). Interestingly, Chr1, which contained the most DEGs in Tau OE neurons and was modulated by Tau levels in chromosome PAINT data (Figure 7G,H), was indeed reported to be associated with the NE and the nucleolus (Manuelidis & Borden, 1988), the two nuclear entities LAD regions attach to.

Further, we found that LAD type2 DEGs - including the cholesterol biosynthesis genes *Hmgcs1 Hmgcr,* and *Nsdhl-*, were enriched in Chr13 and ChrX (Figure 8L,M; Figure EV12). Previous reports showed that Chr13 and ChrX are located near the NE (Tanabe *et al*, 2002; Borden & Manuelidis, 1988; Zuleger *et al*, 2011), and our chromosome PAINT data (Figure 7G,H) indicate that these chromosomes may be responsive to changes in Tau levels as well.

Together, our data indicate that high somatodendritic Tau levels can alter the expression of genes located near the NE, mainly associated with facultative LADs type2, which represent genomic elements responsive to conditional cellular adaptation. This includes genes involved in cholesterol biosynthesis. These findings may suggest that increased somatodendritic soluble Tau levels – as a result of physiological stress or disease-induced Tau re-distribution – could downregulate neuronal cholesterol metabolism and upregulate neuronal stress response pathways on the genomic level by interfering with the NE association of chromatin.

## Discussion

The neuronal subcellular redistribution and somatodendritic “invasion” of Tau occurs in response to diverse physiological and pathological neuronal stimuli, including temperature changes, mechanical stimulation, and high neuronal activity. This suggests that resorting (= missorting) of Tau – and its access to somatodendritic cellular organelles/compartments like the nucleus – is part of the physiological neuronal stress-response. Upon prolonged and excessive missorting of Tau, these interactions may exert pathological effects observed in Tau-related neurodegenerative diseases, e.g., irreversible as synaptic decline, senescence, and, ultimately, neuronal death. Deciphering the physiological function of Tau missorting is therefore key to understand the reason for this frequent event, which is also is one of the initial steps in the etiology of multiple neurodegenerative diseases and conditions. Here we show that increasing soluble wildtype Tau levels in the neuronal somatodendritic compartment – to levels comparable to human AD brain neurons – is sufficient to induce nuclear envelope distortions, alterations in chromatin structure, and gene expression changes relevant for neuronal stress response and cholesterol biosynthesis. Furthermore, our data suggest that interactions of soluble Tau with chromatin organizing proteins in the inner nuclear envelope, i.e., LBR and SUN1, underlie these effects, indicating a crucial role of nuclear Tau in neuronal stress response.

### The nuclear envelope – a central role for Tau-related chromatin effects?

In recent years, multiple studies reported the correlation between Tau pathology in AD and FTD brains and in cellular and animal models with FTD-mutant Tau expression (Cornelison *et al*, 2019; Prissette *et al*, 2022; Sun *et al*, 2022; Fernández-Nogales & Lucas, 2020; Jiang & Wolozin, 2021; Monroy-Ramírez *et al*, 2013; Essepian *et al*, 2025), with abnormal neuronal nuclear shapes and NE invaginations and distortions, resembling phenotypic aspects of laminopathies (Frost, 2016; Frost et al, 2016a). It was, for example, proposed that NE deformations in primary tauopathies can be induced by aberrant Tau-related microtubule polymerization through the NE (Paonessa *et al*, 2019), or due to impaired cytoskeletal force transduction due to deficits in LINC components (Sohn *et al*, 2023). In addition, nuclear transport deficits have been reported as a consequence of phosphorylated Tau accumulating at the outer nuclear envelope (Hochmair *et al*, 2022; Kang *et al*, 2021).

Our data now reveal that the sole increase in physiological, soluble somatodendritic Tau – as it occurs in physiological stress conditions and early in AD-, which is coupled to an increase in nuclear Tau (Figure 2; (Sultan *et al*, 2011; Benhelli-Mokrani *et al*, 2018)), is sufficient to induce NE distortions. The observed effects of Tau on the NE may at least in part be induced by interactions of Tau with chromatin organizing proteins located in the inner nuclear membrane. In particular, we identify LBR and the LINC component SUN1 as interactors of Tau, both of which play important roles for the conditional NE association of genomic LADs and the repression of genes contained in these. Notably, interactions of Tau with LBR have been noticed previously (Jiang & Wolozin, 2021) but their consequences have not been further investigated. We found that Tau-induced NE invaginations and nuclear, NE-enveloped cytosol inclusions worsened when we enforced the interaction of Tau with LBR and SUN1 at the inner nuclear membrane, suggesting that direct interactions of Tau with LBR and SUN1 can induce this phenotype. Furthermore, DNA and HP1 proteins got enriched at the nucleoplasmic surface of these NE invaginations and nuclear inclusions, indicating direct, local effects of these structures on chromatin organization. These results suggest that nuclear Tau, even though being manifold less concentrated than cytosolic Tau, can have pronounced effects on the NE and associated chromatin organization.

For LBR, the interaction with Tau may involve biomolecular condensation of the two proteins together with DNA, whereby even small amounts of Tau can efficiently alter the direct interactions of LBR with DNA in the absence of HP1 “adaptor” proteins. We therefore propose that Tau/LBR interactions with DNA may be independent of HP1. Our PLA data from cells, neurons and human brain, however, indicated an interaction of Tau with HP1 proteins. Furthermore, a recent study showed that Tau impacts the binding of HP1α to DNA *in vitro* (Abasi *et al*, 2024). We currently cannot not rule out that Tau may also affect HP1-mediated DNA-binding of LBR and other NE proteins. It further remains to be clarified, whether Tau can change direct LBR/DNA interactions in a basepair sequence selective manner. Previously, A/T-rich sequences have been shown to mediate Tau/DNA binding (Sjöberg *et al*, 2006), and LADs are rich in AT as well (Meuleman *et al*, 2013), suggesting that adenine (A)-thymine (T) base pairs may be responsible for the attraction of Tau to LAD sequences. Tau ChIPseq data indicated genome-wide binding of Tau in coding and non-coding regions (∼30% lncRNAs) and suggested a gene repressor role of Tau in neurons under physiological temperature stress (Benhelli-Mokrani *et al*, 2018).

### Tau-induced chromatin re-organization at the NE

Our data show that a physiologically relevant increase in somatodendritic soluble Tau levels influences chromatin on multiple levels (heterochromatin marks, spatial DNA distribution, chromosome positioning). In our data, the “direction” (activation or repression) of chromatin changes seems, other than previously suggested for AD brains and animal models with Tau aggregation pathology (Frost *et al*, 2014), not unilateral but rather complex. The identified NE Tau interactors, LBR and SUN1, have previously been implicated in regulating interphase chromatin on the level of CTs. LBR, through interactions with the non-coding RNA Xist, couples the inactive ChrX to the lamina for repression (Chen *et al*, 2016). SUN1, together with Emerin that we find to also potentially interacts with Tau, was shown to be essential for positioning ChrX and Chr18 at the nuclear periphery (Goelzer *et al*, 2025; Bikkul *et al*, 2019).

According to a role of nuclear Tau in modulating the interactions of LBR and SUN1 with chromosomes, we find that increasing and/or removing Tau from neurons leads to nuclear repositioning of chromosomes that have been reported to bind LBR or SUN1 or otherwise locate near the NE in interphase nuclei, i.e., Chr1, Chr13, Chr18, and ChrX. To date, however, no comprehensive CT mapping data in neurons (*in vitro* or in the brain) are available, neither at base line nor in disease conditions. Comparative mapping of neuronal CTs (e.g., in AD vs. control brains) in brain regions relevant for Tau pathology in AD, e.g., the hippocampus, would be necessary to understand the actual contribution of Tau-induced CT alterations to disease-associated gene expression changes.

Our transcriptomics data further supported the activity of Tau on chromosome-NE association, showing that elevated Tau levels induce gene expression changes in T2-LAD genes in chromosomes Chr13, Chr18, and ChrX. These findings are in line with observations in human AD brain, where Tau-related genomic changes clustered in large genomic segments reflecting spatial chromatin organization and nuclear lamina association (Klein *et al*, 2019). It remains to be clarified though to which extend disease-driving gene expression changes in AD brains relate to herein described early Tau missorting and/or to later Tau pathology, and whether they depend on chromosome rearrangement or other chromatin regulatory mechanisms.

### Tau - a nuclear regulator of neuronal cholesterol synthesis and metabolism?

The Tau proximity biotinylation interactomics and the transcriptomics data sets from our study converged in an unexpected way: a subset of (potential) nuclear interactors identified for Tau where associated with cholesterol biosynthesis and mTORC1 signaling (STRING analysis of biotinylated proteins significantly enriched in Tau over control; Figure 3H); the same occurred for multiple genes significantly downregulated at elevated Tau levels (Hallmark pathway analysis; Figure 8E). In fact, LBR, which we confirmed as NE Tau interactor, plays an essential role in cholesterol synthesis by catalyzing the conversion from lanosterol to cholesterol (Tsai *et al*, 2016). The unexpected convergence of interactomics and transcriptomics data in cholesterol synthesis pathways suggests a function of Tau in regulating neuronal cholesterol metabolism, thereby provide important new evidence on the consequence and reason for Tau missorting induced by (patho)physiological stimuli.

Among the T2-LAD genes repressed at elevated Tau levels we found three key cholesterol synthesis genes: *Hmgcs1* (3-Hydroxy-3-Methylglutaryl-CoA Synthase 1), catalyzing condensation of acetyl-CoA and acetoacetyl-CoA to form HMG-CoA, and *Hmgcr* (HMG-CoA reductase), both encoded on Chr13 and essential for the first steps in the mevalonate pathway. In addition, we observed downregulation of *Nsdhl* (NAD(P)-dependent steroid dehydrogenase-like; T2-LAD gene on ChrX), and *Mvd* (mevalonate diphosphate decarboxylase) and *Mvk* (mevalonate kinase) to be downregulated. *Hmgcs1, Hmgcr, Mvd* and *Mvk* are all transcriptionally regulated by SREBP2, a master regulator of lipid and cholesterol synthesis gene expression that itself is activated at low cholesterol levels (Lewis *et al*, 2011). SREBP2 activity itself is regulated by mTORC1 (Peterson *et al*, 2011; Lewis *et al*, 2011), hence, depends on the metabolic state of the cell. In our transcriptomics data, we find mTORC1 related pathways to be downregulated upon Tau overexpression, which indirectly suggests reduced SREBP-2 activity. It remains to be tested though whether and through which mechanisms Tau could regulate SREBP-2 activity. In any case, SERBP-2 metabolism could be a potential therapeutic target (Lewis *et al*, 2011) to battle cholesterol and metabolic changes in the AD brain.

Cholesterol is a central component of membrane biology and lipid-based signaling. Downregulation of cholesterol synthesis can be observed in response to changing metabolic needs or environmental cues. For example, during periods of energy scarcity, cells shift their focus to conserving energy, e.g., by breaking down stored lipids for energy production and downregulating cholesterol synthesis to prevent the expenditure of energy on the synthesis of non-essential molecules like cholesterol. A neuronal strategy of “saving” energy by reducing cholesterol production would be in line with the observation that somatodendritic Tau levels (= Tau mis-sorting) occurs in conditions of energy-demanding neuronal hyperexcitation (Askin *et al*, 2024; Zempel *et al*, 2010; Tai *et al*, 2016). In addition to saving energy needed for cholesterol production, reducing cholesterol biosynthesis would have an additional effect on the neuronal energy metabolism by reducing neuronal activity: A decrease in neuronal cholesterol reduces synaptic activity (Shin *et al*, 2024), e.g., by reducing synaptic vesicle release (Mailman *et al*, 2011; Wasser *et al*, 2007). We speculate that Tau missorting under excitatory stress, followed by Tau-induced reduction in cholesterol biosynthesis, could put a break on excitation to avoid further overexcitation. However, extended cholesterol restriction, i.e., because of pathologically prolonged Tau missorting, could lead to synaptic decline and neurotoxicity in AD and tauopathies (Koudinov & Koudinova, 2001; Shin *et al*, 2024). Notably, *Igf1* (insulin growth factor-1), another important regulator of cell metabolism, appeared to be upregulated at high Tau levels as well, suggesting that Tau missorting may be involved in a global adaptation of the neuronal metabolism. Further studies are needed to test this currently speculative feedback loop.

Neurons receive most of their cholesterol in form of cholesterol/APOE lipoparticles from astrocytes. However, cholesterol synthesis by astrocytes could also be influenced by Tau since astrocytes themselves – as well as oligodendrocytes - also express low amounts of Tau. A Tau-related reduction of cholesterol synthesis in oligodendrocytes would impact myelin sheets, and therefore disturb neuronal transmission (Li *et al*, 2022). In addition, neuronal cholesterol synthesis seems to be essential for re-myelination processes in the brain (Berghoff *et al*, 2021), which becomes important in neurodegenerative diseases like AD, where de-myelination, leading to network activity abnormalities and promoting and amyloid-beta deposition (Depp *et al*, 2023), occurs early on. A-beta induced Tau missorting, leading to a reduction of neuronal and glial cholesterol synthesis, could therefore accelerate AD-related myelin deterioration and A-beta deposition, thereby creating a fatal feed-back loop with long-reaching consequences for neuronal network function in the brain. In addition, low cholesterol levels also promote neuronal internalization of seeding-competent Tau oligomers (Tuck *et al*, 2022), which could contribute to an accelerated spread of Tau aggregation pathology in the brain. Notably, our data indicate that Hmgcr protein levels are indeed reduced in AD hippocampal neurons having missorted Tau, and brain cholesterols levels have been reported to be reduced in AD (REF). Whether reduced cholesterol levels in AD brains are – at least partially - a consequence of neuronal Tau missorting, and whether this provides the missing link between AD-related A-beta and Tau pathology, myelination defects, and network dysfunction remains to be clarified.

### Tau missorting - sufficient to induce AD-related neuronal stress response pathways?

Since somatodendritic missorting of Tau is considered to be upstream of Tau aggregation in AD (Luna-Muñoz *et al*, 2007), the herein investigated effects of high neuronal Tau levels should reflect initial, pre-pathological Tau changes in AD brains; they may also be interpreted as part of the physiological neuronal response to certain stress stimuli. Accordingly, we find that genes deregulated in response to Tau cluster in pathways of neuronal stress response that are also activated in AD and other neurodegenerative diseases (e.g., TNFα/NFKB signaling, apoptosis, p53 pathway). This includes, for example, the TFs *Rest, Crem*, and *Jun*, and the lncRNA *Neat1*. Furthermore, activation of stress response and pro-inflammatory signaling, including TNF-α/NFK-b signaling observed in neurons with high Tau levels, can induce the acute downregulation of biosynthetic pathways, including cholesterol production, to redirect resources away from non-essential processes like lipid synthesis. A direct link between Tau, neuronal stressors, and cholesterol synthesis still needs to be confirmed though.

Pathological conditions presenting with Tau missorting and aggregation in neuronal cell bodies - e.g., neuronal hyperexcitation in seizures or as a result of A-beta in AD, mechanical brain damage in traumatic brain injury (TBI) and chronic traumatic encephalitis (CTE) - often involve repeated or extreme versions of physiological, Tau-activating stimuli. In these scenarios, prolonged Tau missorting may exacerbate Tau’s function in neuronal stress response leading to neuronal silencing, synapse loss, and eventually neuronal death. This could explain why TF regulation and gene expression changes reported in AD are in part contrary to what we observe for neurons with elevated somatodendritic soluble Tau levels. In addition, the sequestration of soluble Tau into aggregates. i.e., NFTs in AD, could prevent its nuclear activity in regulating TFs and other genes, resulting in a discrepancy in gene expression data between neurons with soluble physiological Tau and AD brains. Accordingly, it was suggested that a loss of physiological nuclear Tau would lead to chromatin relaxation and alter transcription (Rico *et al*, 2018; Mansuroglu *et al*, 2016). Previously observed differences in the effect of Tau on gene expression in human neurons versus AD brains could be explained by this (Klein *et al*, 2019).

In conclusion, our data implicate a novel function of missorted Tau in physically modulating neuronal chromatin organization, preferentially at the NE and involving LBR and SUN1, to induce stress-related gene expression profiles and reduce cholesterol biosynthesis. Interactions of Tau with LBR and SUN1 (and potentially other NE proteins) seem also relevant for Tau-related NE distortions and nuclear abnormalities observed in AD and FTD brains. These findings propel the understanding of somatodendritic Tau missorting in both the physiological neuronal context and as early, pre-pathological change in AD.

## Expanded View Figure Legends

**Figure EV1. Additional data for AAV-mediated modulation of Tau levels in hippocampal neurons.**

(A) Representative images of primary mouse hippocampal neuron cultures (DIV12), in which Tau levels were modulated using AAVs added on DIV5. Total Tau is detected by immunofluorescence. AraC treatment was used to deplete glia cells in cultures. Scale bars = 20 µm.

(B) Schematics AAV ZFP72, a non-gene targeting control AAVs for the MAPT-silencing ZFP90. Scale bar = 50 µm

(C) Tau level variation in cell bodies (= soma) of hippocampal (CA1) neurons of AD brain. Levels were determined based on total Tau immunofluorescence in cell bodies outlined in the MAP2 channel and normalized to tissue background measured in neuropil. Data points represent individual cell bodies in the image shown, as indicated by the colored circles.

(D) Full western blots to determine Tau levels in neuronal lysates (related to Figure 2B).

**Figure EV2. Additional data for Tau and TDP-43ctf nuclear proximity labeling.**

(A) Full western blots (anti-HA, H3 and Biotin) for nuclear-enriched cell lysates (input), related to Figure 3D.

(B) Full western blots for nuclear-enriched cell lysates (input) and pulldown (Streptav.PD), related to Figure 3D.

(C) Domain structure of full-length TDP-43, comprised of an N-terminal domain harboring a nuclear localization signal (NLS), two RNA-binding domains (RRM1+2, with a nuclear export signal (NES) in RRM2), and a C-terminal domain. Aggregation-prone TDP-43ctf (C-terminal fragment, aa 208-414) is generated in the brain via autocleavage at amino acid R208. TurboID constructs and workflow for proximity biotinylation of potential nuclear TDP-43ctf interactors.

(D) Expression of TurboID-TDP-43ctf construct in SH-SY5Y neuroblastoma cells. Anti-HA staining (cyan) shows expression and cellular localization of HA-TurboID construct, successful biotinylation (magenta) is verified by anti-Streptavidin staining. TDP-43 staining was included (yellow). Scale bars = 10 µm.

(E) Volcano plots of proximity-biotinylated proteins associated with TurboID-TDP-43ctf compared to TurboID control. Significant differences in proteins between experimental groups and control were identified using a two-sided t-test. A log2[fold change] >1.5 and p-value <-log10[1] were used as threshold for significance. Selected interactors are indicated as orange data points and labeled with protein names. Data retrieved from 5 biological replicates.

(F) Pathway enrichment analysis (GO Cellular compartment) for proteins biotinylated by TurboID-TDP-43ctf.

(G) Venn diagram showing overlap of biotinylated proteins significantly enriched in TurboID-Tau, TurboID-Tau^P301L^, and TurboID-TDP-43ctf compared to TurboID control (created with https://bioinfogp.cnb.csic.es/tools/venny/)

(H) Pathway enrichment analysis (ORA, over-representation analysis, Gene ontology (GO), Biological Processes analysis of potential nuclear Tau (left), TauP301L (middle), and TDP-43ctf (right) interactors.

**Figure EV3. Additional data for Tau co-IPs and PLAs in cells and neurons.**

(A) Full western blot of endogenous Tau Co-IP with Nesprin, LaminA/C and LBR (related to Figure 4A).

(B) SH-SY5Y cells expressing GFP or GFP-Tau for co-IP of NE proteins using GFP-Trap beads.

(C) Full western blot of LBR Co-IP with GFP-Tau in different buffer conditions (related to Figure 4B).

(D) Full western blots of SUN1 co-IP with GFP or GFP-Tau in high salt buffer (related to Figure 4B).

(E) PLA of endogenous Tau with H4K20me3 or H3K9me3 in SH-SY5Y cells. PLA signal was quantified in nuclear ROIs. N = ∼300 nuclei. One-way ANOVA with Dunnett post-test. Scale bars = 10 µm.

(F) PLAs of endogenous Tau with heterochromatin markers in hippocampal neurons (DIV12). PLA signal was quantified in nuclear ROIs. Scale bars = 10 µm.

**Figure EV4. Additional data for human brain IHCs and PLAs.**

(A) Example low magnification overview images of human brain sections immunolabeled for MAP2, phospho-Tau (pS202/pT205/pT231 or pT205/pS199), and respective Tau interacting nuclear envelope protein (SUN1, LBR or HP1γ in hippocampal CA1; LEM-2 in cortex). Scale bars = 100 µm.

(B) Representative image of immunofluorescently labeled Tau interactor LEM-2 in cortical sections from human AD brain. Zoom-ins show selected neuronal nuclei/cell bodies (MAP2+) having different phospho-Tau (pS202/pT205/pT231) levels and aggregation states. Scale bar = 10 µm.

(C) “Cart wheel”-like DNA structure in neuronal nuclei of human hippocampus. Example images of AD hippocampal section labeled for phospho-Tau (p-Tau), MAP2, and DNA (DAPI). At higher magnification (white boxes; zooms show inverted DAPI signal), the cart wheel like DNA arrangement in the nuclei becomes visible. Quantification of Cart wheel-like neuronal DNA arrangement in AD vs. control brain hippocampi. Data shown as mean±SD, data points indicate 8-11 images analyzed from three 3 AD/Ctrl cases. Unpaired Student’s t-test. Scale bar = 20 µm. Bottom: Example image and radial DNA profile (clock scan) of neuronal cart wheel DNA arrangement.

(D) Immunostaining for MAP2 (cyan) and total Tau (pink) in combination with PLA (yellow) for endogenous Tau with LEM-2 in AD brain sections (hippocampal CA1 and dentate gyrus). White squares indicate location of zoom-ins shown below for combined IHC+PLA and PLA with DAPI. Scale bars = 10 µm.

**Figure EV5. TauP301L induced cytoplasm-filled nuclear NE invaginations**

(A) HEK293 Cells expressing eGFP–TauP301L alone, or mRFP–FKBP–TauP301L co-expressed with LBR–eCFP–SNAP or SUN1–eCFP–SNAP in the absence or presence of rCD1, are shown. Normalized fluorescence intensity line plots along white lines in merged images show spatial distribution of Tau, LBR or SUN1, and DNA. Scale bar = 10 µm.

(B) Co-localization of TauP301L nuclear inclusions with nuclear organelle markers. Cells co-expressing mRFP–FKBP–TauP301L and either LBR–eCFP–SNAP (top) or SUN1–eCFP–SNAP (bottom) were immunostained for the nucleolar NPM1, nuclear speckle protein SRRM2, or chromatin binding HP1γ. White boxes indicate location of enlarged regions shown on the right. Normalized intensity line plots are generated along dashed white lines. Note, HP1γ is enriched around inclusions where DNA (see panel A) is condensed as well. Scale bar = 10 µm

**Figure EV6. Additional data for LBR-npd condensation and DNA binding.**

(A) Disorder (PONDR > 0.5) and LLPS propensity (CatGranule > 0.75) of identified (potential) nuclear Tau interactors.

(B) PhaSePred meta-prediction (10-feature model) for potential nuclear Tau interactors.

(C) Representative overview images showing LBR-npd coating of λ-DNA in the absence and presence of HP1γ and/or Tau. Scale bars = 10 µm.

(D) LBR-npd mobile fractions in the absence and presence of λ-DNA, determined from last 10 data points of average FRAP curves in Figure 5K. One-way ANOVA with Tukey post-test.

**Figure EV7. Heterochromatin analysis in neurons with different Tau levels.**

(A) Workflow of H3K9me3 chromocenter analysis in neuronal nuclei. Projections of z-stacks covering the entire nucleus were analyzes using CellProfiler: Primary objects (Nuclei from DAPI channel) were identified, position and size of chromocenters detected in H3K9me3 channel applying a size threshold (5-10 pixels).

(B) Quantification of H3K9me3 chromocenter number in neurons with different Tau levels. N = number of nuclei (indicated), 3 biological replicates, and 3 images analyzed per condition. One-way ANOVA with Dunnett post-test.

(C) Western blot of heterochromatin markers shows increased repressive H3K9me3 levels in Tau OE and TauPL OE neurons. H3K9ac changes are not significant. n = 3 experiments. One-way ANOVA with Dunnett post-test.

(D) Full blots for data analyzed in S7C.

**Figure EV8. Neuronal cultures and PCA of RNAseq samples.**

(A) Immunostaining shows reduction of astrocytes/glia cells without impairing neuronal integrity after treatment with AraC (1 µM, 8 h). Scale bar = 50 µm.

(B) Principal component analysis (PCA) of RNAseq data per condition (Tau base, Tau OE, Tau KD).

(C) Adjusted R^2^ values from linear regression for each batch used for RNAseq.

(D) Batched-corrected gene expression of *MAPT* across the conditions.

**Figure EV9. Additional analysis of DEGs in neurons with high and low Tau levels.**

(A) Number of differentially expressed genes (DEGs) by gene classes for Tau OE versus Tau base, Tau KD versus Tau base, and Tau OE versus Tau KD neurons.

(B) Gene set enrichment analysis (GSEA) of all DEGs in Tau o/e versus Tau base and Tau OE vs. Tau KD neurons using GO Biological processes.

(C) Non-protein coding DEGs in Tau OE versus Tau base, Tau OE versus Tau base, and Tau OE vs. Tau KD neurons organized by condition and classes.

**Figure EV10. Transcription factor (TF) analysis**

(A) UniProt keyword enrichment analysis of transcription factors. Bubble plot showing significantly enriched UniProt annotated keywords identified by STRING analysis of transcription factors differentially regulated in the RNAseq dataset of Tau OE vs Tau base (see Figure 8G). Enriched categories include Activator, Repressor, DNA-binding, and Transcription regulation. The x-axis indicates the enrichment signal. Bubble size represents the number of genes associated with each keyword, and color denotes the false discovery rate (FDR).

(B) Table summarizing transcription factors (TFs) identified among significant DEGs in Tau OE vs Tau base neurons, obtained by comparison with the AnimalTFDB v4.0 database. For each TF, chromosome location, regulatory activity (activator or repressor), direction of regulation (up-or downregulated), log2(fold change, FC), and p-value are indicated. Functional clustering of transcription factors into activators and repressors was derived from STRING protein–protein interaction analysis.

(C) Transcription factor (TF) analysis (wPGSA) of RNA sequencing data considering all gene expression data except transposable elements. Positive wPSGA t-score of a TF indicates overall upregulation of its target genes, a negative wPSGA t-score an overall down-regulation of the target genes. Threshold of significance was set as [wPSGA t-score] >4 and adjusted p-value < 10^-5^. Nomenclature from wPGSA analysis was used.

**Figure EV11. CUT&Tag for H3K9ac.**

(A) Experimental outline for CUT&Tag.

(B) Heat map of significant enriched H3K9ac loci in different genomic regions, clustered by condition.

**Figure EV12. LAD type 1 association of DEGs in neurons with high Tau levels.**

DEG association with LAD type1 and chromosomes for Tau OE vs. Tau base (top) or Tau OE vs. Tau KD (bottom) shows no significant enrichment of DEGs in LAD type1.

## Material and Methods

### Human samples

#### DAPI and Tau imaging

Images of AD and control brain array tomography sections stained for misfolded Tau (Alz50) and DAPI were provided by Prof. Tara Spires-Jones (Centre for Discovery Brain Science, University of Edinburgh). The use of human tissue for these post-mortem studies was reviewed and approved by the Edinburgh Brain Bank ethics committee and the medical research ethics committee. Post-mortem brains of 61-to 96-year-old (24 female, 41 male) AD patients (Braak V-VI, n=40) and healthy controls (n=25) were used (Supplemental Table T10).

#### IHC and PLA

Post-mortem, FFPE-fixed AD and control brain samples for IHC and PLA were received from the Biobank of the Department of Neuropathology, Charité-Universitätsmedizin Berlin, Germany (Supplemental Table T11). All experiments performed were approved by the local ethics committee at the Charité Berlin, patient consensus to the use of the tissue was given to the biobanks the tissue was received from.

### Animals

All experiments involving mice were carried out according to the guidelines stated in the directive 2010/63/EU of the European Parliament on the protection of animals used for scientific purposes and were approved by local authorities in Berlin and the animal welfare committee of the Charité university in Berlin, Germany.

### Plasmid constructs

#### Mammalian expression plasmids

Tau (human full-length wild-type Tau, 2N4R isoform), FTD-mutant Tau^P301L^, and TDP-43ctf (aa208-414) were cloned into 3xHA-TurboID-NLS_pCDNA3 plasmid (Addgene #107171) under a CMV promotor. The TurboID-NLS_pCDNA3 backbone was cut with EcoRI/XhoI to remove the NLS sequence and enable cytoplasmic expression and localization. Tau, Tau^P301L^ or TDP-43ctf inserts were PCR-amplified from different sources (key resource table) and inserted into the backbone using Gibson Assembly cloning. A flexible linker [(SGGG)]3 was inserted between TurboID and the fusion partners to enable correct protein folding. 3xHA-TurboID was cloned from 3xHA-TurboID-NLS_pCDNA3 plasmid (Addgene #107171) using EcoRI to remove the NLS sequence, followed by blunting using T4 polymerase and ligation using T4 ligase.

The eGFP-Tau (human, 2N4R) sequence was subcloned from a pRK5 backbone (kindly provided by Eckhard Mandelkow) into an AAV vector with CAG promotor. GFP-Tau^P301L^ and GFP-Tau^P301L/S320F^ were generated by site directed mutagenesis. Tau-GFP-NLS was generated by fusing GFP-3xNLS (three tandem repeats of a classical SV40 large T-antigen nuclear localization signal (DPKKKRKV)\₃) C-terminally to full-length human Tau (2N4R), separated by a 3xGSSS linker. Tau^P301L^ and Tau^P301L/S320F^ variants were generated by site directed mutagenesis.

For iRFP670-Tau or iRFP670-Tau^P301L^, human full-length Tau (2N4R) or Tau^P301L^ were fused to PCR amplified iRFP670 (Addgene #45466) and inserted into a pRK5-EGFP backbone with CMV promotor (Addgene #46904) using Gibson Assembly cloning.

For mRFP-FKBP-Tau or mRFP-FKBP-Tau^P301L^, human full-length wild-type Tau (2N4R) or Tau^P301L^ were cloned into a pmRFP-FKBP-PRPF38A backbone carrying a CMV promotor (kindly provided by Florian Heyd, FU Berlin). SUN1 (aa1-916) and full-length LBR were fused to eCFP and SNAP.

#### AAV constructs

Plasmids for eGFP, eGFP-Tau, eGFP-Tau^P301L^, eGFP.p2a.ZFP90, and eGFP.p2a.ZFP72 were sub-cloned into an AAV backbone using standard cloning procedures. Genes for ZFP90 and ZFP70 were synthesized based on their published protein sequence (Wegmann *et al*, 2021). AAV particles (AAV 2/9 PhP.B) were produced by the Charité viral core facility (VCF).

### Cell culture and transfection

Human SH-SY5Y cells obtained from ATCC (CRL-2266) were cultured in DMEM (Life Technologies) containing 10% FBS, 1% Pen/Strep (Sigma-Aldrich) and 2 mM L-Glutamine (ThermoFisher) and maintained at 37°C and 5% CO2. Cells were grown in 10 cm dishes (60-70% confluency) for proximity biotinylation pulldowns as well as co-immunoprecipitations (co-IPs)/WBs and in µ-slide 8-well imaging dishes (ibidi; ibiTreat) for proximity ligation assays (PLA) and immunofluorescence (IF). Cells were transfected with Lipofectamine 2000 (Life Technologies) according to the manufacturer’s protocol.

Human HEK-293 cells obtained from ATCC (CRL-1573) were cultured in DMEM (Life Technologies) containing 5% FBS and 1% Pen/Strep (Sigma-Aldrich) and maintained at 37°C and 5% CO2. Cells were passaged in 0.05% Trypsin-EDTA (Life Technologies) and plated in µ-slide 8-well imaging dishes (ibidi; ibiTreat) for immunofluorescence (IF) and for co-transfection of Tau with LBR/Sun1.

### Primary hippocampal neurons

*Primary mouse neurons*were prepared from hippocampi dissected from P0-P1 wildtype mice (C57/B6) of either sex. Hippocampi were dissected in ice-cold HBSS (Millipore) containing 1% P/S. Hippocampal tissue was digested in enzyme solution (DMEM, ThermoFisher Scientific) with 3.3 mM Cystein, 2 mM CaCl_2_, 1 mM EDTA and papain (20U/ml, Worthington) at 37°C for 30 min. Papain reaction was inhibited by incubating digested hippocampi in DMEM containing 10% FBS (ThermoFisher Scientific), 1% P/S, 38 mM BSA, and 95 mM trypsin inhibitor at 37°C for 5 min. Cells were triturated in complete Neurobasal-A (NBA) medium containing 10% FBS, 2% B-27, 1% Glutamax, 1% penicillin/streptomycin (P/S) and seeded in PLL (0.1 mg/mL) coated glass bottom µ-slide 8-well imaging dishes (ibidi) at a seeding density of ∼30’000 cells/cm^2^ (for IF/PLA experiments). For Neuronal nuclear levels of Tau, Cut&Tag and RNA sequencing experiments, cells were seeded into PLL-coated 6-well dishes (300’000 cells/dish). Every second day, one-fifth of the media was replaced by fresh complete NBA medium.

#### AraC treatment

To inhibit glial proliferation and obtain mostly neuronal cultures, seeded cells were subjected to a treatment with cytosine arabinoside (AraC) on DIV2. For that, 50% of the medium was removed and kept at 4°C as conditioned medium. Cells were treated with 1 µM (f.c.) AraC for 8 h at 37°C. Then, 80% of the medium containing AraC was removed and conditioned and fresh medium added to obtain the final volume. Success of AraC treatment was exemplary verified by immunostaining with anti-GFAP, anti-MAP2, and anti-Tau antibodies, showing almost full removal of astrocytes without obvious neurotoxicity.

#### AAV transduction

AAV particles for Tau expression (AAV CAG.eGFP-Tau, AAV CAG.eGFP-Tau^P301L^, AAV CAG.eGFP-Tau^P301L/S320F^), GFP expression (AAV CAG.eGFP), knockdown of endogenous mouse Tau (AAV hSyn.ZFP90-p2A-eGFP and control AAV hSyn.ZFP72-p2A-eGFP) were added directly into the medium of cultured hippocampal neurons on DIV5, excess particles removed via culture medium change on DIV7, and expression was allowed until DIV12.

### Neuronal nuclear levels of Tau

Hippocampal mouse neurons were seeded in PLL-coated 6-well plates, AraC-treated on DIV2, transduced with AAVs as needed on DIV5. On DIV 12, cells were washed once with PBS and scraped in hypotonic buffer (250 mM sucrose, 10 mM KCl, 2 mM CaCl2, 1.5 mM MgCl2, 20 mM HEPES pH 7.4, supplemented with protease and phosphatase inhibitors). Cells were lyzed by passing the cell suspension 10-20 times through a 25-26-gauge needle. The cytosolic fraction was obtained as supernatant from centrifugation for 5 min at 1,000 g. For the nuclear fraction, the pellet was washed 1-2 times in hypotonic buffer and centrifugation for 10 min at 1,000 g. The final nuclei pellet was resuspended in RIPA supplemented with 1.25 mM MgCl2 and protease and phosphatase inhibitor, and then treated with nuclease (1 units/25 µl) for 10 min at 4 °C and collected as the supernatant of a final centrifugation for 10 min at 10,000 g. Protein samples were mixed with 6x reducing SDS-containing loading buffer, boiled at 95°C for 5 min, and separated by SDS–PAGE (NuPAGE 4-12% Bis-Tris, Invitrogen).

### Streptavidin pulldown of biotinylated proteins from nuclear-enriched fractions

SH-SY5Y cells seeded in 10 cm dishes were lipotransfected (0.8% lipofectamine 2000, Life Technologies) with expression plasmids for 3xHA-TurboID-Tau, 3xHA-TurboID-Tau^P301L^, 3xHA-TurboID-TDP-43ctf (10 µg DNA) or 3xHA-TurboID control (5 µg DNA) to achieve similar expression levels. After 15 h, Biotin (50 μM f.c.; Sigma) was added to the medium for 10 min, before cells were washed once with ice-cold phosphate-buffered saline (1xPBS) to stop the biotinylation reaction. Cells were immediately subjected to a cellular fractionation protocol to obtain nuclear-enriched fractions. For this, cells were scraped in ice-cold hypotonic buffer (250 mM sucrose, 10 mM KCl, 2 mM CaCl_2_, 1.5 mM MgCl_2_, 20 mM HEPES pH 7.4; supplemented with protease and phosphatase inhibitors (ThermoFisher Scientific); 0.5 ml/10 cm dish) and kept on ice for 10 min. Cell lysis was achieved by passing the cells four-times through a 25/26-gauge needle (Omnifix). The crude nuclear fraction (nuclei and some cell debris) was obtained as a pellet from a 1’000 g spin for 5 min. The nuclear pellet was washed once in hypotonic buffer (0.5 ml/sample), passed 5x through a 25/26-gauge and subsequently centrifugated again at 1’000 g for 10 min. After removal of the supernatant, the nuclei pellet was resuspended in 50 µl Urea buffer (8 M urea, 50 mM Tris-HCl pH 7.5; supplemented with 1.25 mM MgCl_2_ and DNAse-I (ThermoFisher Scientific), and protease and phosphatase inhibitor), mixed for 20 min on a rotating wheel, and sonicated three times with 2 s-pulses. Nuclear lysates were centrifuged at 20000 g at 4°C for 15 min and the supernatant transferred into a fresh Eppendorf tube. Total protein concentration of nuclear-enriched fractions was measured by BCA assay, and the lysates stored at-80°C until further use.

For the pulldown of biotinylated proteins, streptavidin agarose beads (GE Healthcare) were used in combination with spin columns with inserted 10 µm filter (MoBiTec). Streptavidin-agarose bead slurry (20 µl) was added to each column, washed once with 300 µl PBS, and centrifuged for 1 min at 2’000 rpm. To block unspecific binding, beads were incubated with 300 µl 1% BSA in PBS for 1 h at room temperature under rotation. Meanwhile, all sample protein concentrations were adjusted by dilution in urea buffer to a volume of 400 µl and 1% of input samples was withdrawn and saved for analysis by western blot. For protein pulldown, blocked beads were incubated with nuclear-enriched fractions over night at 4°C under rotation. Columns were centrifuged to separate flow-through from the pulled-down biotinylated proteins, beads were washed 3-times with 300 µl wash buffer (2 M urea buffer) and once with detergent-free PBS. Finally, beads were resuspended in PBS supplemented with protease and phosphatase inhibitor (50 µl/sample). 20% of the samples were used for immunoblotting, the remaining 80% were kept at-20°C until analysis by mass spectrometry. Five independent biological replicates (proteins extracted from five different cell cultures at 5 different days) were performed for mass spectrometry.

### Mass spectrometry

#### In solution tryptic digest

First, proteins in samples were denatured in 8 M urea in 50 mM TEAB, pH 8.5, then samples were reduced with 5 mM DTT at 37°C for 60 min. This was followed by alkylation with 40 mM chloroacetamide at room temperature for 30 min in the dark. Protein digestion was carried out using Lysyl endopeptidase C (Wako) in a first step at an enzyme-to-protein ratio of 1:150 (w/w) at 37°C for 4 h. The sample was diluted using 50 mM TEAB, pH 8.0 to 2 M urea. The second digestion step was performed with trypsin (Serva) at an enzyme-to-protein ratio of 1:100 (w/w) at 37°C overnight. Samples were desalted with C8 material (Waters) in 300 µl pipette tips, dried under vacuum, and stored at-20°C until further use.

#### LC-MS analysis

LC-MS analysis was performed using an UltiMate 3000 RSLC nano LC system coupled online to an Orbitrap Fusion mass spectrometer (Thermo Fisher Scientific) with instrument software version 3.4. Proteins were separated by an in-house packaged 50 cm analytical column (Material: Poroshell 120 EC-C18, 2.7 μm, Agilent Technologies) with 120 min gradients of linearly increasing ACN concentration. Data acquisition for proteomics was performed in positive mode using a MS1(orbitrap)-MS2(ion trap) method. The following parameters were applied: MS1 resolution 120,000; scan range 375 – 1500 m/z; 400k AGC target; 50 ms max. injection time; MS2 scan rate rapid; first mass 110 m/z; 10k AGC target, 35 ms max. injection time; charge state 2-4 enable for MS2; HCD energy 30%; dynamic exclusion duration: 40 s.

#### Data analysis

LC-MS data were searched using MaxQuant (v.1.6.2.6a) with default parameters against the human proteome (retrieved from Uniprot, 2020, reviewed sequences only). A mass tolerance of 10 ppm was set for precursors on MS1 level, 0.5 Da for fragments on MS2 level. Methionine oxidation and N-terminal acetylation were set as variable, carbamidomethylation of cysteines was set as fixed modification. Proteins were quantified using label free quantification (LFQ) within MaxQuant. Statistical data from five replicates were processed using Perseus (v.1.6.7.0) with default parameters for imputation. For volcano plots and further analysis, a log2-fold change of 1.5 and p-value of 1 were used as threshold for significance. Pathway enrichment analysis was performed using WEB-based Gene Set Analysis Toolkit (https://www.webgestalt.org/) using over-representation (ORA) analysis as enrichment method and geneontology (GO)/cellular component as enrichment categories (default background). The Venn diagram for the representation of overlapping hits was produced using https://csbg.cnb.csic.es/BioinfoGP/venny.html. Cluster analysis was performed using only significantly enriched interactors (log2 fold change > 1.5 and a −log₁₀(p-value) > 1), using the k-means clustering algorithm implemented in STRING (https://string-db.org/).

### SDS-PAGE and Western blots

Protein samples were mixed with 6x reducing SDS-containing loading buffer, boiled at 95°C for 5 min, and separated by SDS–PAGE (NuPAGE 4-12% Bis-Tris, Invitrogen). Following electrophoresis, proteins were transferred to a nitrocellulose membrane, then the membrane was blocked with 3% BSA in PBS containing 0.05% Tween (PBS-T) at room temperature for 1 h. Membranes were incubated with primary antibodies diluted in blocking solution (PBS-T, 3% BSA), overnight at 4°C, washed 3-times for 5 min in PBS-T, and then incubated with fluorescent dye conjugated secondary antibodies diluted in blocking solution for 1 h at room temperature. Membranes were imaged using a Licor imaging system (Odyssey DLx). A complete list of antibodies used in Western blots can be found in the key resource table.

### Co-immunoprecipitation (co-IP) from nuclear-enriched fraction

SH-SY5Y cells were lipotransfected with eGFP-Tau or eGFP (Addgene #176015) plasmids (10 µg/10 cm dish) and allowed to express the constructs for 24 h. The nuclear-enriched fraction was prepared as described above via cellular fractionation with the following changes:

i) For co-IP experiments under high-salt conditions, HEPES buffer (20 mM HEPES pH 7.9, 400 mM NaCl, 1 mM EDTA, and 1 mM EGTA, containing protease/ phosphatase inhibitors) was used to resuspend the final nuclear pellet, followed by sonication 3-times for 3 sec. After nuclei lysis, GFP-trap magnetic particles M-270 (#gtd-20) were used for immunoprecipitation following the manufacturer protocol. In short, 25 µl beads slurry were used per sample and beads were equilibrated once with wash buffer (10 mM Tris/Cl pH 7.5, 150 mM NaCl, 0.05 % NP40, 0.5 mM EDTA; 0.5 ml/sample). For pulldown, nuclear lysates were adjusted to 0.5 ml in dilution Buffer (10 mM Tris/Cl pH 7.5, 150 mM NaCl, 0.5 mM EDTA, containing protease/phosphatase inhibitors) and incubated for 1 h at 4°C under rotation. Flow-through was separated from magnetic beads using a magnet, and beads were washed 3 times with wash Buffer. Protein elution was achieved by resuspending the beads in 80 µl educing 2x SDS sample buffer and boiling at 95°C for 5 min. Supernatants containing proteins were loaded on SDS-PAGE gels.

### Co-Immunoprecipitation (co-IP) of endogenous Tau

SH-SY5Y cell suspension was centrifuged for 5 min at 1000 g and 4°C. The cell pellet was resuspended in HEPES buffer (20 mM HEPES pH 7.9, 400 mM NaCl, 1 mM EDTA, and 1 mM EGTA, 0.5% NP-40, containing 25 U/µL nuclease, MgCl2, and protease/phosphatase inhibitors), followed by homogenization using 10-15 passes through a 26G needle. The cell suspension was kept 30 min on ice and sonicated 3-times for 3 s on ice. Samples were centrifuged at 10000x g for 20 min at 4°C. The supernatants were transferred to LowBind Eppendorf tube, and total protein was quantified in a Nanodrop. Protein G magnetic Dynabeads (Life Technologies, #10003D) were washed three times with PBS and blocked for 1 h at room temperature with 5% BSA in PBS at constant rotation. After blocking, beads were washed twice with PBS, and blocked beads (30 µL per antibody) incubated with 8 µg antibody in PBS containing 0.02% Tween-20 for 30 min at room temperature with constant rotation. The following antibodies were used: mouse anti-Nesprin-2 (Invitrogen, MA5-18075), mouse anti-Lamin A/C (Abcam, ab238303), mouse anti-LBR (Abcam, ab232731), and mouse IgG (Invitrogen) as a negative control. Beads were washed three times with PBS containing 0.02% Tween-20. For co-IP, lysate containing 250 µg of total protein was added to antibody-coupled beads in a final volume of 500 µl PBS and incubated overnight at 4°C at constant rotation. 5% of the lysate was retained for Western Blots (input). Next day, beads were collected using a magnetic rack, and the flow-through fraction was retained for Western Blot. Beads were washed three times with PBS. Proteins were eluted by boiling beads in 2xSDS sample buffer at 95°C for 5 min. Eluates, input (5%), and acetone-precipitated flow-through samples were analyzed by SDS-PAGE followed by immunoblotting using anti–total Tau antibody (DAKO). VeriBlot secondary antibody (1:1000) was used to minimize IgG heavy and light chain detection. Signal was detected by enhanced chemiluminescence (ECL).

### Immunocytochemistry (ICC) and immunohistochemistry (IHC)

Cells (SH-SY5Y, HEK-293T, or primary mouse hippocampal neurons, DIV12) were washed with PBS before fixation with 4% PFA in PBS for 15 min, followed by a wash with TBS for 10 min. After fixation, cells were permeabilized using 0.5% Triton-X100 in PBS for 20 min, blocked with 3% normal goat serum (NGS) in PBS for 1 h at room temperature, and incubated with primary antibodies diluted in 3% NGS/PBS over night at 4°C, or at room temperature for 2 h. After 3×10 min washes in PBS, cells were incubated with secondary antibodies in 3% NGS/PBS for 2 h at room temperature, then washed 3-times in PBS again before nuclei were counterstained with DAPI (1:1000 in PBS) for 15 min.

For IHC of human brain tissue, glass slide-mounted 3 µm-thick formalin-fixed, paraffin-embedded human hippocampus and temporal cortex sections were deparaffinized and baked in citrate buffer for antigen retrieval using Ventana Benchmark GX Staining System according to manufacturer protocol, followed by 3×10 min washes in PBS. Sections were surrounded with a hydrophobic barrier, followed by permeabilization in PBS containing 0.5% Triton-X for 10 min and washing in PBS for 10 min. Sections were blocked at room temperature for 1 h in PBS containing 10% NGS and 0.2% Triton-X100 (150-200 µl/slide). Primary antibodies were diluted in blocking solution and applied to sections overnight at 4°C in a humidity chamber under gentle agitation (100-200 µl/slide, depending on tissue size). After 3×10 min washes in PBS at room temperature, sections were incubated secondary antibodies diluted in 10% NGS and 0.2% Triton-X100 (150-200 µl/slide) for 1 h at room temperature. Finally, tissue auto-fluorescence was reduced by immersing the sections in 0.1% Sudan Black B in 70% ethanol for 20 min at room temperature followed by washing in PBS. The sections were mounted with mounting medium containing DAPI (DAKO). A complete list of antibodies used for ICC/IHC can be found in the key resource table.

### Nuclear envelope circularity in neurons

Hippocampal mouse neurons were seeded in PLL-coated 8-well plates (ibidi), AraC-treated on DIV2, transduced with AAVs on DIV5. On DIV12, cells were fixed and immunolabeled for Lamin A/C. Nuclear circularity was analyzed by detecting nuclei in the Lamin A/C channel and analyzing their shape using standard Image J plugins.

### BAF analysis

Human HEK-293T cells were plated in μ-slide 8-well imaging dishes (ibidi; ibiTreat) and lipotransfected (0.8% lipofectamine 2000, Life Technologies) with plasmids expressing SNAP-BAF in combination with eGFP and eGFP-Tau variants. After 24 h, cells were treated with 0.1 µM of SNAP dye650 diluted in DMEM for 30 min at 37°, immediately fixed with 4% PFA/PBS, and processed for Emerin immunostaining.

### Proximity ligation assays (PLAs)

PFA fixed cells (SH-SY5Y or primary mouse hippocampal neurons, DIV12) or FPPE human brain tissue were permeabilized as described above and subsequently blocked (Cells: 3% BSA in PBS/0.1% Triton-X100; Tissue: Duolink blocking solution) for 1 h at room temperature. Proximity ligation assays were performed according to manufacturer protocol (Duolink®InSituDetection Reagents Red, Sigma-Aldrich). Cells/tissue were incubated with diluted primary antibodies (Cells: in 3%BSA, 0.1% Triton in PBS, 80µl per sample; Tissue: in Duolink antibody diluent, 120 µl per sample) for 1 h at 37°C or overnight at 4°C in a humidity chamber under gentle agitation. Unbound primary antibodies were removed by washing samples 3-times in wash buffer A (0.01 M Tris, 0.15 M NaCl, 0.05%Tween) under gentle agitation. Samples were incubated with PLA probes (Duolink®Anti-Rabbit PLUS and Anti-Mouse MINUS) diluted 1:5 in antibody diluent (80-120 µl/sample) for 1 h at 37°C in a humidity chamber, then washed 2-times for 5 min in wash buffer A under gentle agitation to remove unbound PLA probes. For the ligation reaction, samples were incubated with Duolink®Ligation reagent (Ligation stock 1:5 in antibody diluent plus ligase, 1:40; 80-120 µl/sample) for 30-60min at 37°C in a humidity chamber, followed by 2×5 min washes with wash buffer A. For signal amplification, Duolink®Amplification stock was diluted 1:5 in high-purity water, and polymerase (1:80) was added before slides were incubated in 80-120 µl/sample for 120 min at 37°C in a humidity chamber. Final washing was performed twice for 10 min in 1X wash buffer B (0.2 M Tris, 0.1 M NaCl) and once for 1 min 0.01 X wash buffer B. Cells in 8-well chambers were incubated with DAPI in PBS (1:1000; D9542, Sigma) for 15 min at room temperature. Human tissue was additionally counterstained after PLA reaction by incubating the sections with primary antibody diluted in 10% NGS/PBS overnight at 4 °C under gentle agitation, washing 10 min with PBS at room temperature, and incubating with secondary antibody in 10% NGS/PBS for 1 h at room temperature. To reduce tissue autofluorescence, sections were immersed in 0.1% Sudan Black B in 70% ethanol for 20 min at room temperature and washed with PBS. Brain sections were mounted with mounting medium containing DAPI (DAKO). A complete list of antibodies used for ICC/IHC and PLA can be found in the key resource table. Imaging of PLAs was performed on a scanning confocal microscope (Nikon Spinning Disk Confocal CSU-X) using a 60x oil objective.

### Array tomography of human brains

For high resolution imaging of human nuclei, tissue from inferior temporal gyrus, BA20/21 from 40 AD and 25 control cases was collected at autopsy, fixed for 2 hours in 4% paraformaldehyde, dehydrated and embedded in LR white resin(Kay *et al*, 2013). Per case two blocks of tissues from Brodmann area 20/21 (BA20/21) were cut, stained and imaged as confocal z-stacks, and then analyzed blinded for case ID and Tau channel. Ribbons of 70 nm ultra-thin sections were cut on a Leica Ultracut and stained with Alz50 (courtesy Peter Davies) and DAPI. Images were acquired at 63x magnification on a Zeiss AxioImager and aligned with custom macros (https://github.com/Spires-Jones-Lab/Array_tomography_analysis_tool). Each stack contained 12 to 24 planes. For each case, at least three stacks from different sections with and without NFT pathology were analyzed.

### Recombinant proteins

All plasmids were verified by Sanger sequencing prior to protein production. Final protein concentrations were measured by BCA assays (Pierce) and the proteins were stored at−80°C.

#### Tau

Human full-length wild-type Tau (2N4R) was expressed in *E. coli* BL21 Star (DE3) (Invitrogen) and as previously described (Barghorn *et al*, 2005). Protein expression was induced with 0.5 mM IPTG at OD600 = 0.6 for ∼3 h at 37°C. Cells were harvested, resuspended in lysis buffer (20 mM MES, 1 mM EGTA, 0.2 mM MgCl_2_, 1 mM PMSF, 5 mM DTT, protease Inhibitors (Pierce Protease Inhibitor Mini Tablets, EDTA-free) and lysed using a French press. After initial purification by adding 500 mM NaCl and boiling at 95°C for 20 min, cell debris was removed by centrifugation (50’000 g, 40 min) and the supernatant was dialyzed against low salt buffer (Buffer A: 20 mM MES, 50 mM NaCl, 1 mM MgCl_2_, 1 mM EGTA, 2 mM DTT, 0.1 mM PMSF, pH 6.8), filtered (0.22 µm membrane filter), run through a cation exchange column (HiTrap SP HP,5 ml, GE Healthcare), and eluted with a high salt buffer (Buffer B: 20 mM MES, 1000 mM NaCl, 1 mM MgCl_2_, 1 mM EGTA, 2 mM DTT, 0.1 mM PMSF, pH 6.8). Fractions containing Tau were pooled, concentrated using spin column concentrators (Protein concentrators; 10–30 kDa MWCO, Pierce), and run through a size exclusion column (Superose 6 10/300, GE Healthcare). Fractions containing purified monomeric Tau were concentrated as before and buffer exchanged to PBS, 1 mM DTT, pH 7.4.

LBR-npd: For His-tagged LBR-NPD, the nucleoplasmic domain of human Lamin B receptor (LBR-npd; aa 1-211) was codon-optimized, and the gene synthesized by VectorBuilder and provided to us in a pET vector for protein expression (pET.LBR-npd(1-211)-TEV-10xHis). Protein expression (E. coli BL21 Star (DE3) (Invitrogen)) was induced with 0.5 mM IPTG at OD600 = 0.6 for 2 h at 37°C. Cells were harvested (6’000 rpm, 20 min), resuspended in lysis buffer (10 mM Tris-HCL, pH 7.5, 500 mM NaCl, 35 mM Imidazole, 1 mM DTT, protease Inhibitors (Pierce Protease Inhibitor Mini Tablets, EDTA-free) and lyzed using a french press. Afterwards, 10 µl Benzonase and 5 mg Lysozyme (Sigma-Aldrich) was added and incubated for 1 h at 4°C under rotation. After centrifugation (1 h at 35’000 g at 4°C), the supernatant of the lysate was filtered through a 0.22 µm membrane filter and loaded onto a Ni-NTA column (HiTrap, 5 ml, GE Healthcare) using a low imidazole buffer (Buffer 1: 10 mM Tri-HCl, pH 7.5 - 8.0, 500 mM NaCl, 35 mM Imidazole, 1 mM DTT), and eluted with a high imidazole buffer (Buffer 2: with 500 mM Imidazole). Fractions containing the protein were pooled, concentrated using spin column concentrators (Amicon Ultra 15 concentrators; 10kDa MWCO, Millipore), and buffer exchanged, using 25 mM HEPES, 500 mM NaCl, 1 mM DTT as final storage buffer.

#### HP1γ

GST-tagged HP1γ (Addgene: #24076) was expressed in *E. coli* BL21 Star (DE3) (Invitrogen). Protein expression was induced with 0.5 mM IPTG at OD600 = 0.6 for 2 h at 37°C. Cells were harvested (6’000 rpm, 20min), resuspended in lysis buffer (25 mM HEPES-KOH, pH 7.9, 0.1 mM EDTA, 12.5 mM MgCl_2_, 20 % glycerol, 0.1 mM KCl, 0.1% NP-40 Alternative (Millipore), 1 mM DTT, protease Inhibitors (Pierce Protease Inhibitor Mini Tablets, EDTA-free) and lysed using a French press. Afterwards, 10 µl Benzonase and 5 mg Lysozyme (Sigma-Aldrich) was added and incubated for 1 h at 4°C under rotation. After centrifugation (1 h, 30’000g, at 4°C) the supernatant of the lysate was filtered through a 0.22 µm membrane filter. The purification was carried out using Glutathion-Sepharos 4B (GE Healthcare) beads, the supernatant was incubated with the beads overnight at 4°C with end-to-end rotation. After incubation the beads were washed with lysis buffer (25 mM HEPES-KOH, pH 7.9, 0.1 mM EDTA, 12.5 mM MgCl_2_, 20% glycerol, 0.1 mM KCl, 0.1% NP-40 Alternative (Millipore), 1 mM DTT, protease Inhibitors (Pierce Protease Inhibitor Mini Tablets, EDTA-free) and the protein was eluted using 50 mM Tris, 200 mM NaCl, 20 mM reduced glutathione, 0.3 mM TCEP buffer. Fractions containing the protein were pooled, concentrated using spin column concentrators (Amicon Ultra 15 concentrators; 30 kDa MWCO, Millipore). The GST tag was cleaved using Glutathion-Sepharose 4B (GE Healthcare) beads and thrombin protease (Sigma-Aldrich). After overnight incubation with protease, the supernatant was run through a size exclusion column (Superose 6 10/300, GE Healthcare) and buffer exchanged, using PBS containing 5% glycerol, 1 mM DTT as final storage buffer.

### Fluorescent labeling of proteins

Recombinant proteins were fluorescently labeled using amine-reactive DyLight-NHS ester dyes (Thermo Scientific; Tau: DyLight488/DyLight650; LBR-npd: DyLight650; HP1γ: DyLight550) following manufacturer instructions. The dye was dissolved in DMSO to a final concentration of 10 µg/µl, then added in 5-to 10-fold molar excess to the protein in PBS with 1 mM DTT, and the labeling reaction was performed for 2 h at room temperature at 250 rpm. Excess dye was removed by dialysis (Pur-A-Lyzer Mini Dialysis tubes, Sigma-Aldrich; MWCO 12–14 kDa) against PBS, pH 7.4, 1 mM DTT at 4°C. The labeling degree (amount of dye per molecule protein) was determined by measuring the final protein concentration and correlating it to the maximum absorbance of the attached dye. For calculations see manufacturer instructions.

### Protein condensation assays

#### LBR-npd

8 µM LBR-npd (1% LBP-npd-DyLight561) were imaged alone or in the presence of 5.4 µM HP1γ (containing 1% HP1γ-DyLight650), 4 µM Tau (containing 2% Tau-DyLight488), and/or 75 ng/µl λ-DNA in 20 mM HEPES, pH 7.4, 130 mM NaCl. Samples (3 µl) were pipetted onto amine-coated 35-mm dishes (Matek), equipped with ddH_2_O-soaked tissue to maintain humidity, and imaged after 1-2 hours.

LBR-npd condensates were imaged after 1 h on a spinning disk confocal microscope (Nikon Spinning Disk Confocal CSU-X) using a 60x oil immersion objective. DNA was visualized by adding DAPI (1:1000 diluted) to the assay mix. FRAP was assessed by bleaching individual condensates in grape-like condensates assemblies (in the absence of DNA) to 20-30% of their pre-bleach intensity, or by bleaching LBR-npd on DNA to ∼10% pre-bleach intensity.

### Tau-related NE distortions (membrane invaginations and inclusions)

HEK293 cells plated in µ-slide 8-well imaging dishes (ibidi; ibiTreat) were transfected using lipofectamine 2000 and expression allowed for 48h. Cells were treated with DMSO or 6 µM of the reversible chemical dimerizer 1 (rCD1) for 2.5 h before washing with 1XPBS followed by fixation with 4%PFA/PBS for 15 min at room temperature, and washing with TBS 3-times for 10 min. CMV-mRFP encoded in a pFUGW backbone was used as fluorophore only control (kindly provided by Craig Garner). Immunostaining was performed after permeabilization for NPM1, HP1γ, SRRM2, Lamin B1, Lamin A/C, Lamin B receptor (LBR), nuclear pore complex proteins, vimentin, GAPDH, and TUBA4A. Antibody details are listed in the Key Resources Table. Nuclei were stained with either 1:1000 siRDNA or 1:3000 DAPI dilutions for 10 minutes at room temperature. Invaginations and inclusions were counted manually, blinded for condition.

### Electrophoretic mobility shift assay (EMSA)

To analyze binding of recombinant LBR-npd and Tau to DNA, electrophoretic mobility shift assays (EMSAs) were performed using an EMSA Kit (Cat# E33075) following manufacturer instructions. In brief, 20 ng DNA fragments (500 bp NoLimit DNA fragments Cat#SM1641) were incubated with recombinant protein at indicated amounts in 20 µl 5x binding buffer (50 mM Tris HCl, pH 7.4, 750 mM KCl, 0.5 mM DTT, 0.5 mM EDTA) for 30 min at 30°C. Samples were supplemented with 6x EMSA gel-loading solution and separated via non-denaturing PAGE on 6% DNA retardation gels 0.5x TBE gel running buffer at 110 V for 150 min. Afterwards, DNA in the gel was stained using SYBR® Green EMSA Nucleic Acid Gel Stain and visualized using a laser-based Typhoon scanner with parameters and filter sets to visualize 488 dyes. Band intensities at the end of the lanes was interpreted as unshifted DNA, all signal above as DNA bound to protein.

### RNA sequencing

Hippocampal neurons were seeded in PLL-coated 6-well plates, AraC-treated on DIV2, transduced with AAVs for Tau expression (AAV eGFP-Tau) or mouse Tau knockdown (AAV ZFP90.P2a.GFP) on DIV5. On DIV12, cells were washed once with warm PBS before they were scraped-off in Trizol (ThermoFisher). RNA isolation was carried out using a RNeasy Micro Kit (Qiagen) following the manufacturer’s protocol. RNA quantity and integrity were evaluated using the High Sensitivity RNA assay on the TapeStation 4200 system (Agilent). For RNA sequencing, mRNA libraries were constructed using the TruSeq Stranded mRNA Library Prep Kit (Illumina), with 500 ng of total RNA as the input material. Library quantification was conducted with the Qubit High Sensitivity dsDNA assay, and the size distribution of library fragments was assessed using the D1000 ScreenTape assay on the TapeStation 4200 system (Agilent). Sequencing was performed in paired-end mode (75 cycles) on a NovaSeq 6000 system (Illumina) using the NovaSeq S4 v1.5 (200 Cycles) chemistry. Raw sequencing data were demultiplexed using bcl2fastq2 v2.20. RNA seq was performed on 2 biological replicates with 4 technical replicates each.

#### RNA Seq analysis

For RNAseq data, fastq files were aligned to GRCm38 using STAR (Dobin *et al*, 2013) with the following settings to maximize repeat mapping (--outFilterMultimapNmax 100, --winAchnorMultimapNmax 100). The TETranscripts (Jin *et al*, 2015) package was used to generate a count table using gencode.vM25.basic.annotation.gtf and the prepared repeat masker file mm10_rmsk_TE.gtf (https://labshare.cshl.edu/shares/mhammelllab/www-data/TEtranscripts/TE_GTF/). The following analysis steps were performed in R (v4.3.0) and R Studio (v1.4.1717). The count table was re-annotated using biomaRt (v.2.56.1) (Durinck *et al*, 2009). Genes with less than 10 counts in at least 5 samples were excluded from the analysis resulting in 19,214 genes kept in the analysis. Normalization of the count matrix was computed with R/DESeq2 (v1.40.2) (Love *et al*, 2014a) and a variance stabilizing transformation applied using the DESeq2 vst function at default settings. A batch between experiment days was corrected with limma (v3.56.2) for the visualizations and included as covariate in the following analysis. Differential expression analysis based on the DESeq2 package was performed adjusting p-values according to independent hypothesis weighting from the IHW package (v1.28.0) (Ignatiadis *et al*, 2016) and applying apeglm shrinkage from the apeglm package (v1.22.1) (Zhu *et al*, 2019). DEGs were defined based on a fold change threshold >2 and an adjusted p-value threshold of <0.05. A gene set enrichment analysis (GSEA) was performed with the transformed data as the input using R/fgsea (v1.26.0) (Korotkevich *et al*, 2021), whereby the Gene Ontology (GO) (Ashburner *et al*, 2000; Carbon *et al*, 2021) and the Molecular Signature Database (MSigDB) (Subramanian *et al*, 2005; Liberzon *et al*, 2015) Hallmark gene set was used.

#### Transposable elements (TEs)

The technique used for the locus-specific TE analysis is based on the method established by Schwarz et al. (Schwarz *et al*, 2022), who modified the work process of SalmonTE (v1.9.0; “SalmonTE*: a modified version for locus-specific expression analysis of transposable elements.”HackMD, 2023.

https://hackmd.io/@fgaudilliere/B1nlcJ66o) to count TE on a locus level instead of on a family level (Jeong *et al*, 2018). In brief, a library of TEs was generated from the reference genome (GRCm38) using RepeatMasker (v4.0.9; http://www.repeatmasker.org) and transformed to a fasta file format with the *alignToFasta.sh* helper script. Short repeats and N-rich repeated sequences were excluded, and a salmonTE index was subsequently created. With the indices locus-specific quantification using the *SalmonTE.py* script was performed.

The DESeq2 analysis was done as above described with the following changes. TEs with fewer than 16 counts were filtered out, resulting in a final dataset of 212,571 TEs. For DE analysis the Benjamini-Hochberg method for multiple testing correction and the normal shrinkage estimator for fold change moderation was applied(Love *et al*, 2014b).

#### Transcriptional Factors (TFs)

Differentially expressed genes (DEGs) identified as upregulated or downregulated, were compared against a curated list of mouse (*Mus musculus*) transcription factors obtained from AnimalTFDB v4.0. This analysis yielded 17 transcription factors, which were subsequently subjected to network analysis using STRING, confirming their involvement in gene regulatory processes. Activator and repressor transcription factors were identified based on STRING functional annotations and clustering.

#### LADs

For the LAD association data from mid brain human neurons was used and converted to mouse(Shah *et al*, 2023). The overlap between DEGs and LADs was tested with R/GenomicRanges (v1.52.0) (Lawrence *et al*, 2013) and a hyper-geometric test was performed per chromosome followed by a Benjamini-Hochberg adjustment with R/stats (v4.3.0).

#### wPGSA analysis

To predict transcription factor (TF) changes upstream of altered gene expression, wPGSA (Kawakami *et al*, 2016) analysis was performed as previously reported(Sakurai *et al*, 2024). In brief, normalized count matrixes generated by R/DESeq2 were used, and count values of each perturbed sample (Tau o/e or Tau k/d) were normalized to the mean count value of control samples (Tau base) using R (v4.3.3) and R Studio (v2023.12.1+402). The count matrixes normalized to the mean control values were imputed into the wPGSA package (v1.1.0) to predict altered TF activity in the perturbed samples. Significantly altered transcription factors were defined on a t-value threshold of >4 and an adjusted p-value (FDH) threshold of <1*10^-5^. We looked for TFs that are included in the DEG list using R.

### qPCR

Hippocampal neurons were seeded in PLL-coated 6-well plates, AraC-treated on DIV2, transduced with AAVs for Tau expression (AAV eGFP-Tau) or mouse Tau knockdown (AAV ZFP90.P2a.GFP) on DIV5. On DIV12, the culture medium was removed, cells were gently washed once with warm PBS, and then lyzed directly on the plate by adding 65 μL of lysis buffer per well, followed by cell scraping and pipetting to ensure complete cell disruption. Total RNA was purified from cell lysates using High Pure RNA Isolation Kit (ROCHE, Cat. No. 11828665001), according to the manufacturer’s instructions with minor modifications. The purified RNA was used immediately for RNA quantification in a Nanodrop and stored at −80°C until further analysis. Complementary DNA (cDNA) was synthesized from total RNA using the Transcriptor High Fidelity cDNA Synthesis Kit (Roche, Cat. No. 05081955001) according to the manufacturer’s instructions, using random hexamer primers. The resulting cDNA was either used immediately for downstream applications, including quantitative PCR, or stored at−20°C until further use. Quantitative real-time PCR (qPCR) was performed using LightCycler® SYBR Green I Master (Roche, Cat. No. 04707516001) on a LightCycler® 480 Instrument (Roche). Reactions were carried out on 96-well plates. Each qPCR reaction had a final volume of 25 μL and contained 12.5 μL of 2× SYBR Green I Master, 8.5 μL nuclease-free water, 1 μL of each forward and reverse primer (10 μM), and 2 μL of cDNA template. Primers were obtained from OriGene, and primer accession numbers and sequences are provided in the Key Resources Table. The genes analyzed were *GAPDH*, *NSDHL*, *TGIF1*, and *HMGCS1*. All samples were run in technical duplicates. Thermal cycling was performed under the following conditions: an initial denaturation step at 95°C for 10 min, followed by 40 cycles of amplification consisting of 95°C for 30 s, 60°C for 20 s, and 72°C for 30 s. Melt curve was conducted with continuous fluorescence acquisition from 95°C to 60°C following a brief denaturation step at 95°C. The run concluded with a cooling step at 37°C for 1 s. Gene expression analysis was performed in the Light Cycler 480 Software (Roche). Primer efficiency was determined by standard curve analysis and incorporated into the relative quantification calculations using the Advanced Relative Quantification method. Amplification efficiencies ranged from 1.895 to 2.429 across all primer pairs. GAPDH was used as the endogenous reference gene. Relative expression levels for each target gene were normalized to non-transduced neurons and are presented as fold change.

### Cleavage under targets & tagmentation (CUT&Tag) assay

For the genome-wide analysis of H3K9ac histone mark in primary mouse neurons the CUT&Tag-IT assay kit (Active Motif) was used according to manufacturer’s instructions. Hippocampal neurons were seeded in PLL-coated 6-well plates, AraC-treated on DIV2, transduced with AAVs for Tau expression (AAV eGFP-Tau) ort mouse Tau knockdown (AAV ZFP90.P2a.GFP) on DIV5. On DIV12 cells were harvested from 6-Wells by scratching in medium, pooling 3 Wells per condition. Cells were bound to Concavalin A beads, followed by incubation with 1.5 µg primary antibody (H3K9ac) o/n at 4°C using an orbital mixer. Following, Guinea Pig Anti-Rabbit secondary antibody was added to bind to primary bound antibody. The following steps included i) binding of CUT&Tag-IT™Assembled pA-Tn5 Transposomes (protein A fused to Tn5 transposase enzyme that is pre-loaded with sequencing adapters and binds to secondary antibody), ii) Tagmentation (fragmentation and tagging with sequencing adaptors) and iii) DNA extraction according to the protocol. PCR amplification was performed using unique combinations of one i7 Indexed Primer and one i5 Indexed Primer per condition. The final step to obtain libraries for sequencing contained a clean-up using magnetic SPRI beads. After library generation, the quality was assessed using Agilent TapeStation, yielding mostly fragments below 500 bp. Libraries were sequenced in a paired end configuration using NextSeq2000, 10-15 million reads per sample at the BIH/MDC Genomics Technology Platform, Berlin.

#### Analysis

Demultiplexed reads were aligned to GRCm39 using bowtie2 (Langmead & Salzberg, 2012) with the parameters --end-to-end --very-sensitive --no-mixed --no-discordant –phred33-I 10-X 700, and sorted and indexed using samtools. Duplicate reads were removed using picard. Read fragments from aligned reads were prepared using bedtools bamtobed to extract read pairs on the same chromosome with fragments smaller than 1000bp. Fragment bed files were converted to bedgraphs suitable for SEACR (Meers *et al*, 2019) using bedtools genomecov-bg. SEACR was run on each prepared bedgraph file with 0.005 (top 0.05%) norm stringent as parameters. Peaks identified in each sample were merged into a single master peak file. A count matrix of counts from each sample over all peaks was generated using the getCounts from the chromVAR package (Schep *et al*, 2017). Differential peaks were tested using DESeq2 and annotated with genes and genomic region using annotatePeak with tss region (-3000, 3000) from the ChIPseeker library. Overlaps with Repeat elements and T1 and T2 LAD elements were detected using findOverlaps.

### Chromosome painting

Hippocampal neurons were seeded in PLL-coated 12-well plates equipped with glass coverslips, AraC-treated on DIV2, and transduced with AAVs on DIV5. None-transduced neurons were included as controls (Tau base). On DIV12, neurons were fixed in 4% paraformaldehyde (PFA) for 10 min at room temperature, washed for 3 min in 0.1 M Tris-HCl (pH 7.4), and subsequently rinsed twice for 5 min in PBS. Cells were permeabilized with 0.5% Triton X-100 in PBS for 15 min at room temperature, followed by equilibration in 20% glycerol in PBS for 1.5 h at room temperature. Coverslips with cells were subjected to five freeze–thaw cycles: each cycle consisted of the immersion in liquid nitrogen for 30 s, thawing on tissue until the frozen layer disappeared, and incubation in 20% glycerol/PBS for 3 min. Coverslips were then washed three times for 10 min in PBS. For DNA depurination and probe accessibility, cells were incubated in freshly made 0.1 N HCl for 5 min at room temperature, washed three times for 3 min in 2X saline-sodium citrate (SSC) buffer, and three times for 5 min in PBS. Coverslips were prehybridized in freshly made 50% v/v formamide (pH 7.0) in 2X SSC for 1 h on ice. Chromosome painting probes (Metasystem, see key resource table for each probe information) were applied to a cleaned glass slide (2–2.5 μl per ¼ coverslip), and coverslips were placed cell-side down on the probe and sealed with nail polish. Samples and probes were denatured simultaneously on a hotplate at 75°C (±1°C) for 2 min and hybridized overnight for at least 16 h in a humidified chamber at 37°C (±1°C). Post-hybridization washes were performed by carefully removing the nail polish and detaching the coverslips from the slide, followed by washing the coverslips in 0.4X SSC at 72°C (±1°C) for 2 min. Coverslips were then rinsed in 2X SSC with 0.05% Tween-20 at RT for 30 s, briefly rinsed in distilled water, and air-dried. Cells were counterstained with DAPI/antifade mounting medium (MetaSystems, D-0902-500-DA) by applying a 2 - 3 μl drop into a clean glass slide and placing the coverslips cell-side down on DAPI, allowing to incubate for 10 min at room temperature, and sealed with nail polish. Slides were stored at 4°C protected from light until imaging. Confocal images were acquired as serial z-stacks with a step size of 0.3 μm using a 60x objective at 3x digital zoom.

### Fluorescence microscopy

Fluorescent images were acquired with a Nikon scanning Confocal A1Rsi+ microscope with a DS-Q12 Nikon camera at the Advanced Medical BIOimaging (Ambio) Core Facility at the Charité, Berlin. To prevent crosstalk among the fluorophores, channels were scanned sequentially, starting from the highest excitation wavelength to the lowest. Fluorescence lifetime imaging microscopy (FLIM) of SiR-DNA in primary mouse neurons and NesprinTS in HEK293 cells was performed using a Leica Stellaris 8-Falcon-STED confocal microscope at the Systems Biology Imaging core at BIMSB/MDC, Berlin. FRAP for LBR and LEM2 condensates, as well as co-transfections with SUN1 or LBR and Tau in HEK-293 cells, were performed on a Nikon Spinning Disk Confocal CSU-W1 SoRa microscope, as well as in a Zeiss Axio Observer Laser Scanning Confocal Microscope at DZNE, Berlin. Epifluorescence images of immunolabeled human brain slices, primary hippocampal neurons for nuclear circularity and HEK cells for BAF analysis, were collected on a Keyence BZ-X system. All images were processed with ImageJ or CellProfiler.

| Image modality / Microscope | Point scanning confocal / Nikon A1Rsi+ microscope |
| --- | --- |
| Illumination /Wavelength Selection | Excitation: DAPI = 405 nm; GFP = 488 nm; mCherry = 561 nm; Cy5 = 647 nm |
|  | Emission filters/Bandwidth: DAPI = 450/50; GFP = 525/50; mCherry = 595/50; Cy5 = 700/75 |
| Optics | Magnification: 20x; Type: Plan Apo; Immersion: Air; NA: 0.8; WD (µm) 1,000 |
|  | Magnification: 40x; Type: Plan Fluor; Immersion: Oil; NA: 1.3; WD (µm) 200 |
|  | Magnification: 60x; Type: Plan Apo; Immersion: Oil; NA: 1.4; WD (µm) 130 |
| Image modality / Microscope | Inverted fluorescence-phase contrast microscope / Keyence BZ-X800 |
| Illumination /Wavelength Selection | Excitation: DAPI = 360/40 nm; GFP = 470/40 nm; TRITC = 545/25 nm; Cy5 = 620/60 nm |
|  | Emission filters/Bandwidth: DAPI = 460/50; GFP = 525/50; TRITC = 605/70; Cy5 = 700/75 |
| Optics | Magnification: 60x; Type: Plan Apo; Immersion: Oil; NA: 1.40; WD (μm): 130 |
| **Image modality / Microscope** | **Nikon SoRa Spinning Disk Confocal CSU-W1** |
| Illumination /Wavelength Selection | Excitation: DAPI = 405 nm; CFP = 455 nm; GFP = 488 nm; mCherry = 561 nm; Cy5 = 647 nm |
|  | Emission filters/Bandwidth: DAPI = 447/60; CFP = 472/30; GFP = 525/50; mCherry = 600/50; Cy5 = 685/40 |
| Optics | Magnification: 60x; Type: Plan Apo; Immersion: Oil; NA: 1.4; WD (μm) 130 |
| **Image modality / Microscope** | **Carl Zeiss Axio Observer Z1 Laser Scanning Confocal with iXon ultra (Andor) camera** |
| Illumination /Wavelength Selection | Excitation: CFP = 455 nm; RFP = 561 nm; Cy5 = 647 nm |
|  | Emission filters/Bandwidth: CFP = 480; RFP = 584; Cy5 = 670 nm |
| Optics | Magnification: 63x; Type: Plan Apo; Immersion: Oil; NA: 1.40; WD (μm): 512 |
| **Image modality / Microscope** | **Confocal Leica SP8 Stellaris** |
| Illumination /Wavelength Selection | Excitation: CFP = 450 nm; GFP = 480 nm; YFP/Venus = 510 nm; iRFP/SirDNA = 645 nm |
| Optics | Emission filters/Bandwidth: CFP/TFP1 = 500/50; GFP = 525/50; YFP/Venus = 555/50; iRFP/SirDNA = 700/75 |
| **Sample Preparation** |  |
|  | 8-well imaging dishes (ibiTreat; Polymer coverslip No. 1.5H) |
|  | 8-well glass bottom imaging dishes (Glass coverslip No. 1.5H) |
|  | 35 mm dishes (Glass coverslip No. 1.5H) |

### Image analysis

#### Heterochromatin markers

An image analysis pipeline was created in CellProfiler to analyze heterochromatin intensity and chromocenter number and size. The pipeline for nuclear intensity consisted of: IdentifyPrimaryObjects (in DAPI channel) > MeasureObjectIntensity (in DAPI and HC channel) > MeasureObjectSizeShape. The pipeline for chromocenter analysis consisted of: IdentifyPrimaryObjects (in DAPI channel) > EnhanceOrSuppressFeatures (Enhance: HC in heterochromatin channel) > MasImage > IdentifyPrimaryObjects > MeasureObjectSizeShape > MeasureObjectIntensityDistribution (Bin =4). > RelateObjects

#### Line plots

Define horizontal line of same length as ROI; ImageJ > Analyze > Plot profile.

#### Clock scan

For the chromatin and the chromosome territories analysis, the *Clock scan* plugin of ImageJ was used. This plugin (i) measures the fluorescence intensity profile along a line (radial line) from the center to the outer boundary of an ROI, and (ii) creates an average of all 360 intensity profiles measured upon 360° rotation of the radial line. The output is a graph presenting average fluorescence intensity (y-axis) over the distance to the center in % of radius (x-axis; 100% = nuclear border). For the analysis of the average radial distribution of chromatin, neuronal nuclei were analyzed in SiR-DNA images collected from living neurons. For the analysis of the averaged radial distribution of chromosome territories, chromosome signals were quantified in the equatorial plane of the z-stacks acquired from neuronal nuclei of fixed neurons. Clock scans of nuclei with fluorescence signal accumulation at the NE produced a graph with a peak of ∼90-100% radius.

#### Nuclear invaginations in human brain

Z-stacks of DAPI-stained nuclei in AD brain were obtained. Nuclei were identified as “invaginated” when at least one membrane fold (=invagination) was spotted across the z-stack. The presence of misfolded Tau (Alz50+) around nuclei was determined afterwards by merging DAPI and ALZ50 channels.

#### TurboID constructs

In representative confocal images (60x) of TurboID constructs expressed in SH-SY5Y cells, nuclei were outlined using the DAPI channel and added as white outlines using *Flatten* in ROI Manager.

#### PLAs

To quantify PLA signals, nuclei ROIs were defined by thresholding images in the DAPI channel (ImageJ) and defining them as individual particles. Afterwards, PLA signal spots within nuclear ROIs were detected in the PLA channel (Cy3) using the *Find maxima* function. The number of PLA signal spots were normalized to the number of cell nuclei. 2-3 experiments were performed for each condition, and 3-6 images analyzed per condition.

#### Protein condensates

Surface coverage was determined on an image-by-image base by measuring the area covered by condensates after thresholding the image to remove surface background in ImageJ. Co-partitioning coefficients (= c_dense phase_/c_light phase_) were determined on an image-by-image base by dividing the mean fluorescence intensities measured inside condensates by the one measured outside of condensates. Coefficient variance of DNA inside condensates was determined on a condensate-by-condensate base by dividing the standard deviation by the mean of individual condensates (CoeffVar = SD/mean). Line Plots: Lines of same length were drawn across condensates in the LEM2 channel using ImageJ, and the intensity profiles for all channels along the same line measured by the plot profile function. The y-values for each channel were normalized to min=0 and max=1 and plotted in GraphPad Prism 10.

#### Nuclear circularity

An image analysis pipeline was created in CellProfiler to analyze the circularity and eccentricity of the nuclei using LaminA/C staining from the different transduced groups. Individual nuclei were cropped using the DAPI channel > IdentifyPrimaryObjects (in Lamin A/C channel) > MeasureObjectSizeShape.

#### BAF area

For each image, HEK293 cells expressing Tau constructs were identified based on GFP fluorescence. The SNAP-BAF channel was then analyzed to quantify the total BAF positive area per cell. In parallel, the number of discrete BAF foci (dots) per cell was manually counted. For each cell, the BAF area was normalized to the number of BAF foci, and the resulting values were used for statistical analysis and plotting.

### Statistical analysis

For the Statistical analysis GraphPad Prism versions 9 and 10 (GraphPad Software, San Diego, CA, USA; www.graphpad.com) were used. For comparison of two groups, unpaired two-tailed Student’s t-tests were used. For comparison of multiple groups, one-way ANOVAs with Tukey post-test (comparison between all means) or Dunnett post-test (comparison to mean of control) were applied. Significance is indicated as *p<0.05, **p<0.01, ***p<0.001, ****p<0.0001.

### Data Availability

#### Mass spectrometry

Proteomics data have been deposited to the ProteomeXchange Consortium via the PRIDE partner repository with the dataset identifier PXD058565 (Reviewer access details: Username; Password: yPkItu6tXRW5).

#### Sequencing data

The bulkRNAseq and Cut&Tag data are available under the following GEO accession number: GSE283514.

## Contributions

L.D. and S.W. conceived the project, conceptualized the study, curated data, designed figures, and wrote the manuscript.

L.D. and R.P-L. designed and performed experiments and interpreted data.

S.W. conducted FLIM and LLPS experiments and analysis.

C.K. and C.B. helped with PLA experiments.

B.A. developed AAV and TurboID vectors and deigned cloning of cell constructs

T.K. developed experimental protocols and tools for TurboID experiments.

C.W. performed Mass Spectrometry.

K.S. and P.G. performed RNA-Seq data analysis.

K.T. analyzed human array tomography images.

R.A., A.D.-B., M. M., S.H., S.M., B.K.N-H., R.S., M.F. assisted with experiments.

H.R. supplied human brain tissue for PLA and IHC analysis.

T.S.-J. supplied array tomography images from human brain tissue.

E.DeD. and M.B. performed RNA-Seq.

A.N. analyzed CUT&Tag data and consulted in RNA-Seq and CUT&Tag experiments.

T.U. discussed data and reviewed manuscript.

F.L. discussed MS data.

A.v.A. discussed data, reviewed manuscript.

T.T. discussed data and reviewed manuscript.

All authors reviewed and approved the manuscript.

## Acknowledgements

We thank Britta Eickholt, Charité Berlin, for enabling the use of Charité department equipment and general support, Carmelo Ferrai, University Göttingen, for data discussions and advise on chromatin imaging, Gabriele Kaminski Schierle, University Cambridge, for advice on SiR-DNA FLIM, Francsico Garciá-Sierra, Cinvestav Mexico City, for sharing protocols for immunofluorescence in human brain tissue, Donato Di Monte and his team, DZNE Bonn, for sharing protocols for PLA in brain tissue, and Eda Yildrim, Koc University, for chromosome PAINT advise. We thank Thorsten Trimbuch and the Charité viral core facility for AAV vectors production, Bettina Brokowski from Charité viral core facility for help with qPCR experiments, the Jan Schmoranzer and the Charité Bioimaging core (AMBIO) for support in confocal microscopy, Robert Zinzen, MDC BIMSB, for support in FLIM imaging, the single cell sequencing core PRECISE in Bonn, for RNA sequencing and data analysis, the Core Facility Genomics, Berlin Institute of Health at Charité (BIH), for Cut&Tag sequencing, and Max Ruwolt for help with the MS data upload. The authors are most grateful to the individuals and their relatives for consenting to autopsy and subsequent research, which were facilitated by the Biobank of the Department of Neuropathology, Charité-Universitätsmedizin Berlin.

