## Supplemental Figures for "Tau modulates nuclear integrity, chromatin, and cholesterol synthesis genes via the nuclear envelope"

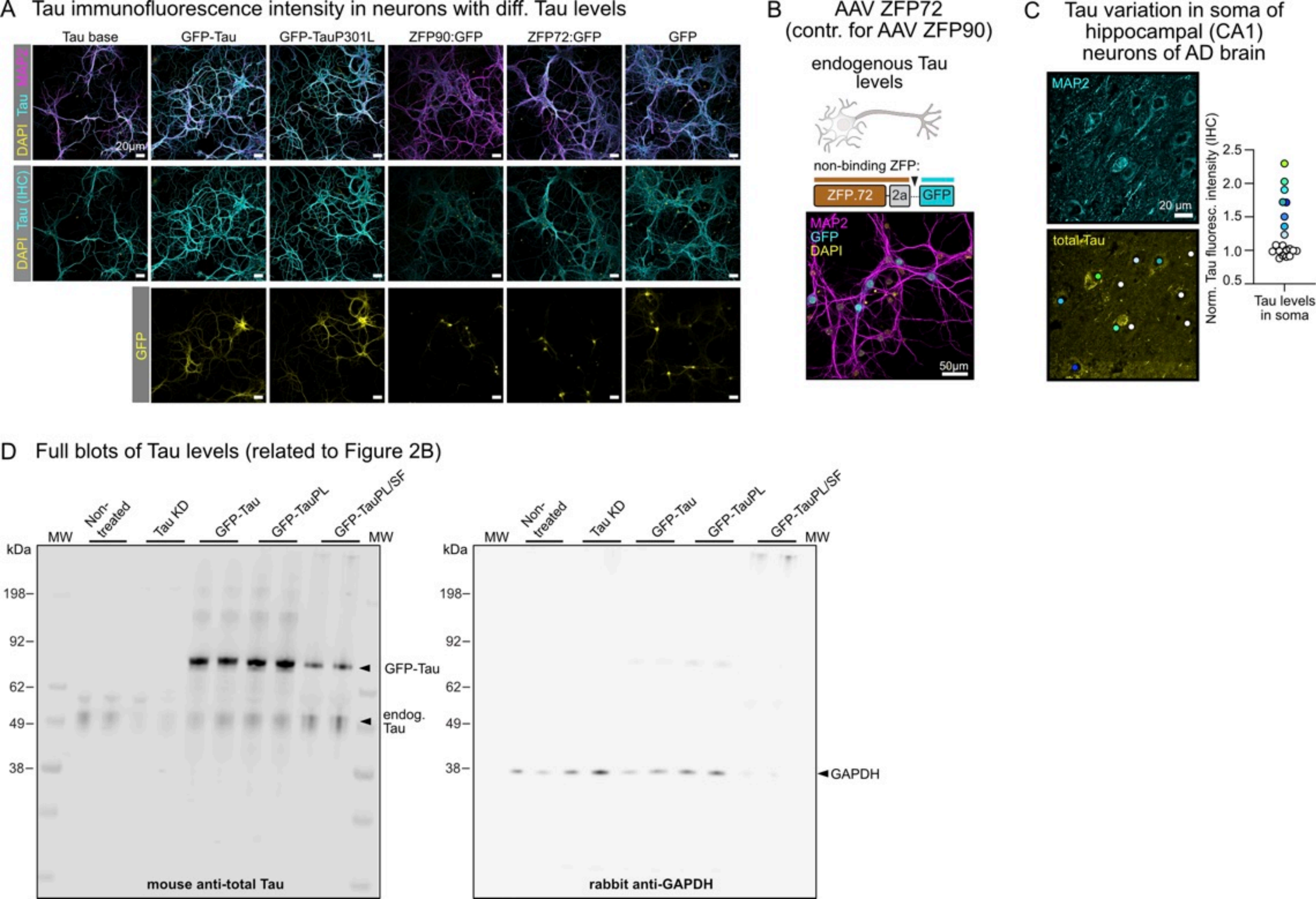
A Tau immunofluorescence intensity in neurons with diff. Tau levels
Tau base
GFP-Tau
GFP-TauP301L
ZFP90:GFP
ZFP72:GFP
GFP
DAPI Tau MAP2
DAPI Tau (IHC)
GFP
20µm
B AAV ZFP72 (contr. for AAV ZFP90)
endogenous Tau levels
non-binding ZFP:
ZFP.72
2a
GFP
MAP2
GFP
DAPI
50µm
C Tau variation in soma of hippocampal (CA1) neurons of AD brain
MAP2
20 µm
total Tau
Norm. Tau fluoresc. intensity (IHC)
2.5
2.0
1.5
1.0
0.5
Tau levels in soma
D Full blots of Tau levels (related to Figure 2B)
MW
Non-treated
Tau KD
GFP-Tau
GFP-TauPL
GFP-TauPL/SF
MW
kDa
198
92
62
49
38
GFP-Tau
endog. Tau
mouse anti-total Tau
GAPDH
rabbit anti-GAPDH

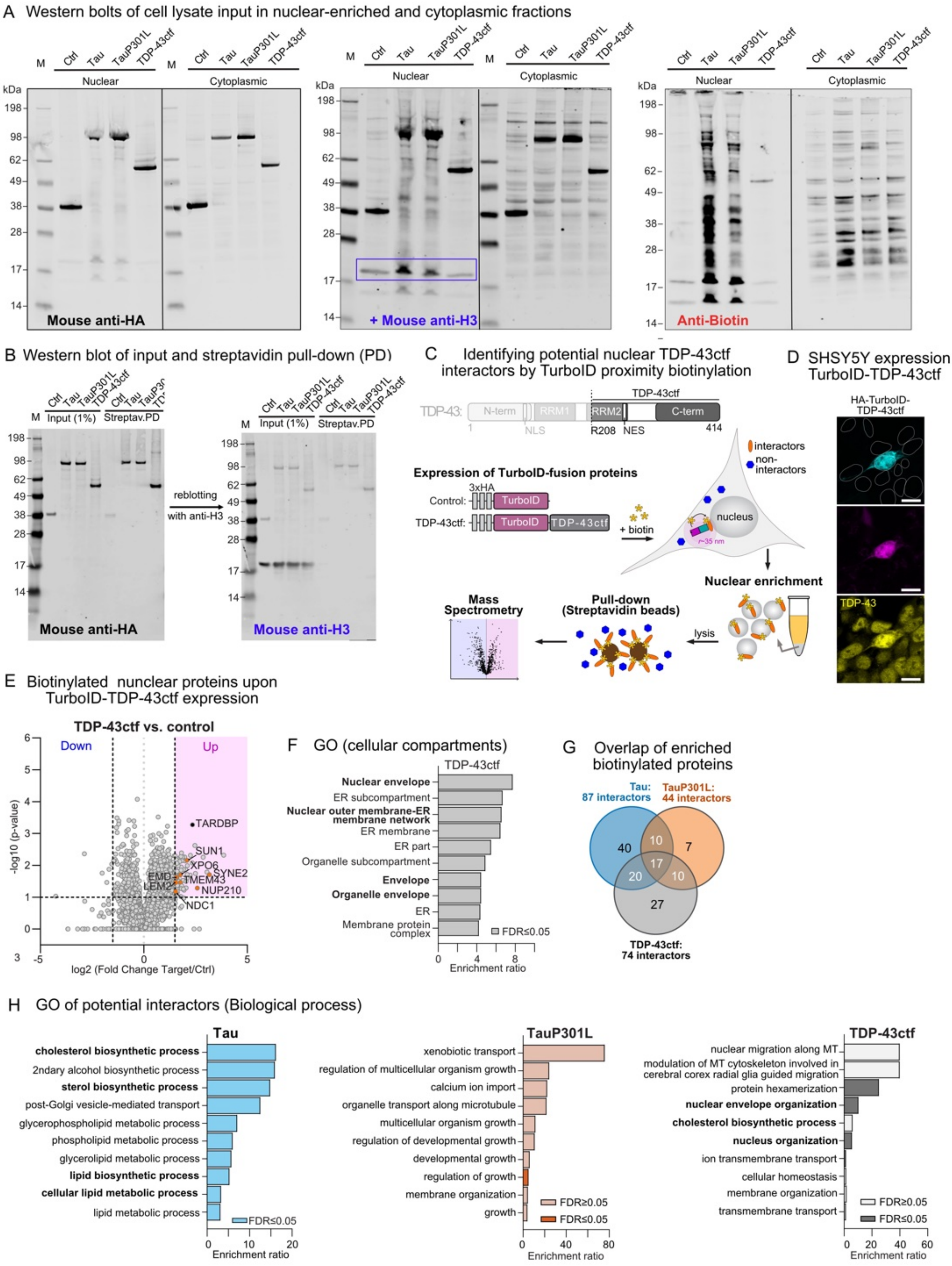
A Western bolts of cell lysate input in nuclear-enriched and cytoplasmic fractions
M
Ctrl
Tau
TauP301L
TDP-43ctf
Nuclear
Cytoplasmic
kDa
198
98
62
49
38
28
17
14
Mouse anti-HA
+ Mouse anti-H3
Anti-Biotin
B Western blot of input and streptavidin pull-down (PD)
Input (1%)
Streptav.PD
reblotting
with anti-H3
Mouse anti-HA
Mouse anti-H3
C Identifying potential nuclear TDP-43ctf interactors by TurboID proximity biotinylation
TDP-43ctf
TDP-43:
N-term
RRM1
RRM2
C-term
1
NLS
R208
NES
414
Expression of TurboID-fusion proteins
3xHA
Control:
TurboID
TDP-43ctf:
TurboID
TDP-43ctf
+ biotin
interactors
non-interactors
nucleus
r~35 nm
Nuclear enrichment
lysis
Pull-down
(Streptavidin beads)
Mass
Spectrometry
D SHSY5Y expression TurboID-TDP-43ctf
HA-TurboID-TDP-43ctf
TDP-43
E Biotinylated nunclear proteins upon TurboID-TDP-43ctf expression
TDP-43ctf vs. control
Down
Up
-log10 (p-value)
log2 (Fold Change Target/Ctrl)
TARDBP
SUN1
XPO6
SYNE2
EMD
TMEM43
LEM2
NUP210
NDC1
F GO (cellular compartments)
TDP-43ctf
Nuclear envelope
ER subcompartment
Nuclear outer membrane-ER membrane network
ER membrane
ER part
Organelle subcompartment
Envelope
Organelle envelope
ER
Membrane protein complex
FDR≤0.05
Enrichment ratio
G Overlap of enriched biotinylated proteins
Tau:
87 interactors
TauP301L:
44 interactors
TDP-43ctf:
74 interactors
40
10
7
17
20
10
27
H GO of potential interactors (Biological process)
Tau
cholesterol biosynthetic process
2ndary alcohol biosynthetic process
sterol biosynthetic process
post-Golgi vesicle-mediated transport
glycerophospholipid metabolic process
phospholipid metabolic process
glycerolipid metabolic process
lipid biosynthetic process
cellular lipid metabolic process
lipid metabolic process
FDR≤0.05
Enrichment ratio
TauP301L
xenobiotic transport
regulation of multicellular organism growth
calcium ion import
organelle transport along microtubule
multicellular organism growth
regulation of developmental growth
developmental growth
regulation of growth
membrane organization
growth
FDR≥0.05
FDR≤0.05
TDP-43ctf
nuclear migration along MT
modulation of MT cytoskeleton involved in cerebral corex radial glia guided migration
protein hexamerization
nuclear envelope organization
cholesterol biosynthetic process
nucleus organization
ion transmembrane transport
cellular homeostasis
membrane organization
transmembrane transport
FDR≥0.05
FDR≤0.05

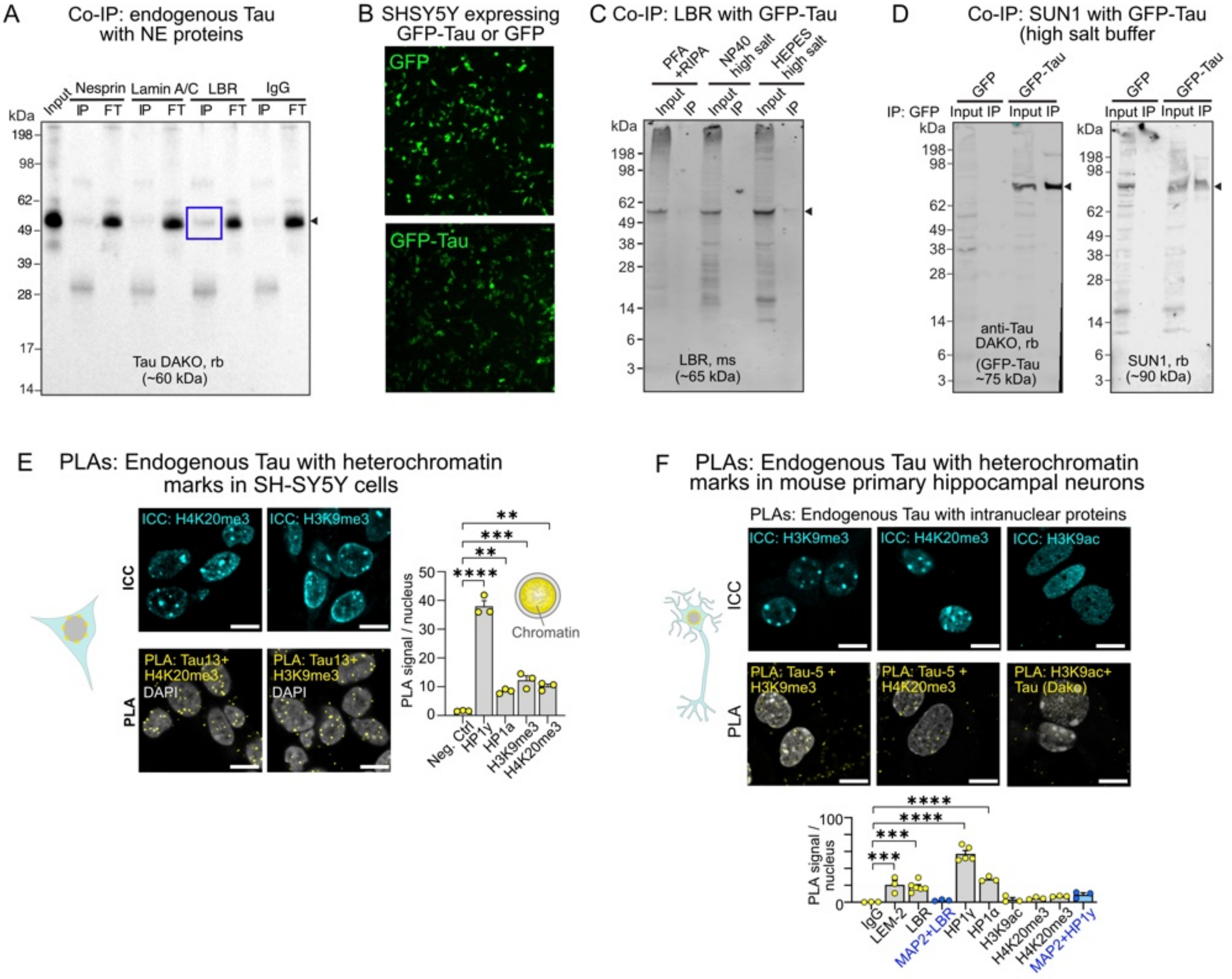

A Co-IP: endogenous Tau with NE proteins
Input
Nesprin
Lamin A/C
LBR
IgG
IP FT
kDa
Tau DAKO, rb (~60 kDa)
B SHSY5Y expressing GFP-Tau or GFP
GFP
GFP-Tau
C Co-IP: LBR with GFP-Tau
PFA +RIPA
NP40 high salt
HEPES high salt
Input IP
LBR, ms (~65 kDa)
D Co-IP: SUN1 with GFP-Tau (high salt buffer)
IP: GFP
GFP
GFP-Tau
anti-Tau DAKO, rb (GFP-Tau ~75 kDa)
SUN1, rb (~90 kDa)
E PLAs: Endogenous Tau with heterochromatin marks in SH-SY5Y cells
ICC: H4K20me3
ICC: H3K9me3
PLA: Tau13+ H4K20me3 DAPI
PLA: Tau13+ H3K9me3 DAPI
PLA signal / nucleus
Chromatin
Neg. Ctrl
HP1γ
HP1α
H3K9me3
H4K20me3
F PLAs: Endogenous Tau with heterochromatin marks in mouse primary hippocampal neurons
PLAs: Endogenous Tau with intranuclear proteins
ICC: H3K9me3
ICC: H4K20me3
ICC: H3K9ac
PLA: Tau-5 + H3K9me3
PLA: Tau-5 + H4K20me3
PLA: H3K9ac+ Tau (Dako)
PLA signal / nucleus
IgG
LEM-2
LBR
MAP2+LBR
HP1γ
HP1α
H3K9ac
H4K20me3
MAP2+HP1γ

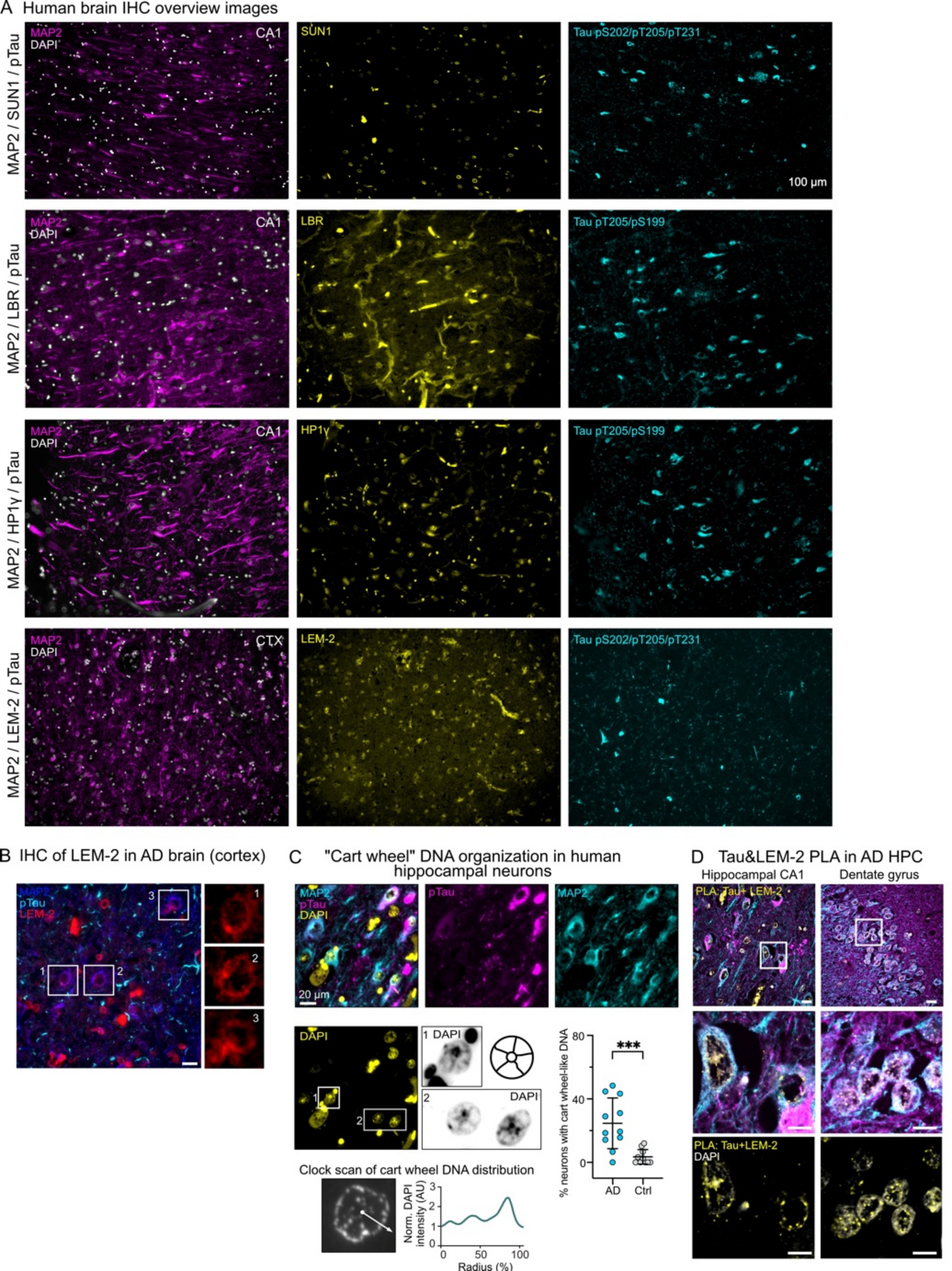
A Human brain IHC overview images
MAP2 / SUN1 / pTau
MAP2
DAPI
CA1
SUN1
Tau pS202/pT205/pT231
100 µm
MAP2 / LBR / pTau
LBR
Tau pT205/pS199
MAP2 / HP1γ / pTau
HP1γ
MAP2 / LEM-2 / pTau
CTX
LEM-2
B IHC of LEM-2 in AD brain (cortex)
pTau
C "Cart wheel" DNA organization in human hippocampal neurons
pTau
20 µm
1 DAPI
2 DAPI
% neurons with cart wheel-like DNA
***
AD
Ctrl
Clock scan of cart wheel DNA distribution
Norm. DAPI intensity (AU)
Radius (%)
D Tau&LEM-2 PLA in AD HPC
Hippocampal CA1
Dentate gyrus
PLA: Tau+ LEM-2
PLA: Tau+LEM-2

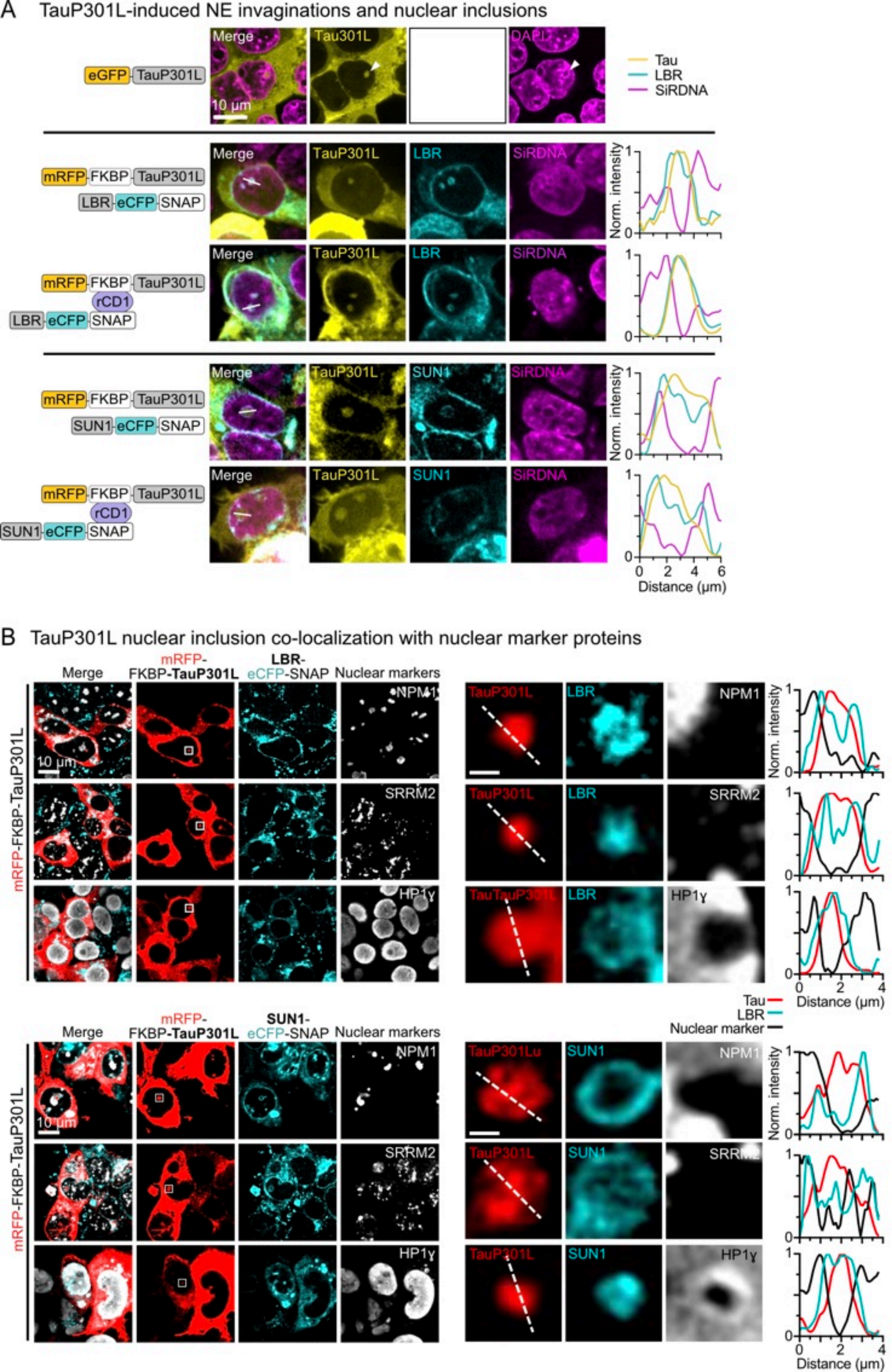
A TauP301L-induced NE invaginations and nuclear inclusions
Merge
Tau301L
DAPI
eGFP
TauP301L
10 µm
Tau
LBR
SiRDNA
mRFP
FKBP
LBR
eCFP
SNAP
rCD1
SUN1
Norm. intensity
Distance (µm)
B TauP301L nuclear inclusion co-localization with nuclear marker proteins
mRFP-FKBP-TauP301L
LBR-eCFP-SNAP
SUN1-eCFP-SNAP
Nuclear markers
NPM1
SRRM2
HP1γ
TauTauP301L
TauP301Lu
Nuclear marker

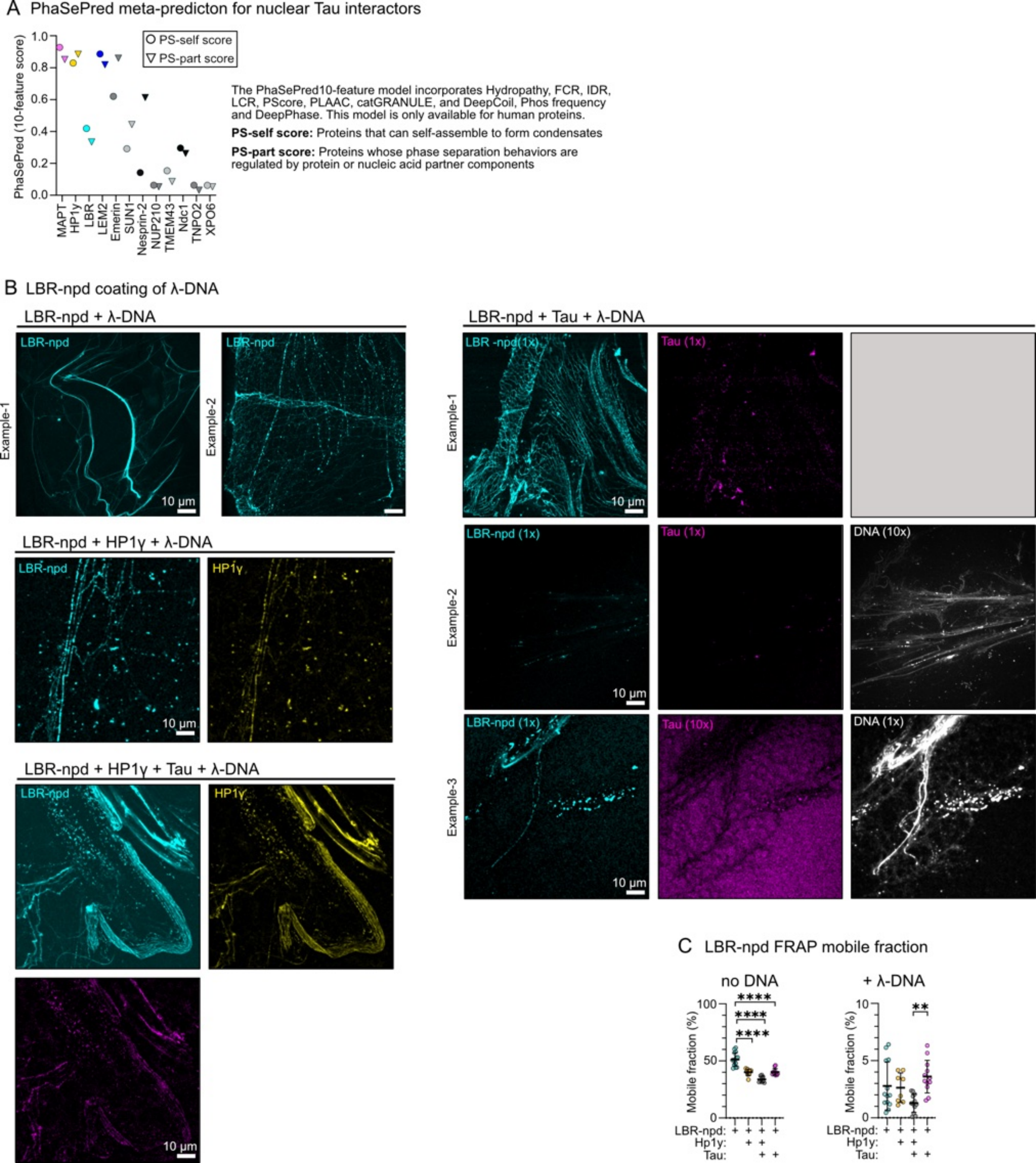
A PhaSePred meta-predicton for nuclear Tau interactors
PhaSePred (10-feature score)
PS-self score
PS-part score
MAPT
HP1γ
LBR
LEM2
Emerin
SUN1
Nesprin-2
NUP210
TMEM43
Ndc1
TNPO2
XPO6
The PhaSePred10-feature model incorporates Hydropathy, FCR, IDR, LCR, PScore, PLAAC, catGRANULE, and DeepCoil, Phos frequency and DeepPhase. This model is only available for human proteins.
PS-self score: Proteins that can self-assemble to form condensates
PS-part score: Proteins whose phase separation behaviors are regulated by protein or nucleic acid partner components
B LBR-npd coating of λ-DNA
LBR-npd + λ-DNA
LBR-npd + Tau + λ-DNA
LBR-npd + HP1γ + λ-DNA
LBR-npd + HP1γ + Tau + λ-DNA
Example-1
Example-2
Example-3
LBR-npd
HP1γ
LBR -npd(1x)
LBR-npd (1x)
Tau (1x)
Tau (10x)
DNA (10x)
DNA (1x)
10 µm
C LBR-npd FRAP mobile fraction
no DNA
+ λ-DNA
Mobile fraction (%)
****
**
LBR-npd:
Hp1γ:
Tau:

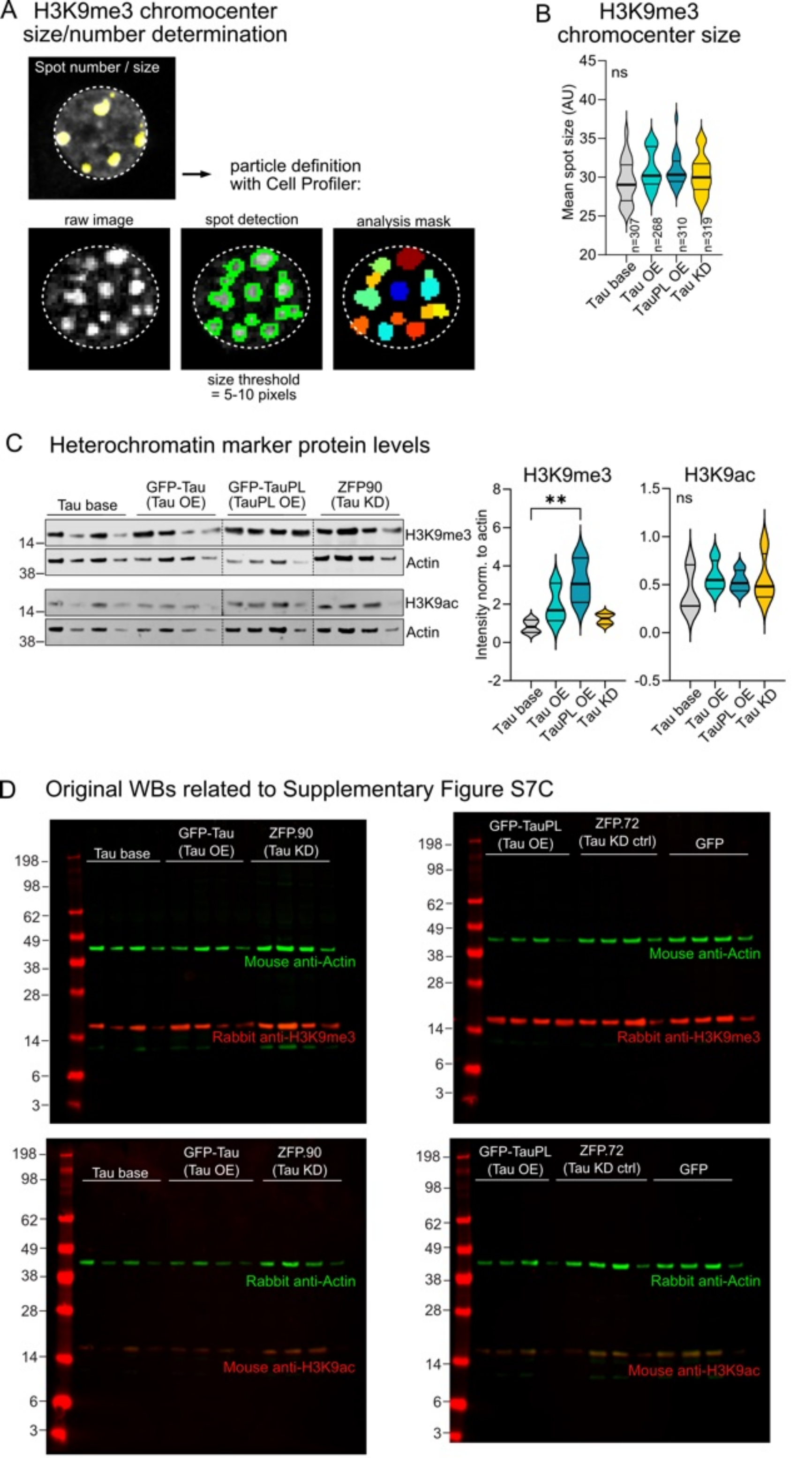
A H3K9me3 chromocenter size/number determination
Spot number / size
particle definition with Cell Profiler:
raw image
spot detection
analysis mask
size threshold = 5-10 pixels
B H3K9me3 chromocenter size
Mean spot size (AU)
ns
n=307
n=268
n=310
n=319
Tau base
Tau OE
TauPL OE
Tau KD
C Heterochromatin marker protein levels
Tau base
GFP-Tau (Tau OE)
GFP-TauPL (TauPL OE)
ZFP90 (Tau KD)
H3K9me3
Actin
H3K9ac
Actin
H3K9me3
H3K9ac
Intensity norm. to actin
**
ns
D Original WBs related to Supplementary Figure S7C
Tau base
GFP-Tau (Tau OE)
ZFP.90 (Tau KD)
GFP-TauPL (Tau OE)
ZFP.72 (Tau KD ctrl)
GFP
Mouse anti-Actin
Rabbit anti-H3K9me3
Rabbit anti-Actin
Mouse anti-H3K9ac

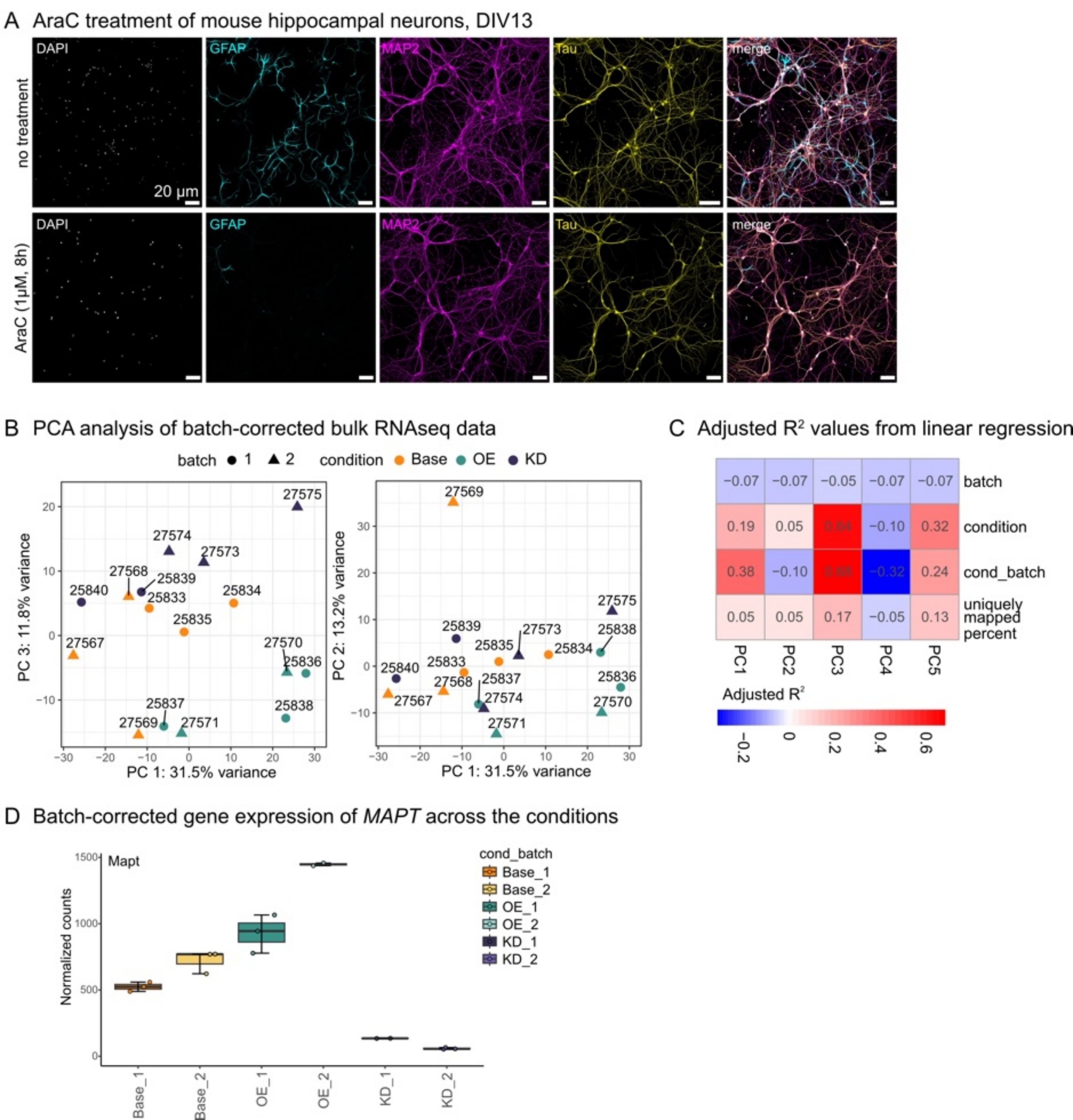
A AraC treatment of mouse hippocampal neurons, DIV13
no treatment
AraC (1µM, 8h)
DAPI
GFAP
MAP2
Tau
merge
20 µm
B PCA analysis of batch-corrected bulk RNAseq data
batch ● 1 ▲ 2 condition ● Base ● OE ● KD
PC 3: 11.8% variance
PC 1: 31.5% variance
PC 2: 13.2% variance
27575
27574
27573
27568
25839
25840
25833
25834
25835
27567
27570
25836
25837
25838
27569
27571
C Adjusted R² values from linear regression
batch
condition
cond_batch
uniquely mapped percent
PC1
PC2
PC3
PC4
PC5
−0.07 −0.07 −0.05 −0.07 −0.07
0.19 0.05 0.64 −0.10 0.32
0.38 −0.10 0.68 −0.32 0.24
0.05 0.05 0.17 −0.05 0.13
Adjusted R²
D Batch-corrected gene expression of MAPT across the conditions
Mapt
Normalized counts
cond_batch
Base_1
Base_2
OE_1
OE_2
KD_1
KD_2

A Number of DEGs by classes

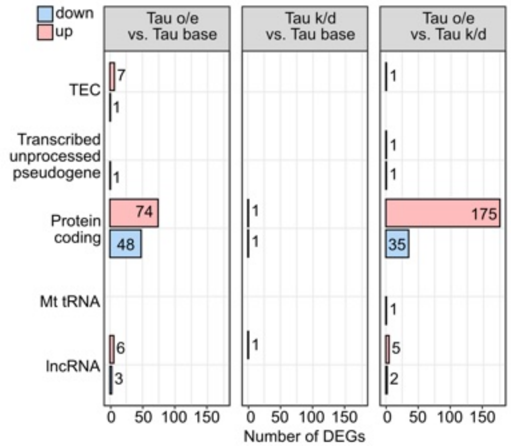


B GO Biological process / all DEGs

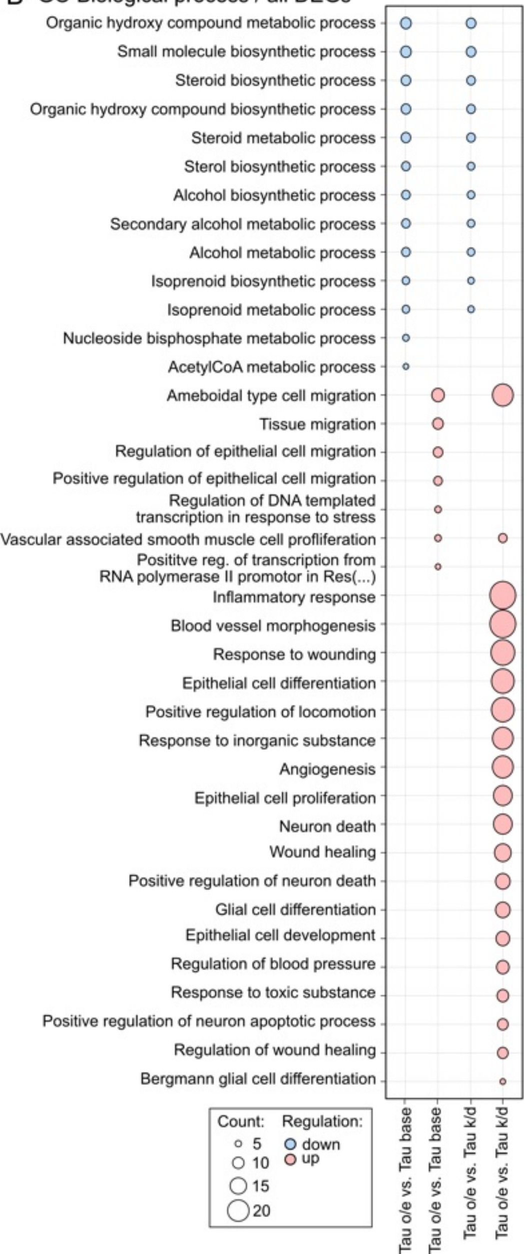


C Non-protein coding DEGs by condition and classes

**Tau o/e vs. Tau base**

| GENEID | Regulation |
|---|---|
| **Protein coding** | |
| n=74 | up |
| **TEC (n=7)** | |
| B230112I24Rik | up |
| Gm37407 | up |
| Gm37959 | up |
| Gm43430 | up |
| Gm45461 | up |
| Gm47075 | up |
| Gm48871 | up |
| **lncRNA (n=6)** | |
| Gm11816 | up |
| Gm34466 | up |
| Gm45346 | up |
| Gm48740 | up |
| Gm50012 | up |
| Neat1 | up |
| **Protein coding** | |
| n=48 | down |
| **TEC (n=1)** | |
| Gm37583 | down |
| **Transcribed unprocessed pseudogene (n=1)** | |
| Gm47135 | down |
| **lncRNA (n=3)** | |
| 1500012K07Rik | down |
| Gm2464 | down |
| Gm48342 | down |

**Tau k/d vs. Tau base**

| GENEID | Regulation |
|---|---|
| **Protein coding** | |
| n=1 | up |
| **lncRNA (n=1)** | |
| Gm42463 | up |
| **Protein coding** | |
| n=1 | down |

**Tau o/e vs. Tau k/d**

| GENEID | Regulation |
|---|---|
| **Protein coding** | |
| n=175 | up |
| **TEC (n=1)** | |
| Gm42928 | up |
| **Transcribed unprocessed pseudogene (n=1)** | |
| Ifi203-ps | up |
| **lncRNA (n=5)** | |
| 1700071M16Rik | up |
| Gm15966 | up |
| Gm49767 | up |
| Neat1 | up |
| Prss23os | up |
| **Protein coding** | |
| n=35 | down |
| **Transcribed unprocessed pseudogene (n=1)** | |
| Gm47135 | down |
| **Mt tRNA (n=1)** | |
| mt-Tf | down |
| **lncRNA (n=2)** | |
| 4921539H07Rik | down |
| Gm48342 | down |

**TEC** (To be Experimentally Confirmed): Regions with EST clusters that have polyA features that could indicate the presence of protein coding genes. These require experimental validation, either by 5' RACE or RT-PCR to extend the transcripts, or by confirming expression of the putatively-encoded peptide with specific antibodies. **lncRNA**: A non-coding gene/transcript >200bp in length. **Unprocessed pseudogene**: Pseudogene that can contain introns since produced by gene duplication. (http://www.ensembl.org/info/genome/genebuild/biotypes.html)

A UniProt keyword enrichment of deregulated TFs in Tau OE vs. Tau base neurons

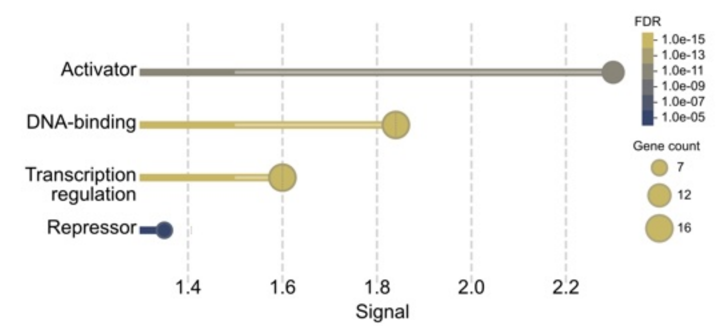


B Deregulated TFs in TauOE vs. Tau base neurons

| TF | Chromosome | Activity | Regulation | log2 [FC] | p-value |
|---|---|---|---|---|---|
| Foxo6 | 4 | Activator | down | -1.826515959 | 5.2851E-08 |
| Neurod6 | 6 | Activator | down | -1.331818147 | 6.18085E-40 |
| Arid5a | 1 | Activator/Repressor | up | 1.989693935 | 4.14874E-19 |
| Arid5b | 10 | Activator | up | 1.501416596 | 1.67707E-14 |
| Atf3 | 1 | Repressor | up | 2.751830962 | 4.2123E-10 |
| Creb5 | 6 | Activator | up | 1.207251673 | 1.27352E-05 |
| Crem | 18 | Activator/Repressor | up | 1.11066829 | 2.93765E-12 |
| Dbx2 | 15 | Repressor | up | 1.137353144 | 3.64843E-05 |
| Ddit3 | 10 | Activator/Repressor | up | 1.18817345 | 3.8227E-12 |
| Hlf | 11 | Activator | up | 1.045046129 | 9.80975E-09 |
| Jun | 4 | Activator | up | 1.238639091 | 2.76742E-20 |
| Nfe2l2 | 2 | Activator | up | 1.01309451 | 1.89093E-14 |
| Nfil3 | 13 | Activator/Repressor | up | 1.540545768 | 7.54893E-10 |
| Rest | 5 | Repressor | up | 1.022649757 | 1.00093E-10 |
| Rorb | 19 | Activator | up | 1.11653135 | 1.73406E-08 |
| Sox9 | 11 | Activator/Repressor | up | 1.097503301 | 4.57973E-07 |
| Tgif1 | 17 | Repressor | up | 1.018888765 | 0.000176675 |

C wPSGA analysis for TFs of DEGs

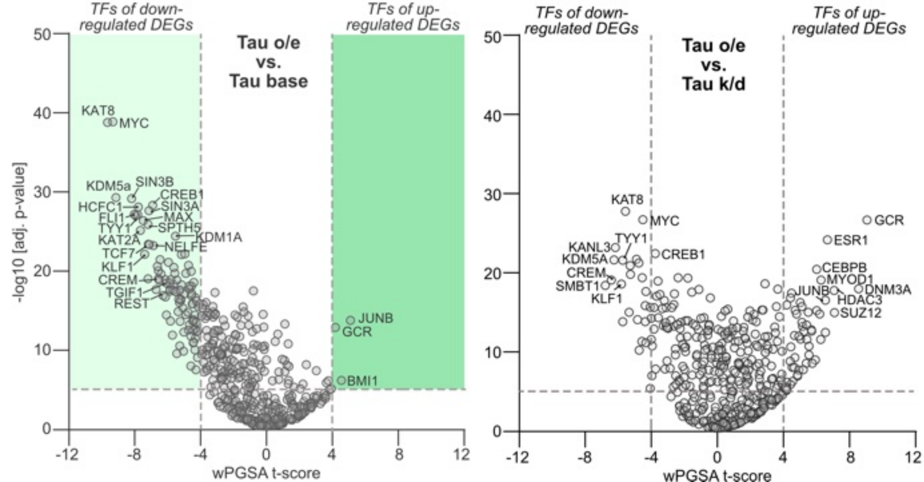

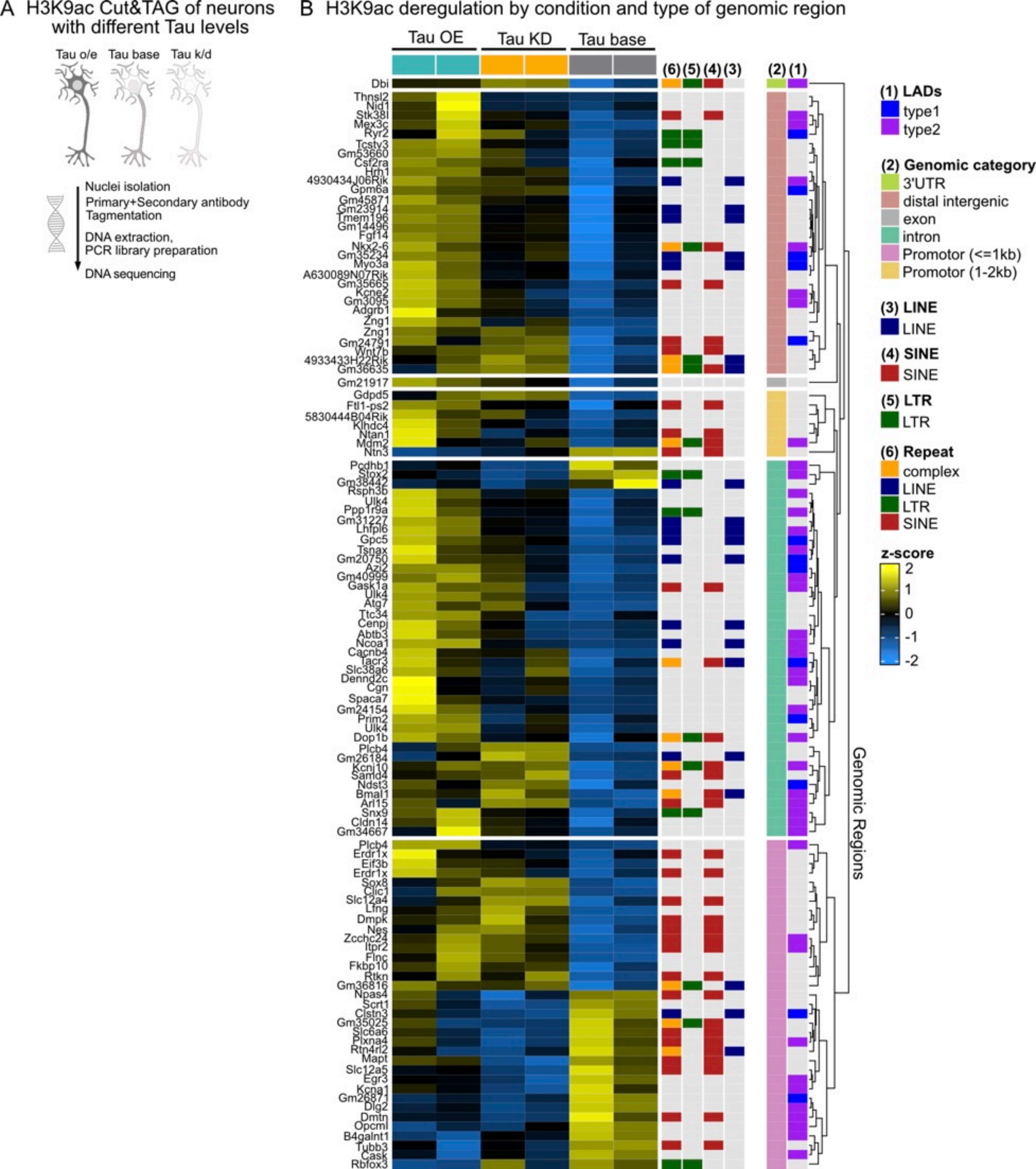
A H3K9ac Cut&TAG of neurons with different Tau levels
Tau o/e
Tau base
Tau k/d
Nuclei isolation
Primary+Secondary antibody
Tagmentation
DNA extraction,
PCR library preparation
DNA sequencing
B H3K9ac deregulation by condition and type of genomic region
Tau OE
Tau KD
Tau base
(6) (5) (4) (3)
(2) (1)
Dbi
Thnsl2
Nid1
Stk38l
Mex3c
Ryr2
Tcstv3
Gm53660
Csf2ra
Hrh1
4930434J06Rik
Gpm6a
Gm45871
Gm23914
Tmem196
Gm14496
Fgf14
Nkx2-6
Gm35234
Myo3a
A630089N07Rik
Gm35665
Kcne2
Gm3095
Adgrb1
Zng1
Zng1
Gm24791
Wnt7b
4933433H22Rik
Gm36635
Gm21917
Gdpd5
Ftl1-ps2
5830444B04Rik
Klhdc4
Ntan1
Mdm2
Ntn3
Pcdhb1
Stox2
Gm38442
Rsph3b
Ulk4
Ppp1r9a
Gm31227
Lhfpl6
Gpc5
Tsnax
Gm20750
Azi2
Gm40999
Gask1a
Ulk4
Atg7
Ttc34
Cenpj
Abtb3
Ncoa1
Cacnb4
Tacr3
Slc38a6
Dennd2c
Cgn
Spaca7
Gm24154
Prim2
Ulk4
Dop1b
Plcb4
Gm26184
Kcnj10
Samd4
Ndst3
Bmal1
Arl15
Snx9
Cldn14
Gm34667
Plcb4
Erdr1x
Eif3b
Erdr1x
Sox8
Clic1
Slc12a4
Lfng
Dmpk
Nes
Zcchc24
Itpr2
Flnc
Fkbp10
Rtkn
Gm36816
Npas4
Scrt1
Clstn3
Gm35025
Slc6a6
Plxna4
Rtn4rl2
Mapt
Slc12a5
Egr3
Kcna1
Gm26871
Dlg2
Dmtn
Opcml
B4galnt1
Tubb3
Cask
Rbfox3
Genomic Regions
(1) LADs
type1
type2
(2) Genomic category
3'UTR
distal intergenic
exon
intron
Promotor (<=1kb)
Promotor (1-2kb)
(3) LINE
LINE
(4) SINE
SINE
(5) LTR
LTR
(6) Repeat
complex
LINE
LTR
SINE
z-score
2
1
0
-1
-2

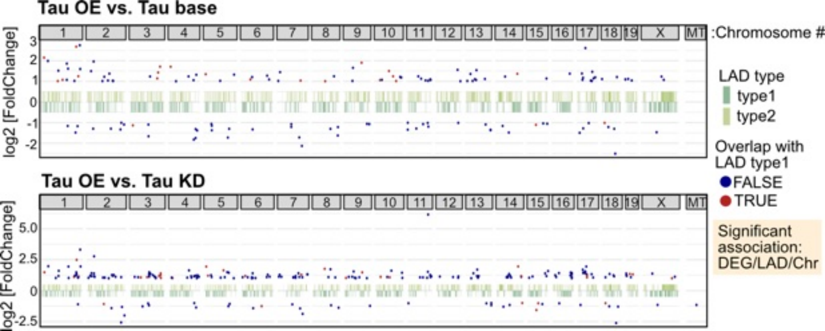
Tau OE vs. Tau base
Tau OE vs. Tau KD
1 2 3 4 5 6 7 8 9 10 11 12 13 14 15 16 17 18 19 X MT
:Chromosome #
log2 [FoldChange]
3
2
1
0
-1
-2
5.0
2.5
0
-2.5
LAD type
type1
type2
Overlap with LAD type1
FALSE
TRUE
Significant association: DEG/LAD/Chr
